# The incentive circuit: memory dynamics in the mushroom body of *Drosophila melanogaster*

**DOI:** 10.1101/2021.06.11.448104

**Authors:** Evripidis Gkanias, Li Yan McCurdy, Michael N Nitabach, Barbara Webb

**Affiliations:** Institute of Perception Action and Behaviour, School of Informatics, University of Edinburgh, UK; Department of Cellular and Molecular Physiology, Yale University, New Haven, CT, USA; Department of Genetics, Yale University, New Haven, CT, USA; Department of Neuroscience, Yale University, New Haven, CT, USA

## Abstract

Insects adapt their response to stimuli, such as odours, according to their pairing with positive or negative reinforcements, such as sugar or shock. Recent electrophysiological and imaging findings in *Drosophila melanogaster* allow detailed examination of the neural mechanisms supporting the acquisition, forgetting and assimilation of memories. We propose that this data can be explained by the combination of a dopaminergic plasticity rule that supports a variety of synaptic strength change phenomena, and a circuit structure (derived from neuroanatomy) between dopaminergic and output neurons that creates different roles for specific neurons. Computational modelling shows that this circuit allows for rapid memory acquisition, transfer from short-term to long-term, and exploration/exploitation trade-off. The model can reproduce the observed changes in the activity of each of the identified neurons in conditioning paradigms and can be used for flexible behavioural control.

## Introduction

Animals deal with a complicated and changing world and they need to adapt their behaviour according to their recent experience. Rapid changes in behaviour to stimuli accompanied by intense reinforcement requires memories in the brain that are readily susceptible to alteration. Yet associations experienced consistently should form long-term memories that are hard to change. Memories that are no longer valid should be forgotten. Every neuron cannot have all of these properties, but they must be connected in a circuit, playing different roles such as supporting short- or long-term memory, and enabling processes to form, retain and erase memories. This complex interaction of memory processes is familiar in principle but its implementation at the single neuron level is still largely a mystery.

The fruit fly *Drosophila melanogaster* is able to form, retain and forget olfactory associations with reinforcements, e.g., electric shock. The key neural substrate is known to lie in neuropils of their brain called the *mushroom bodies* (MB) (***Davis, 1993***; ***Heisenberg, 2003***; ***Busto et al., 2010***). There are two MBs in the insect brain, one in each hemisphere, composed of intrinsic and extrinsic neurons. Extrinsic *projection neurons* (PN) deliver sensory input to the only intrinsic neurons of the mushroom bodies, the *Kenyon cells* (KC), whose long parallel axons travel through the pendunculus and then split forming the vertical (α/α’) and medial (β/β’ and γ) mushroom body lobes (see ***Figure 1***). The extrinsic *mushroom body output neurons* (MBON) extend their dendrites in different regions of the lobes, receiving input from the KCs and forming 15 distinct compartments (***Turner et al., 2008***; ***Tanaka et al., 2008***; ***Campbell et al., 2013***; ***Aso et al., 2014a***). Their activity is thought to provide motivational output that modulates the default behaviour of the animal (***Aso et al., 2014b***). Different groups of extrinsic *dopaminergic neurons* (DAN) terminate their axons in specific compartments of the MB, and modulate the connections between KCs and MBONs (***Aso et al., 2014a***). Many of the DANs respond to a variety of reinforcement signals (***Mao and Davis, 2009***; ***Schwaerzel et al., 2003***; ***Claridge-Chang et al., 2009***; ***Liu et al., 2012***; ***Lin et al., 2014***) and therefore they are considered the main source of reinforcement signals in the mushroom body. Finally, many of the MBON axons and DAN dendrites meet in the convergence zones (CZ), where they create interconnections, such that the motivational output can also influence the activity of the reinforcement neurons (***Li et al., 2020***).

**Figure 1.**
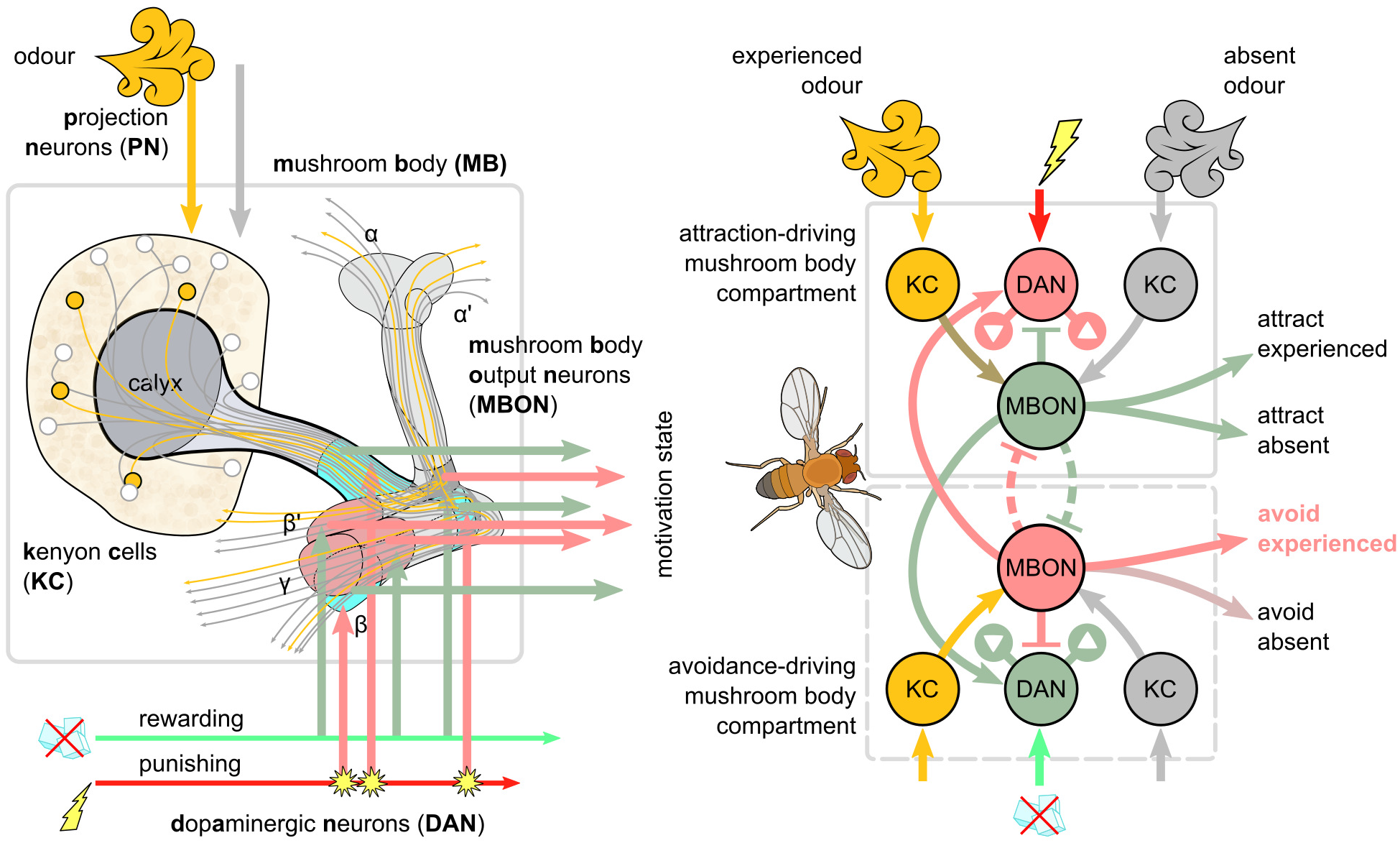
Overview of the mushroom body circuit. Left: the main anatomical pathways. In the illustration, the presented odour activates the Kenyon cells (KCs) through the projection neurons (PNs). The parallel axons of KCs propagate this signal to the lobes of the mushroom body. The mushroom body output neurons extend their dendrites in the mushroom body lobes, receiving input from the KCs. Electric shock creates a punishing signal that excites some dopaminergic neurons (DANs), whose axons terminate in the lobes and modulate the synaptic weights between KCs and MBONs. Right: schematic of potential connections between punishment/reward DANs and approach/avoidance MBONs. Note that although DANs transferring punishing signals modulate the KC activation of MBONs that encode positive motivations [decreasing attraction to the presented odour and increasing attraction to odours not present (The dopaminergic plasticity rule)], MBONs that encode negative motivations will also gain higher responses due to release of inhibition between MBONs, and the feedback connections from MBONs to other DANs. In our model, we further decompose these functions using three DANs and three MBONs for each motivation (positive or negative), and map these units to specific identified neurons and microcircuits in the brain of Drosophila. These circuits include some direct (but not mutual) MBON-MBON connections (dashed inhibitory connections).

Several computational models have tried to capture the structure and function of the mushroom bodies, usually abstracting the common features of this network across various insect species. Modellers have treated the mushroom bodies as performing odour discrimination (***Huerta et al., 2004***), olfactory conditioning (***Balkenius et al., 2006***; ***Smith et al., 2008***; ***Finelli et al., 2008***; ***Young et al., 2011***; ***Wessnitzer et al., 2012***; ***Peng and Chittka, 2017***; ***Faghihi et al., 2017***; ***Zhao et al., 2021***; ***Springer and Nawrot, 2021***; ***Eschbach et al., 2020***; ***Bennett et al., 2021***) or calculating the scene familiarity (***Wu and Guo, 2011***; ***Baddeley et al., 2012***; ***Arena et al., 2013***; ***Bazhenov et al., 2013***; ***Ardin et al., 2016***). However, it seems like they can subserve all these functions, depending on context (or experience), i.e., what is driving the activity of the KCs (***Cohn et al., 2015***). This suggests that the output neurons of the mushroom body do not just inform the animal whether an odour is known or attractive, or if a scene is familiar, but they actually motivate the animal to take an action like approach, avoid, escape or forage. There is emerging evidence supporting this idea of the MBONs driving non-binary but antagonistic motivations (***Schwaerzel et al., 2003***; ***Krashes et al., 2009***; ***Gerber et al., 2009***; ***Waddell, 2010***; ***Lin et al., 2014***; ***Perisse et al., 2016***; ***Senapati et al., 2019***), which has started to be explored in recent models.

In addition to the structural and functional depiction of the mushroom bodies, a variety of plasticity rules have been used in order to explain the effect of dopamine emitted by the DANs on the KC→MBON synapses. Although the best supported biological mechanism is that coincidence of DAN and KC activity depresses the output of KCs to MBONs, most of the models mentioned before use variations of the Hebbian rule (***Hebb, 2005***), where the coincidence of the input (KCs) and output (MBONs) activation strengthens the synaptic weight (or weakens itforthe anti-Hebbian case) and this is gated by the reinforcement (DANs). More recent approaches that try to model the activity of DANs and MBONs in the brain have used plasticity rules (***Zhao et al., 2021***) or circuit structures (***Springer and Nawrot, 2021***; ***Bennett et al., 2021***; ***Eschbach et al., 2020***) that implement a reward prediction error (RPE) (***Rescorla and Wagner, 1972***), which is the most widely accepted psychological account of associative learning with strong validation evidence in the vertebrate brain (***Niv, 2009***). For the mushroom body, this plasticity rule is interpreted as the output (MBON) being the prediction of the reinforcement (DAN), so their difference (gated by the activity of the input, KC) drives the synaptic plasticity. However, details of neuronal dynamics in fruit flies (***Hige et al., 2015***; ***Dylla et al., 2017***; ***Berry et al., 2018***) suggest that neither Hebbian or RPE plasticity rules capture the plasticity dynamics in the mushroom bodies [also in larva (***Schleyer et al., 2018***, ***2020***)], as both rules need the CS to occur (KCs to be active) for synaptic weight change. This highlights the importance of investigating new plasticity rules that are a closer approximation to the actual dopaminergic function.

In this work, we propose such a novel plasticity rule, named the *dopaminergic plasticity rule* (DPR), which reflects recent understanding of the role of dopamine in depression and potentiation of synapses. Based on evidence of existing MBON→DAN connections, we build a 12-neuron computational model, that we call the *incentive circuit* (IC), which uses the proposed plasticity rule. In this model, we name 3 types of DANs (‘discharging, ‘charging and ‘forgetting’) and 3 types of MBONs (‘susceptible’, ‘restrained’ and ‘long-term memory’) for each of two opposing motivational states. We demonstrate that the neural responses generated by this model during an aversive olfactory learning paradigm replicate those observed in the animal; and that simulated flies equipped with the incentive circuit generate learned odour preferences comparable to real flies. Finally, we suggest that such a model could work as motif that extends the set of motivations from binary (e.g., avoidance vs attraction) to a spectrum of motivations whose capabilities are equivalent to ‘decision making’ in mammals.

## Results

### The dopaminergic plasticity rule

We implement a novel *dopaminergic plasticity rule* (DPR) to update the KC→MBON synaptic weights proportionally to the dopamine level, KC activity and current state of the synaptic weight with respect to its default (rest) value. Our DPR is based on recent findings regarding the role of dopamine (and co-transmitters) in altering synaptic efficacy in the fruit fly mushroom body (see Derivation of the dopaminergic plasticity rule). Instead of calculating the error between the reinforcement and its prediction, DPR uses the reinforcement to maximise the separation between the synaptic weights of reinforced inputs, which is functionally closer to the information maximisation theory (***Bell and Sejnowski, 1995***; ***Lee et al., 1999***; ***Lulham et al., 2011***) than the RPE principle. While this rule, in combination with some specific types of circuits, can result in prediction of reinforcements, it can also support a more flexible range of responses to stimulus-reinforcement contingencies, as we will show in what follows.

The dopaminergic learning rule is written formally as

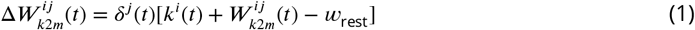

where 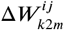 is the change in the synaptic weight connecting a KC, *i*, to an MBON, *j*. The KC→MBON synaptic weight, 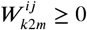, and the KC response, *k^i^*(*t*) ≥ 0, have a lower-bound of 0, while the resting weight, *w*_rest_ = 1, is a fixed parameter. The rule alters the connection weight between each KC and MBON on each time step depending on the *dopaminergic factor*, *δ^j^*(*t*), which is determined by the responses of the DANs targeting this MBON. The dopaminergic factor can be positive [*δ^j^*(*t*) > 0] or negative [*δ^j^*(*t*) < 0], which we motivate from recent observations of the differential roles in synaptic plasticity of DopR1 and DopR2 receptors (***Handler et al., 2019***), as detailed in the methods. When combined with two possible states of KC activity (active or inactive) this results in four different plasticity effects: *depression, potentiation, recovery* and *saturation*.

These effects can be inferred directly from ***Equation 1***. If the dopaminergic factor is zero (contributing DANs are inactive or mutually cancelling), no learning occurs. If the dopaminergic factor is negative and the KC is active (positive), the KC→MBON synaptic weight is decreased (*depression* effect of the plasticity rule, see ***Figure 2***A). However, if the synaptic weight is already low, the synaptic weight cannot change further. The *recovery* effect takes place when the dopaminergic factor is negative and the KC is inactive (*k^i^*(*t*) = 0), in which case the synaptic weights tend to reset to the resting weight (see ***Figure 2***C). When the dopaminergic factor is positive and the KC is active, we have the potentiation effect, which causes increase in the synaptic weights (see ***Figure 2***B). In contrast to the depression effect, as the synaptic weight becomes stronger, it further enhances this effect. If the KC is inactive, and the dopaminergic factor is positive then we have the *saturation* effect, where if the current synaptic weight is higher than its resting weight, the synaptic weight continues increasing, while if it is lower then it continues decreasing (see ***Figure 2***D). This effect enhances diversity in the responses of the MBON to the different past and current CS experiences, which is essential for memory consolidation (i.e., continued strengthening of a memory) and for the formation of long-term memories (i.e., slower acquisition and resistance to further change).

**Figure 2.**
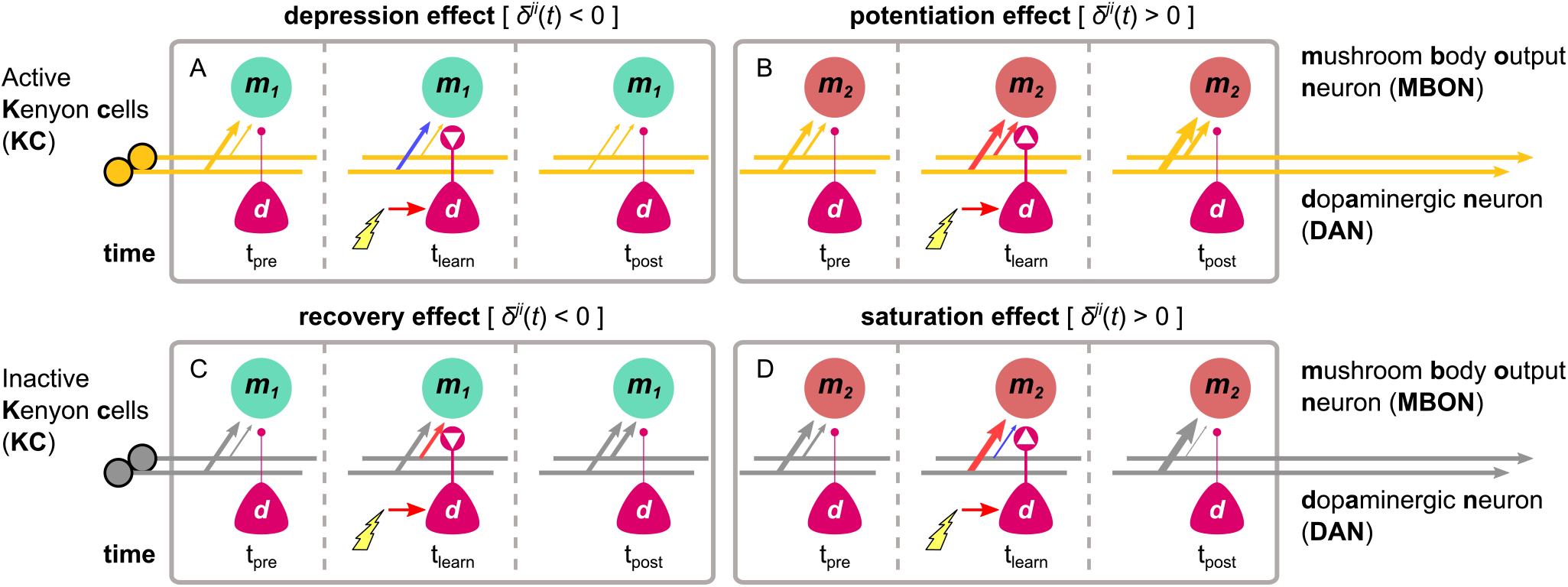
The different effects of the dopaminergic plasticity rule, depending on the activity of the KC (orange indicates active) and the sign of the dopaminergic factor (white arrowheads in dots). The DPR can cause 4 different effects that work in harmony or discord to maximise the information captured in each experience and allow different types of memories to be formed for each KC→MBON synapse. In each box, time-step *t* = *t*_pre_ shows the initial KC→MBON synaptic weights (thickness of the arrows); electric shock activates the DAN in time-step *t* = *t*_learn_ causing modulation of the synaptic weights (red: increase, blue: decrease), while time-step *t* = *t*_post_ shows the synaptic weights after the shock delivery. (A) Example of the *depression effect* - the synaptic weight decreases when *δ^j^*(*t*) < 0 and the KC is active. (B) Example of the *potentiation effect* - the synaptic weight increases when *δ^j^*(*t*) > 0 and the KC is active. (C) Example of the *recovery effect* - the synaptic weight increases when *δ^j^*(*t*) < 0 and the KC is inactive. (D) Example of the *saturation effect* - the synaptic weight increases further (when 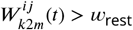) or decreases further (when 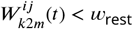) when *δ^j^*(*t*) > 0 and the KC is inactive.

The different effects described above can work together in single KC→MBON synapses (i.e., through influence of multiple DANs) leading in more complicated effects like the formation of short-term memories (e.g., combining the depression/potentiation and recovery effects) or long-term memories (e.g., combining the potentiation and saturation effects). However, we will see that by adding MBON→DAN feedback connections a very wide range of circuit properties can be implemented. We next introduce a set of microcircuits that have been found in the fruit fly mushroom bodies and describe how they could interlock and interact in one ‘incentive circuit’ to control the motivation and hence the behaviour of the animal.

### The incentive circuit

What we call the incentive circuit (IC) is a circuit in the mushroom body of the fruit fly *Drosophila melanogaster* that allows complicated memory dynamics through self-motivation. We have identified and modelled this circuit (shown in ***Figure 3***) which consists of six MBONs that receive KC input, and six DANs that modulate the KC-MBON connections. The circuit includes some MBON-MBON connections and some feedback connections from MBONs to DANs. All the neurons and connections in this circuit are mapped to identified connectivity in the mushroom body as summarised in ***Table 1***. We will describe each of the microcircuits and the biological justification for their assumed function in detail below, but here provide an initial overview of the incentive circuit function.

**Figure 3.**
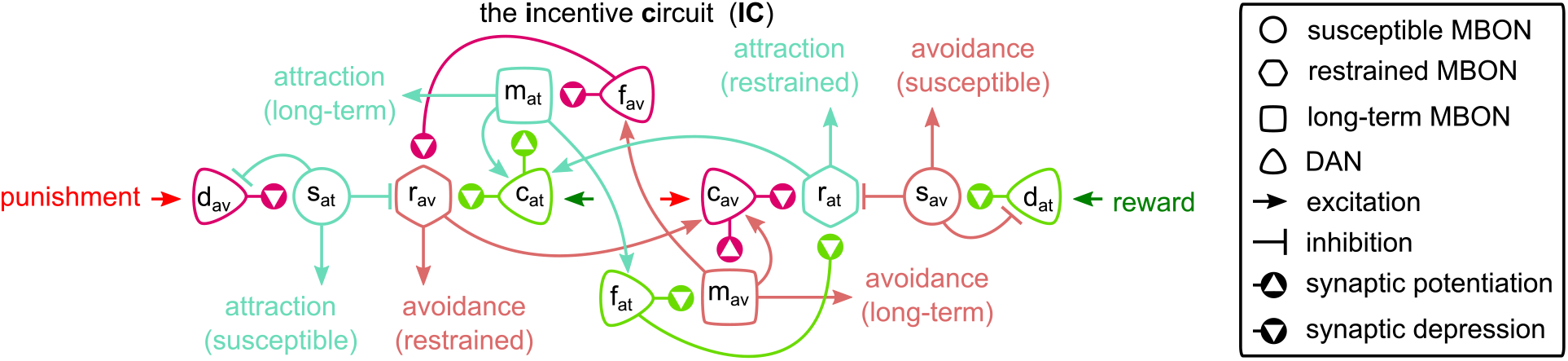
The *incentive circuit* (IC) integrates the different microcircuits of the mushroom body into a unifled model allowing the expression of more complicated behaviours and memory dynamics. It combines the susceptible, restrained, reciprocal short- and long-term memories and the memory assimilation mechanism microcircuits in one circuit that is able to form, consolidate and forget different types of memories that motivate the animal to take actions. *d*_av_ and *d*_at_: avoidance- and attraction-driving discharging DANs. *c*_av_ and *c*_at_: avoidance- and attraction-driving charging DANs. *f*_av_ and *f*_at_: avoidance- and attraction-driving forgetting DANs. *s*_av_ and *s*_at_: avoidance- and attraction-driving susceptible MBONs. *r*_av_ and *r*_at_: avoidance- and attraction-driving restrained MBONs. *m*_av_ and *m*_at_: avoidance- and attraction-driving long-term memory MBONs. **Figure 3–Figure supplement 1.** Behaviour generated by the model correlates with 92 experiments reported in ***Bennett et al.*** (***2021***) — ***Figure 5***. **Figure 3–source data 1.** Experimental data from ***Bennett et al.*** (***2021***) modified to include the predicted neuron types.

**Table 1.**
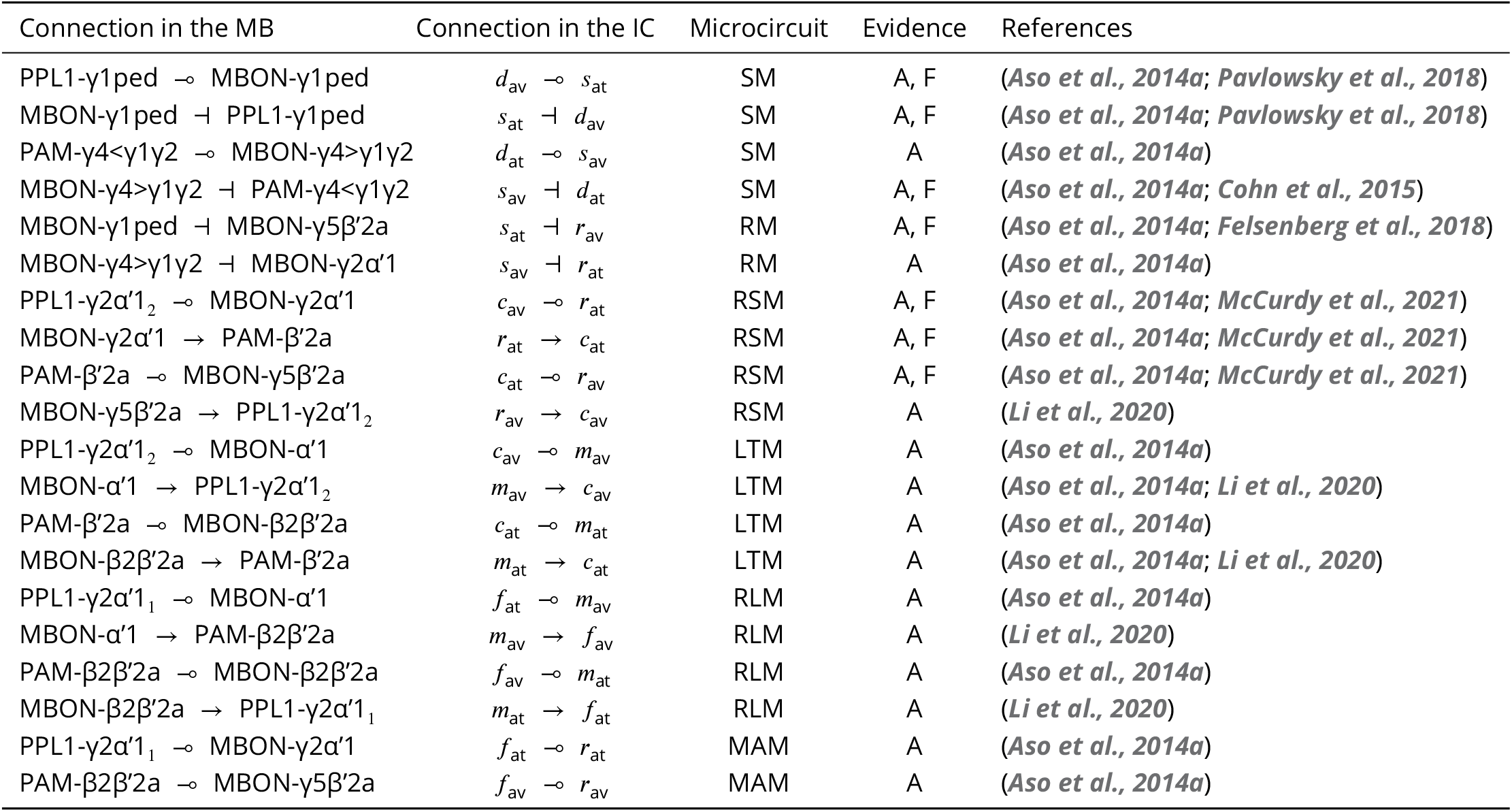
Connections among neurons in the Drosophila mushroom body mapped to the connections of the incentive circuit. Connection types: ‘⊸’, modulates the synaptic weights of the KC→MBON connections terminating in that MBON; ‘→’, excitatory connection; ‘⊣’, inhibitory connection. Microcircuit — SM: susceptible memory, RM: restrained memory, RSM: reciprocal short-term memories, LTM: long-term memory, RLM: reciprocal long-term memories, MAM: memory assimilation mechanism. Evidence — A: anatomical connection is known (i.e., using light or electron microscopy), F: functional connection is known (i.e., whether activating the presynaptic neuron leads to an excitatory or inhibitory effect on the postsynaptic neuron, and/or the neurotransmitter released by the presynaptic neuron).

As presented in ***Figure 3***, for each motivation (attraction or avoidance), the incentive circuit has three types of MBON — susceptible, restrained, and long-term memory — and three types of DAN — discharging, charging and forgetting. More specifically, working from the outer edges of the model, we have ‘discharging DANs that respond to punishment (left side) or reward (right side) and influence the ‘susceptible’ MBONs which by default respond to all KC inputs (not shown). These in turn inhibit the responses of the ‘restrained’ MBONs of opposite valence. When the discharging DANs depress the response of the susceptible MBONs of opposite valence, they release the restrained MBONs of the same valence, and also decrease the inhibitory feedback to the discharging DANs from the susceptible MBONs. The restrained MBONs activate their respective ‘charging’ DANs, which start to potentiate the ‘LTM’ MBONs of same valence, while also depressing the response (to KC input) of the restrained MBON of opposite valence. Similarly, the LTM MBONs enhance the activity of the charging DANs, increasing the momentum of LTM, while simultaneously activating their respective ‘forgetting DANs, to decrease momentum of the opposite valence LTM. The forgetting DANs also depress the restrained MBONs which makes space for acquisition of new memories while preserving old ones.

In the following sections we show in detail how each simulated neuron of this circuit responds during acquisition and forgetting in the aversive olfactory conditioning paradigm shown in ***Figure 4***, and compare this to observed responses in the corresponding identified neurons in the fly from calcium imaging under the same paradigm. We then describe the behaviour of simulated flies under the control of this circuit and learning rule in a naturalistic setting with two odour gradients, paired singly or jointly with punishment or reward. By using more abstracted behavioural modelling, following the approach of ***Bennett et al.*** (***2021***), we are also able to create closely matching results for 92 different olfactory conditioning intervention experiments, i.e., the observed effects on fly learning of silencing or activating specific neurons (***Figure 3***–***Figure Supplement 1***, Δ*f* of the model and experiments are correlated with correlation coefficient *r* = 0.76, *p* = 2.2 × 10^−18^).

**Figure 4.**
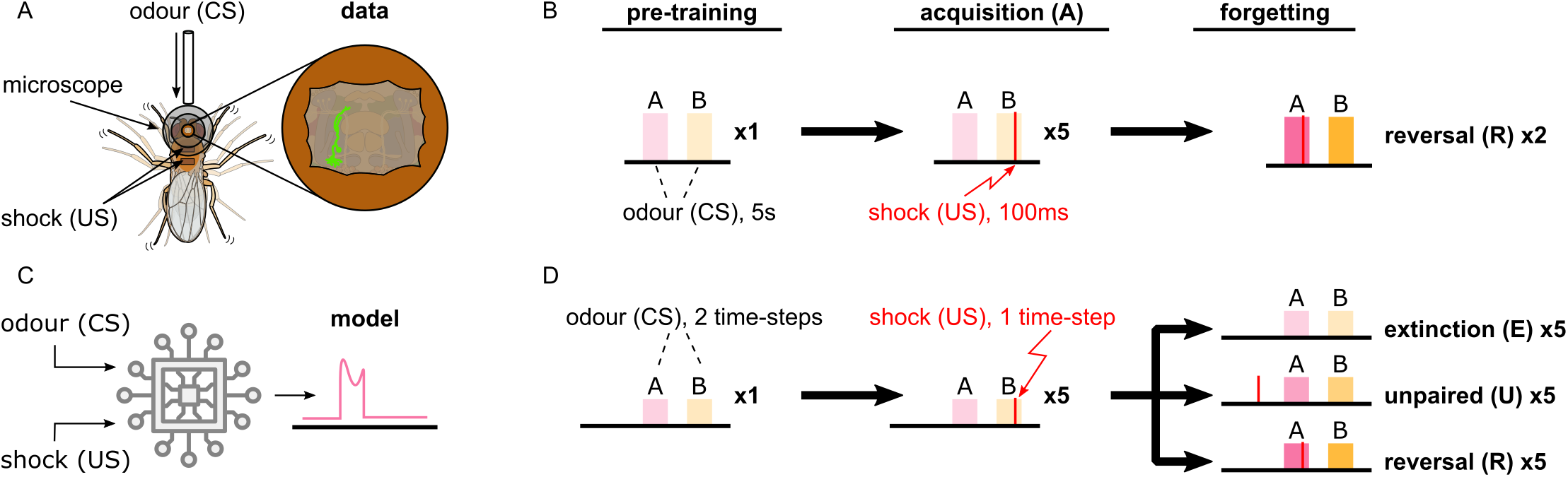
Description of the experimental setup and the aversive olfactory conditioning paradigms. (A) Setup for visualising neural activity via Ca^2+^imaging during aversive olfactory memory acquisition and reversal. Flies are head-fixed and cuticle dissected for ratiometric imaging of Ca^2+^-sensitive GCaMP_6_f and Ca^2+^-insensitive tdTomato. (B) The aversive olfactory conditioning experimental paradigm. 5 sec presentations of odours A (3-octanol — OCT; coloured pink) and B (4-methylcyclohexanol — MCH; coloured yellow) continuously alternate, separated by fresh air, while the shock input (100 ms of 120 V) forms the different phases: 1 repeat of pre-training, where no shock is delivered; 5 repeats of acquisition, where shock (thin red line) is delivered in the last second of odour B; 2 repeats of reversal where shock is paired with odour A. (C) Abstract representation of the computational model as an electronic chip. The model receives the conditional (odour) and unconditional stimuli (electric shock) and produces the DAN and MBON responses using the incentive circuit and the dopaminergic plasticity rule. (D) The aversive olfactory conditioning experimental paradigm modified for testing the model. Odours A (coloured pink) and B (coloured yellow) are presented for two time steps each, in alternation, separated by 1 time-step fresh air, while the shock input forms the different phases and forgetting conditions: 1 repeat of pre-training, where no shock is delivered; 5 repeats of acquisition, where shock is delivered in the second time-step of odour B; 5 repeats of forgetting that can be either extinction (lightest shade of odour colour) where shock is not presented, unpaired (mid shade of odour colour) where shock (thin red line) is paired with the fresh air ‘break’, or reversal (dark shade of odour colour) where shock is paired with odour A. **Figure 4–Figure supplement 1.** The responses from all the recorded neurons in the *Drosophila melanogaster* mushroom body. **Figure 4–source data 1.** Imaging data of all the recorded neurons in the *Drosophila melanogaster* mushroom body.

### Microcircuits of the mushroom body

#### Susceptible & restrained memories

***Pavlowsky et al.*** (***2018***) identified a microcircuit in the mushroom body, where a punishment-encoding DAN (PPL1-γ1pedc) depresses the KC synapses onto an attraction-driving MBON (MBON-γ1pedc>α/β), which in turn inhibits the same DAN. They argue this is a memory consolidation mechanism, as the drop in the MBON response will reduce its inhibition of the DAN, enhancing the formation of the memory if the same odour-punishment pairing is continued. ***Felsenberg et al.*** (***2018***) further showed that the same MBON directly inhibits an avoidance-driving MBON (MBON-γ5β’2a), such that its activity increases (driving avoidance) after punishment as the inhibition is released. ***Figure 5***A shows these neurons in the mushroom body and ***Figure 5***B a schematic representation of their interconnections. Note, the MBON-IMBON inhibition is not reciprocal, rather we assume (see ***Figure 3*** and below) that there is a different microcircuit in which an avoidance-driving MBON inhibits an attraction-driving MBON. ***Figure 5***C-E show the responses of these neurons from experimental data (left) and from our model (right) during aversive conditioning (the paradigm shown in ***Figure 4***), which follow a similar pattern.

**Figure 5.**
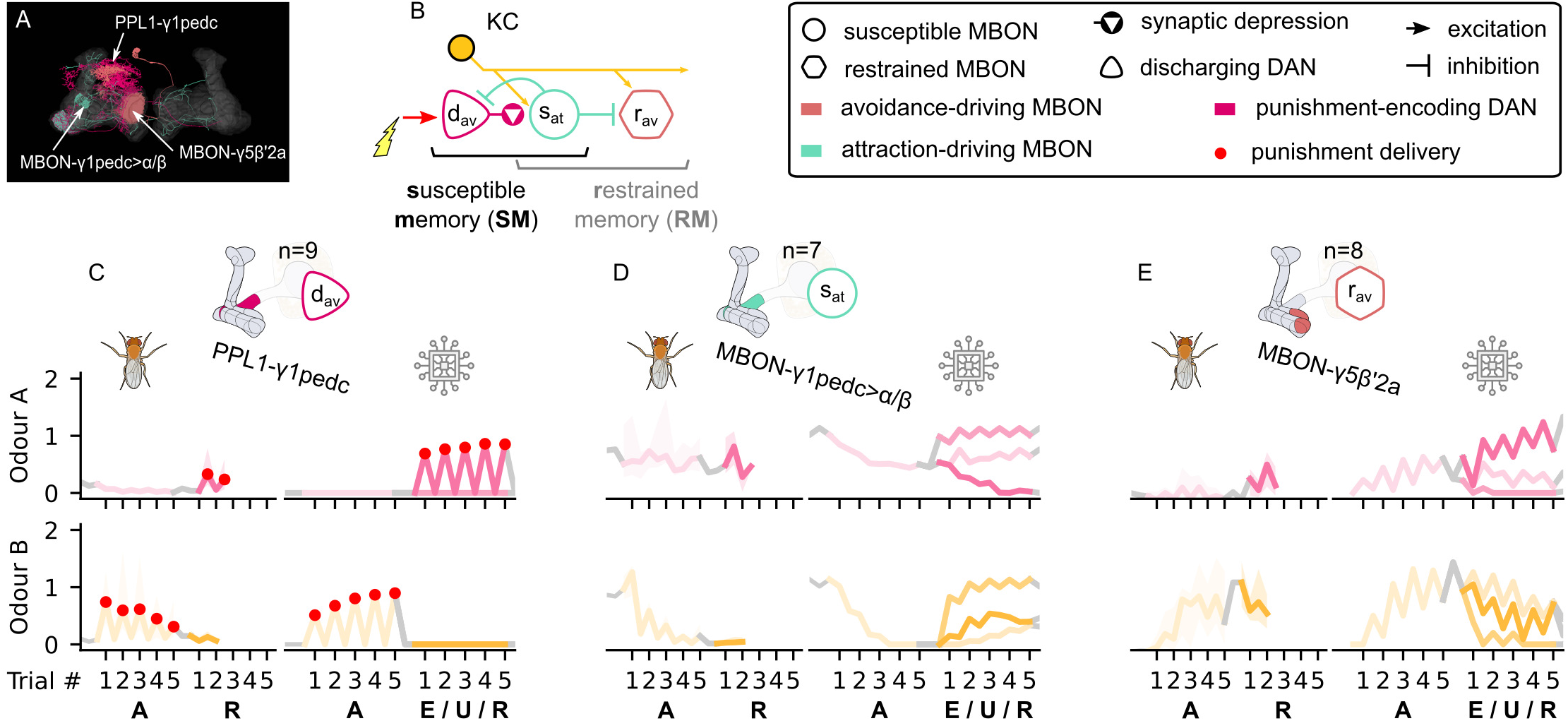
The susceptible and restrained microcircuits of the mushroom body. (A) Image of the attraction-driving susceptible and avoidance-driving restrained memory microcircuits made of the PPL1-γ1pedc, MBON-γ1pedc and MBON-γ5β’2a neurons - created using the Virtual Fly Brain software (***Milyaev et al., 2012***). (B) Schematic representation of the Susceptible & restrained memories microcircuits connected via the susceptible MBON. The responses of (C) the punishment-encoding discharging DAN, *d*_av_, (D) the attraction-driving susceptible MBON, *s*_at_, and (E) the avoidance-driving restrained MBON, *r*_av_, generated by experimental data (left) and the model (right) during the olfactory conditioning paradigms of ***Figure 4***D. Lightest shades denote the extinction, mid shades the unpaired and dark shades the reversal phase. For each trial we report two consecutive time-steps: the off-shock (i.e., odour only) followed by the on-shock (i.e., paired odour and shock) when available (i.e., odour B in acquisition and odour A in reversal phase) otherwise a second off-shock time-step (i.e., all the other phases). **Figure 5–Figure supplement 1.** The avoidance-driving susceptible memory, attraction-driving restrained memory and their responses. **Figure 5–Figure supplement 2.** The responses of the neurons of the incentive circuit using the connections introduced so far. **Figure 5–Figure supplement 3.** The KC→MBON synaptic weights of the neurons of the incentive circuit using the connections introduced so far.

Learning in this circuit is shown by the sharp drop (in both experimental data and model) of the response of MBON-γ1pedc>α/β (***Figure 5***D) to odour B already from the second trial of the acquisition phase. There is a similar drop of the response to odour A in the reversal phase. This rapid decrease is due to the depressing effect of the DAN on the KC→MBON synaptic weight. Note that we name this a ‘discharging DAN as the target synaptic strengths are high or ‘charged’ by default. However, due to our plasticity rule, if the US subsequently occurs without the CS (see unpaired phase in the model, for which we do not have experimental data), the MBON synaptic weights reset due to the recovery effect (see ***Figure 5***–***Figure Supplement 3***A — Odour B). This is consistent with the high learning rate and low retention time observed in ***Aso and Rubin*** (***2016***), and it results in a memory that is easily created and erased: a ‘susceptible memory’. The response of MBON-γ5β’2a (***Figure 5***E) can be observed to have the opposite pattern, i.e., it starts to respond to odour B from the second trial of acquisition as it is no longer ‘restrained’. Note however that the response it expresses, when the restraint is removed, also depends on its own synaptic weights for KC input, which as we will see, may be affected by other elements in the incentive circuit. In ***Figure 5***C, the experimental data shows a slight drop in the shock response (first paired with odour B, then with odour A) of the DAN, PPL1-γ1pedc, during the whole experiment, although it remains active throughout. We assume this drop may reflect a sensory adaptation to shock but have not included it in our model. Consequently, the model data shows a positive feedback effect: the DAN causes depression of the MBON response to odour, reducing inhibition of the DAN, which increases its response, causing even further depression in the MBON. Note this is opposite to the expected effects of reward prediction error.

Similar microcircuits in the mushroom body can be extracted from the connectome described in ***Aso et al***. (***2014a***) and ***Li et al.*** (***2020***) [also identified in larvae (***Eichler et al., 2017***)]. This leads us to the assumption that there are exactly corresponding susceptible and restrained memory microcircuits with opposite valence, i.e., a reward-encoding DAN that discharges the response to odour of an avoidance-driving MBON, which in turn releases its restraint on an attraction-driving MBON (see ***Figure 5***–***Figure Supplement 1***B and right side of the incentive circuit in ***Figure 3***, which mirrors the left side, with opposite valence). We further suggest specific identities for the neurons forming this circuit: PAM-γ4<γ1γ2 as the reward-encoding discharging DAN;MBON-γ4>γ1γ2 as the avoidance-driving susceptible MBON;and MBON-γ2α’1 as the attraction-driving restrained MBON (see ***Figure 5***–***Figure Supplement 1***A). The latter identification is based on the possibility of inhibiting connections from MBONs in the γ4 compartment to the ones in the γ2 compartment suggested by ***Aso et al***. (***2014a***) and ***Cohn et al.*** (***2015***). Although MBON-γ4>γ1γ2 is characterised by the Glutamate neurotransmitter, it is possible that it can inhibit MBON-γ2α’1 though glutamate-gated chloride channels (***Cleland, 1996***; ***Liu and Wilson, 2013***; ***McCarthy et al., 2011***).

#### Reciprocal short-term memories

***McCurdy et al.*** (***2021***) suggest that the attraction-driving restrained MBON in the previous circuit (MBON-γ2α’1) indirectly decreases the synaptic weights from KCs to the avoidance-driving restrained MBON (MBON-γ5β’2a) via an attraction-encoding DAN (PAM-β’2a). This microcircuit is also supported by ***Felsenberg et al.*** (***2018***) and ***Berry et al.*** (***2018***). ***Cohn et al.*** (***2015***) and ***Li et al.*** (***2020***) suggest that the corresponding avoidance-driving restrained MBON (MBON-γ5β’2a) excites an avoidance-encoding DAN (PPL1-γ2α’1), which closes the loop by affecting the KC connections to the attraction-driving restrained MBON, forming what we call the ‘reciprocal short-term memories’ microcircuit as shown in ***Figure 6***A (actual neurons in the mushroom bodies) and ***Figure 6***B (schematic representation of the described connections).

**Figure 6.**
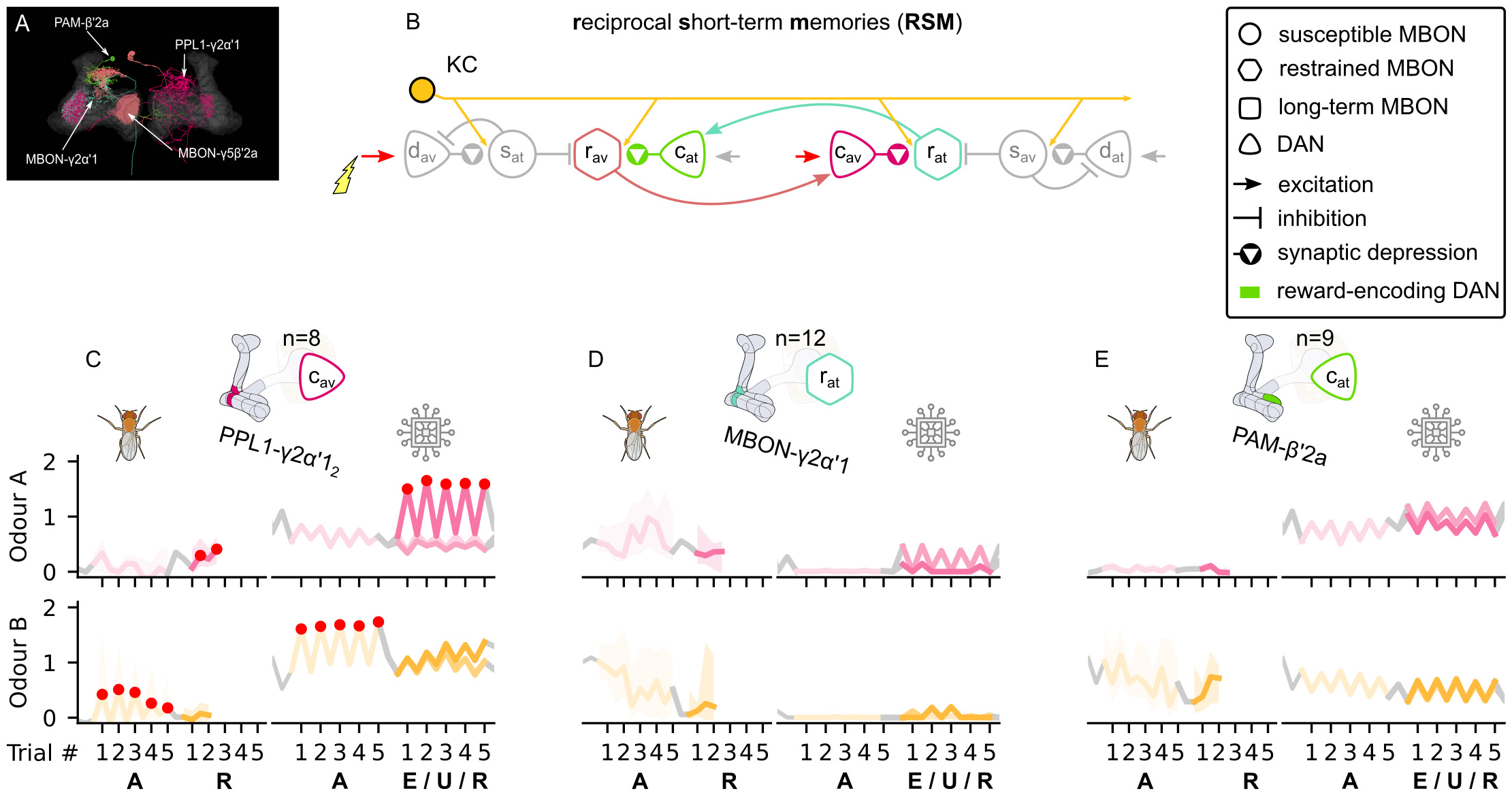
The reciprocal short-term memories microcircuit of the mushroom body. (A) Image of the reciprocal short-term memories microcircuit made of the MBON-γ5β’2a, PAM-β’2a, PPL1-γ2α’1 and MBON-γ2α’1 neurons - created using the Virtual Fly Brain software (***Milyaev et al., 2012***). (B) Schematic representation of the Reciprocal short-term memories microcircuit (coloured) connected to the susceptible memories via the restrained MBONs. The responses of (C) the punishment-encoding charging DAN, *c*_av_, the (D) attraction-driving restrained MBON, *r*_at_, and (E) the reward-encoding charging DAN, *c*_at_, generated by experimental data (left) and the model (right) during the olfactory conditioning paradigms of ***Figure 4***D. Lightest shades denote the extinction, mid shades the unpaired and dark shades the reversal phase. For each trial we report two consecutive time-steps: the off-shock (i.e., odour only) followed by the on-shock (i.e., paired odour and shock) when available (i.e., odour B in acquisition and odour A in reversal phase) otherwise a second off-shock time-step (i.e., all the other phases). **Figure 6–Figure supplement 1.** The responses of the neurons of the incentive circuit using the connections introduced so far. **Figure 6–Figure supplement 2.** The KC→MBON synaptic weights of the neurons of the incentive circuit using the connections introduced so far.

The ‘charging’ DANs, PAM-β’2a and PPL1-γ2α’1 (named after their LTM charging property - i.e., potentiation effect on another KC→MBON synapse - as we describe in the Long-term memory microcircuit), should be activated directly by reinforcement as well as by the restrained MBONs. This allows for memories to be affected directly by the reinforcement, but also by the expression of the opposite valence memories. The latter feature keeps the balance between the memories by automatically erasing a memory when a memory of the opposite valence starts building up and results in the balanced learning rate and retention time as observed in ***Aso and Rubin*** (***2016***). Because the memories in this pair of restrained MBONs are very fragile, we predict that these MBONs store short-term memories.

The effects of this circuit, as shown in ***Figure 6***C-E, are relatively subtle. During acquisition, the shock activates the punishment-encoding charging DAN (see ***Figure 6***C), which decreases the synaptic weights of the KC onto the attraction-driving restrained MBON (see ***Figure 6***–***Figure Supplement 2C***), but this cannot be seen in ***Figure 6***D because this MBON is already strongly inhibited (by the avoidance driving susceptible MBON). This low response means that the opposing reward-encoding charging DAN ***Figure 6***E is largely unaffected for this conditioning paradigm. In our model, the non-zero activity level of this DAN is a consequence of input from the long-term memory microcircuit which we describe next and the activation is similar for both odours because our networks starts in a balanced state (no preference for either odour). The different response to the two odours seen in the experimental data might therefore represent an unbalanced starting state of its long-term memory for these odours due to previous experiences of the fly.

#### Long-term memory

***Ichinose et al.*** (***2015***) describes a microcircuit where a reward-encoding DAN (PAM-α1) potentiates the KC→MBON synapses of MBON-α1, and MBON-α1 in turn excites PAM-α1. Using data from ***Li et al.*** (***2020***), we find numerous similar microcircuits, and in particular, MBONs that appear to have this recurrent arrangement of connectivity to the ‘charging’ DANs we have introduced to the circuit in the previous section. Specifically, we assume that the reward-encoding charging DAN (PAM-β’2a) can potentiate the response of the attraction-driving MBON-β2β’2a; and similarly the punishment-encoding charging DAN (PPL1-γ2α’1) potentiates the avoidance-driving MBON-α’1 (see ***Figure 7***A and C, ***Figure 7***B shows these connections schematically, with the KCs omitted for convenience). Crucially, these connections form positive feedback circuits — the DAN potentiates the response of the MBON to the odour, which increases its excitation of the DAN. As a consequence, even when the reinforcement ceases, the learning momentum can continue — this is the saturation effect of the learning rule (see ***Figure 2***D) and results in long-term memory (LTM) consolidation and enhancement.

**Figure 7.**
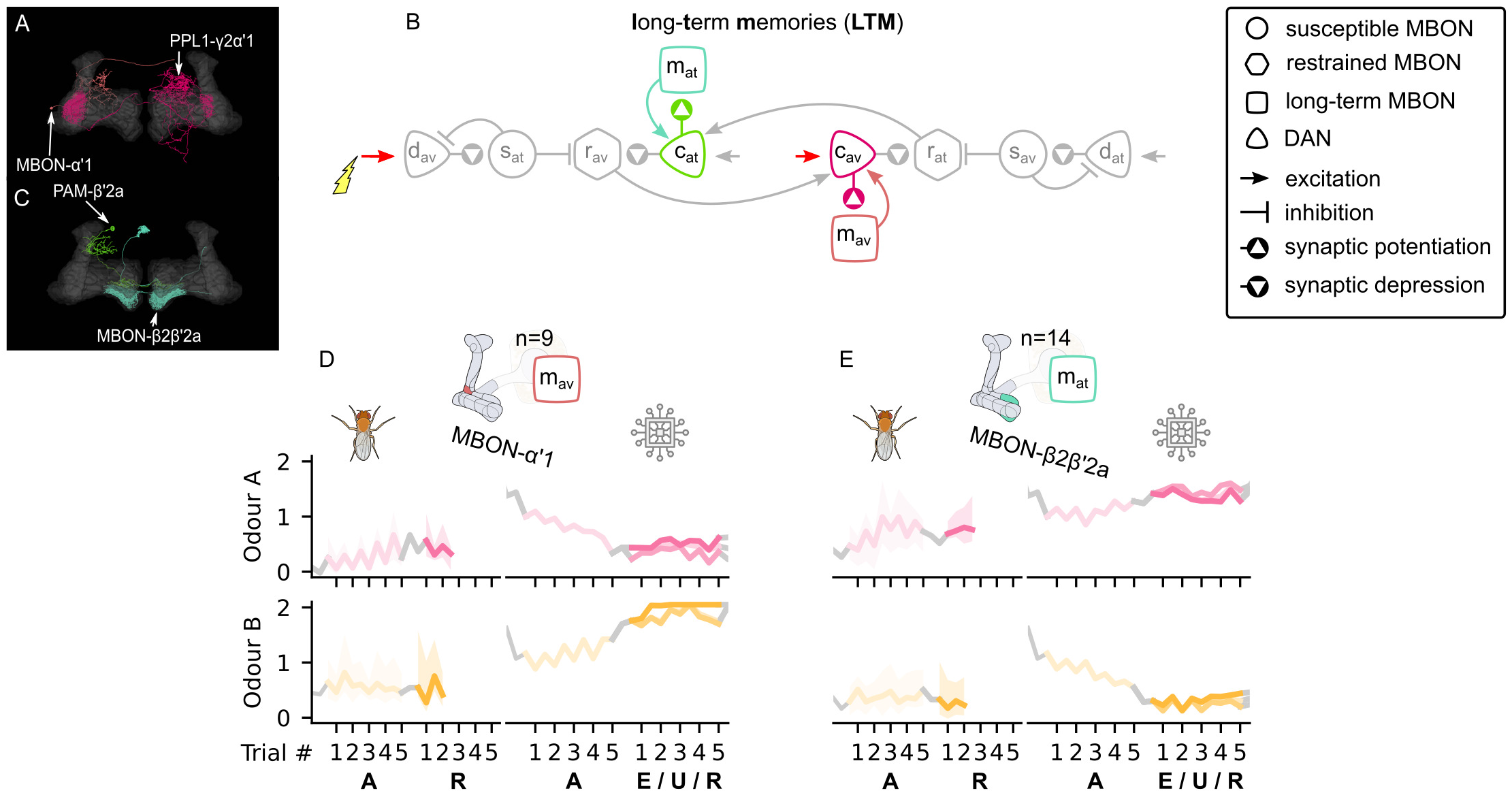
The long-term memory microcircuits of the mushroom body. (A) Image of the avoidance-encoding long-term memory microcircuit made of the MBON-α’ 1 and PPL1-γ2α’ 1 - created using the Virtual Fly Brain software (***Milyaev et al., 2012***). (B) Schematic representation of the Long-term memory microcircuits (coloured) connected to the RSM via the charging DANs. (C) Image of the attraction-encoding long-term memory microcircuit made of the MBON-β2β’2a and PAM-β2a - created using the Virtual Fly Brain software (***Milyaev et al., 2012***). The responses of the (D) avoidance-driving long-term memory MBON, *m*_av_, and (E) the attraction-driving long-term memory MBON, *m*_at_, generated by experimental data (left) and the model (right) during the olfactory conditioning paradigms of ***Figure 4***D. Lightest shades denote the extinction, mid shades the unpaired and dark shades the reversal phase. For each trial we report two consecutive time-steps: the off-shock (i.e., odour only) followed by the on-shock (i.e., paired odour and shock) when available (i.e., odour B in acquisition and odour A in reversal phase) otherwise a second off-shock time-step (i.e., all the other phases). **Figure 7–Figure supplement 1.** The responses of the neurons of the incentive circuit using the connections introduced so far. **Figure 7–Figure supplement 2.** The KC→MBON synaptic weights of the neurons of the incentive circuit using the connections introduced so far.

***Figure 7***D (right) demonstrates the charging of the avoidance-driving LTM MBON during the acquisition (for odour B) and its continued increase during the forgetting phases. However, these trends are not evident in the experimental data as illustrated in ***Figure 7***D (left). We suggest this is because responses of long-term memory neurons depend on the overall experience of the animal and are thus hard to predict during one experiment. For example, it could be the case that the animal has already built some long-term avoidance memory for odour A, such that its presentation without reinforcement in our experiment continues its learning momentum leading to the observed increasing response. Note that the decreasing response to odour A during acquisition in the model, as well as the observed effects in ***Figure 7***E for the attraction-driving LTM MBON, are due to influence from additional microcircuits to be described in the next section. ***Figure 7***–***Figure Supplement 1*** shows the responses of these neurons using only the microcircuits that have been introduced so far. In this case the responses of both LTM MBONs saturate instantly, which shows that another mechanism must exist and regulate them in order for them to become useful for the behaviour of the animal.

#### Reciprocal long-term memories

As described so far, once the Long-term memory microcircuit begins to charge, it will have a self-sustaining increase in the weights during odour delivery, preventing any subsequent adaptation to altered reward contingencies. To allow these weights to decrease, specifically, to decrease in response to charging of the LTM of opposite valence, we connect the two LTM MBONs via respective ‘forgetting DANs (see ***Figure 8***B). Note these forgetting DANs do not receive any direct reinforcement signals. Instead, as long as an LTM MBON is active, its respective forgetting DAN is also active and causes synaptic depression for the opposite LTM MBON (forgetting the learnt memory; see ***Figure 8***C and D). This counteracts any potentiation effect due to the LTM MBON’s respective charging DAN (see ***Figure 8***–***Figure Supplement 2***E and F). As a consequence, sustained reinforcement of one valence can gradually overcome the positive feedback of the LTM circuit of opposite valence, causing the charging momentum to drop and eventually negate. The long-term memories are thus in long-term competition.

**Figure 8.**
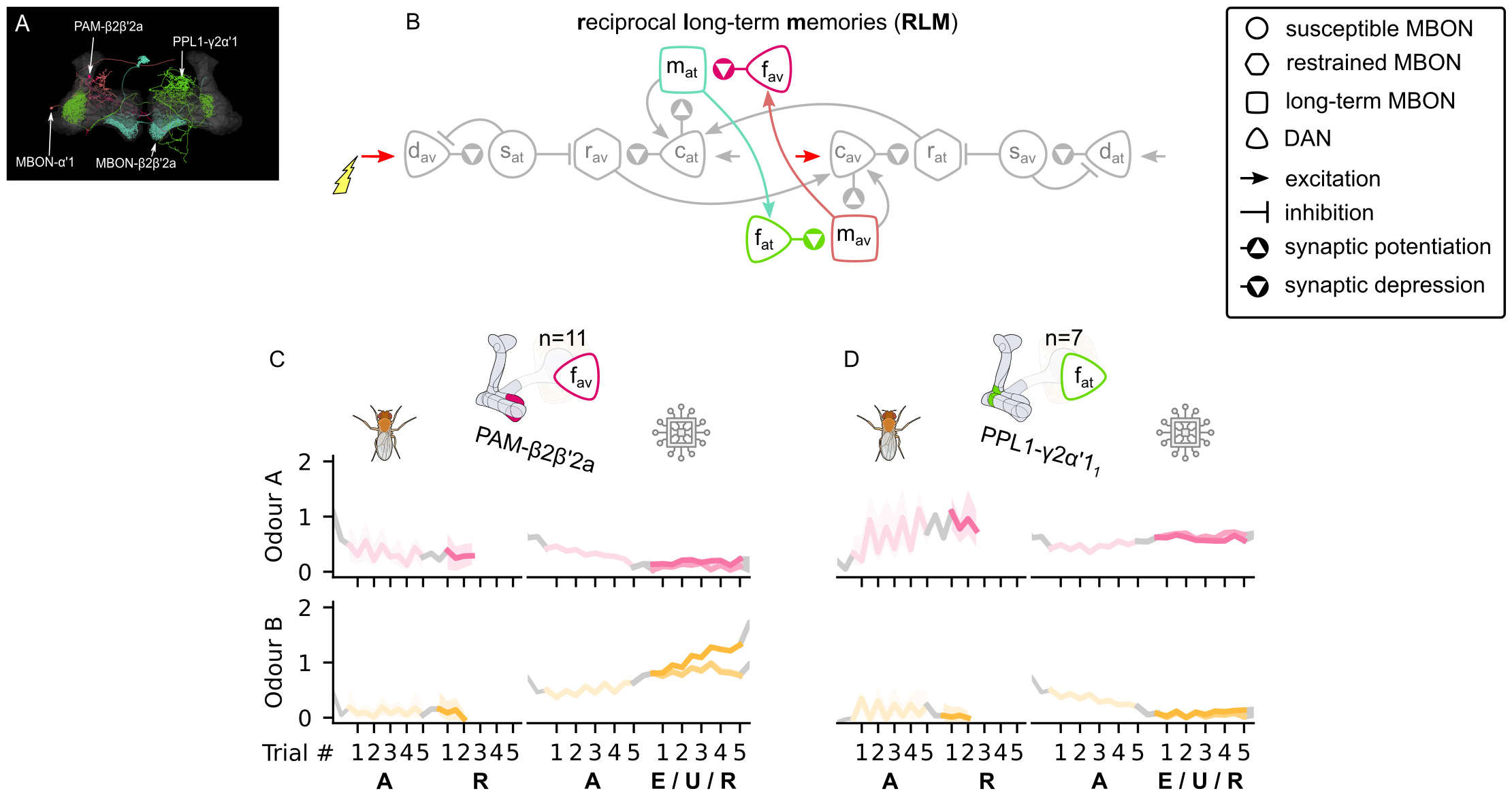
The reciprocal long-term memories microcircuit of the mushroom body. (A) Image of the reciprocal long-term memory microcircuit in the mushroom body made of the MBON-α’1, PAM-β2β’2a, MBON-β2β’2a and PPL1-γ2α’1 - created using the Virtual Fly Brain software (***Milyaev et al., 2012***). (B) Schematic representation of the Reciprocal long-term memories microcircuit (coloured) The responses of (C) the punishment-encoding forgetting DAN, *f*_av_, the (D) reward-encoding forgetting DAN, *f*_at_, generated by experimental data (left) and the model (right) during the olfactory conditioning paradigms of ***Figure 4***D. Lightest shades denote the extinction, mid shades the unpaired and dark shades the reversal phase. For each trial we report two consecutive time-steps: the off-shock (i.e., odour only) followed by the on-shock (i.e., paired odour and shock) when available (i.e., odour B in acquisition and odour A in reversal phase) otherwise a second off-shock time-step (i.e., all the other phases). **Figure 8–Figure supplement 1.** The responses of the neurons of the incentive circuit using the connections introduced so far. **Figure 8–Figure supplement 2.** The KC→MBON synaptic weights of the neurons of the incentive circuit using the connections introduced so far.

We have identified the reciprocal long-term memories microcircuit of ***Figure 8***B in the descriptions of ***Aso et al***. (***2014a***) and ***Li et al.*** (***2020***), where MBON-α’1 is the avoidance-driving LTM MBON, MBON-β2β’2a is the attraction-driving LTM MBON, PAM-β2β’2a is the avoidance-driving forgetting DAN, and PPL1-γ2α’1 is the attraction-driving forgetting DAN, as shown in ***Figure 8***A. One problem with this identification is that there is only one PPL1-γ2α’1 per hemisphere and we have already suggested it should be identified as the punishment-encoding charging DAN in our model. However, there are multiple axon terminals of this neuron in the mushroom body (e.g., MB296B_1_ and MB296B_2_) and each one of them seems to communicate a different response (see ***Figure 4***–***Figure Supplement 1*** — row 5, columns 6 and 7). Interestingly, the responses communicated by the MB296B_1_ terminal are close to the ones produced by the punishment-encoding charging DAN (see ***Figure 6***C) and the ones of the MB296B_2_ are close to the ones produced by the attraction-driving forgetting DAN (see ***Figure 8***D). This implies that different axons of the same DA neuron might create responses that depend on where the axon terminates and actually work as separate processing units. ***Figure 8***C and D show that the reconstructed responses of these neurons from our model are surprisingly similar to the ones observed in the data.

#### Memory assimilation mechanism

The forgetting DANs allow the developing LTM of one valence to induce forgetting of the LTM of the opposite valence. However the forgetting DANs can also be used for another critical function to maintain flexibility for future learning, which is to erase the memory of the same valence from their respective restrained MBONs. We thus predict that the forgetting DANs also suppress the KC synaptic weights of their respective restrained MBONs, forming the ‘memory assimilation mechanism’ (MAM) microcircuit (see ***Figure 9***C). This effectively allows memory transfer between the restrained and the LTM MBONs, enhancing both the adaptability and the capacity of the circuit. This effect can be observed in the difference of the responses of the same neurons in ***Figure 7***, ***Figure 8*** and ***Figure 8***–***Figure Supplement 1***, where the restrained memory becomes weaker as the LTM becomes stronger, driven by the respective forgetting and charging DANs.

**Figure 9.**
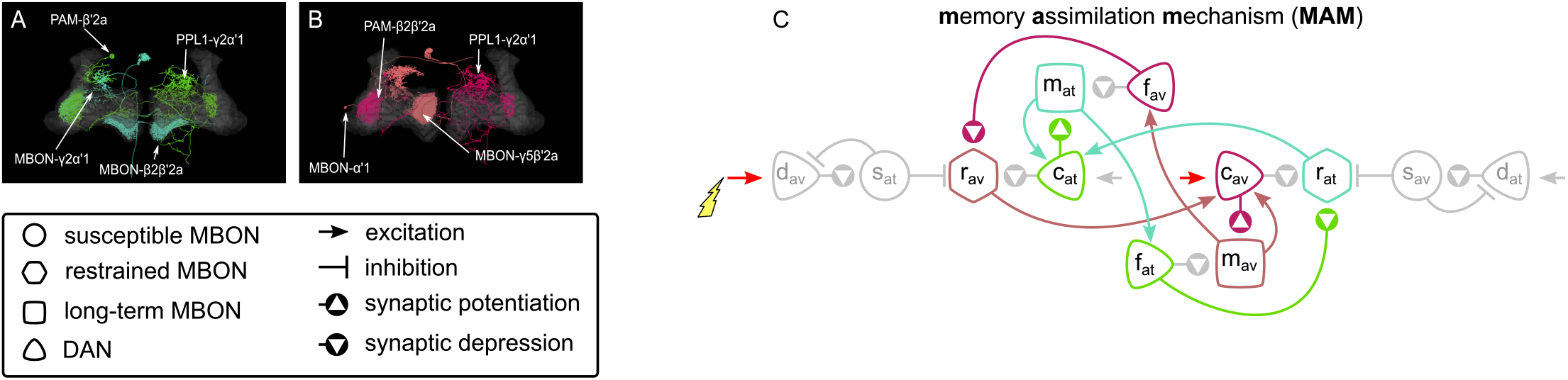
The memory assimilation mechanism microcircuit of the mushroom body. (A) Image of the avoidance-specific MAM microcircuit in the mushroom body made of the MBON-γ5β’2a, PPL1-γ2α’1, MBON-α’1 and PAM-β2β’2a - created using the Virtual Fly Brain software (***Milyaev et al., 2012***). (B) Image of the attraction-specific MAM microcircuit in the mushroom body made of the MBON-γ2α’1, PAM-β’2a, MBON-β2β’2a and PPL1-γ2α’1 - created using the Virtual Fly Brain software (***Milyaev et al., 2012***). (B) Schematic representation of the Memory assimilation mechanism microcircuits (coloured). The forgetting DANs connect to the restrained MBONs of the same valence, hence increasing LTM strength reduces (assimilates) the restrained memory, constituting the MAM microcircuits. **Figure 9–Figure supplement 1.** The KC→MBON synaptic weights of the neurons of the incentive circuit using all its connections. **Figure 9–Figure supplement 2.** The reconstructed responses of the neurons of the incentive circuit using the RPE learning rule. **Figure 9–Figure supplement 3.** The KC→MBON synaptic weights of the neurons of the incentive circuit using the RPE learning rule.

The depression effect of the forgetting DANs on the KC→restrained MBON synapses of the same valence is supported by ***Aso et al***. (***2014a***). More specifically, the avoidance-driving forgetting DAN we have identified as PAM-β2β’2a modulates the KC→MBON-γ5β’2a synapses, while for the attraction-driving forgetting DAN, PPL1-γ2α’1, modulates the KC→MBON-γ2α’1 synapses, as show in ***Figure 9***A and B respectively.

### Modelling the behaviour

In the incentive circuit, three MBON types drive attraction and three avoidance. This results in six driving forces, for each available odour (see ***Figure 10***). A simple ‘behavioural’ readout (used in many previous models) would be to take the sum of all attractive and aversive forces at some time point as a measure of the probability of animals ‘choosing’ odour A or B, and compare this to the standard two-arm maze choice assay used in many Drosophila studies. Following this approach and using the summarised data collected by ***Bennett et al.*** (***2021***), we have tested the performance of our model in 92 olfactory classical conditioning intervention experiments from 14 studies (***Felsenberg et al., 2017***; ***Perisse et al., 2016***; ***Aso and Rubin, 2016***; ***Yamagata et al., 2016***; ***Ichinose et al., 2015***; ***Huetteroth et al., 2015***; ***Owald et al., 2015***; ***Aso et al., 2014b***; ***Lin et al., 2014***; ***Plaçais et al., 2013***; ***Burke et al., 2012***; ***Liu et al., 2012***; ***Aso et al., 2010***; ***Claridge-Chang et al., 2009***), i.e., the observed effects on fly learning of silencing or activating specific neurons, including positive and negative reinforcements. The Δ*f* predicted from the incentive circuit correlated with the ones reported from the actual experiments with correlation coefficient *r* = 0.76, *p* = 2.2 × 10^−18^ (***Figure 3***–***Figure Supplement 1***).

**Figure 10.**
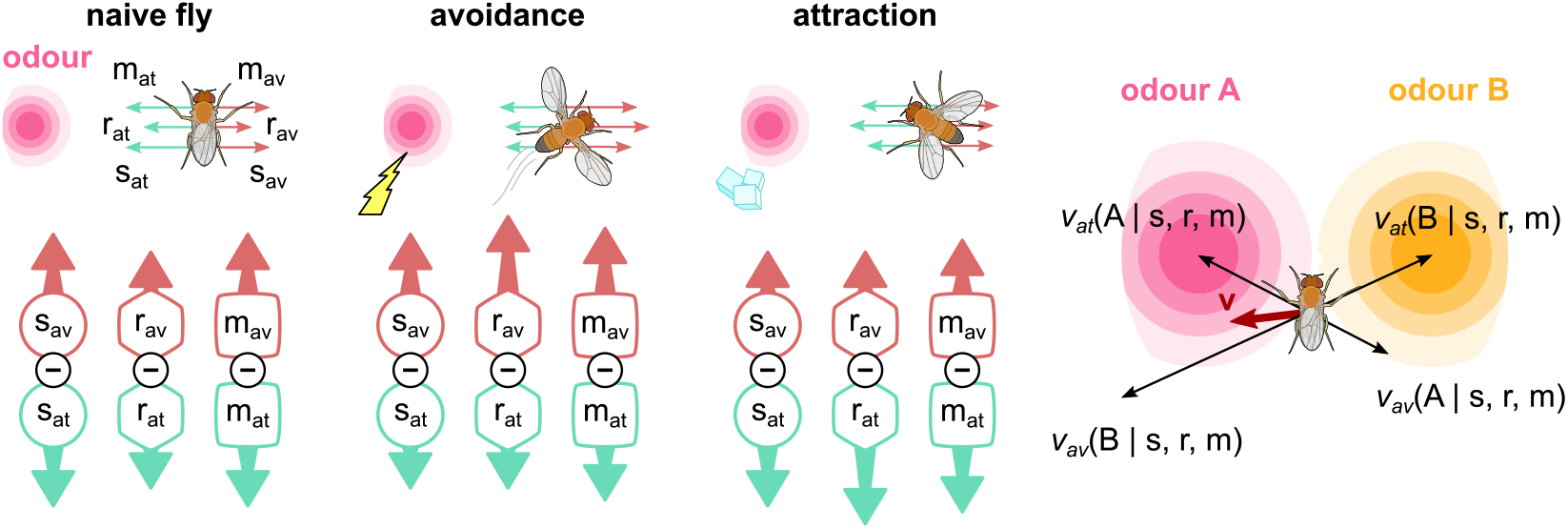
The activity of the six MBONs is translated into forces that drive a simulated fly towards or away from odour sources. For naive flies, the forces are balanced. When electric shock is paired with an odour, the balance changes towards the avoidance-driving MBONs, which drives the fly directly away from that odour. When sugar is paired with an odour, the balance changes to attraction, driving the fly towards that odour. Combining all attractive and repulsive forces for each odour source currently experienced by the fly produces an overall driving force, **v**, which determines the fly’s behaviour.

However, classical conditioning does not allow us to explore the full dynamics of the circuit as animals simultaneously explore, learn, express learning and forget, while moving through a world with odours. Therefore, we further simulate the behaviour produced by the incentive circuit in operant conditioning: simulated flies are placed in a virtual arena, where they are exposed to two odour gradients, of different strengths, and variously paired with reinforcements. As we have full access to the neural responses, the synaptic weights and the position of the simulated flies for every time-step, this allows us to identify different aspects of the produced behaviour and motivation, including the effect of the LTM on the behaviour and whether ‘choice’ occurs because the animal is attracted by one odour or repulsed by the other. We can then derive a behavioural preference index based on the time the simulated flies spent exposed in each odour during relevant time periods. ***Figure 11*** summarises our experimental set-up and results, while details about how we generate the presented behaviours are in the Modelling the behaviour section of the Methods.

**Figure 11.**
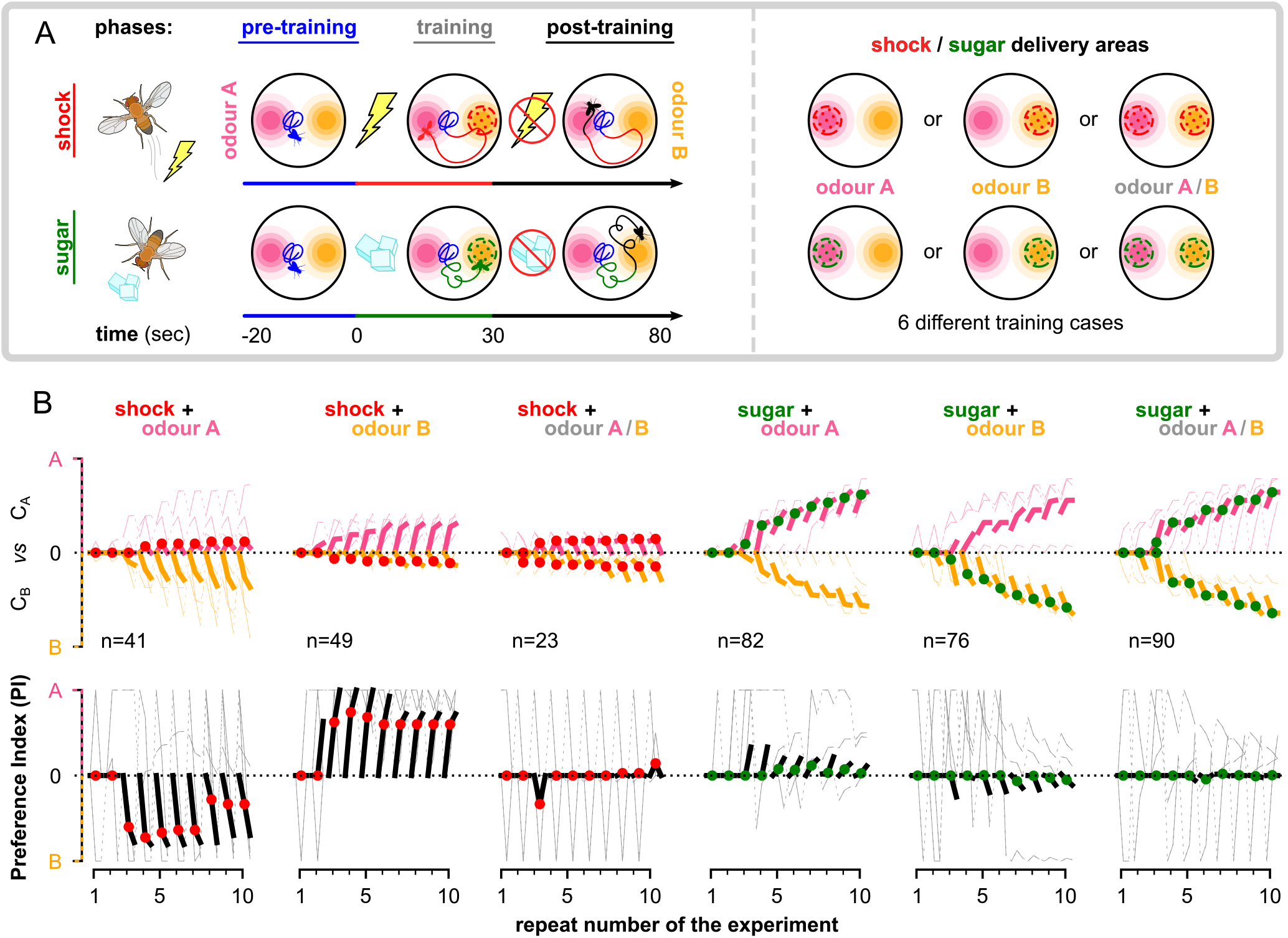
The behaviour of the animal controlled by its neurons during the freely-moving flies simulation. The *n* = 100 simulated flies are exposed to a mixture of two odours, whose relative intensity depends on the position of the simulated flies in space. (A) Each experiment lasts for 100 sec where: the flies are placed at the centre of the arena in time-step *t* = −20 sec. During the first 20 sec (pre-training phase, *t* ∈ [-20, 0)) the flies explore the arena without any reinforcement (blue tracks). In the next 30 sec (training phase, *t* ∈ [0, 30)) they conditionally receive reinforcement under one of the six training cases shown on the right: using sugar (green) or shock (red); and reinforcing around odour A (shock + odour A), odour B (shock + odour B) or both odours (shock + odour A/B). During the last 50 sec (post-training phase) *t* ∈ [30, 80)) they continue being exposed to the odours without receiving a reinforcement (black tracks). We repeat this experiment (including all its phases) for 10 times in order to show the effects of the long-term memory in the behaviour. (B) Behavioural summary of a subsets of simulated flies, that visited both odours at any time during the 10 repeats. Columns show the different conditions and the population that was recorded visiting both odours. Top row: the normalised cumulative time spend exposed in odour A (pink lines) or odour B (yellow lines - note this line is reversed). For each repeat we present 3 values (average over all the pre-training, training and post-training time-steps respectively) where the values associated with the training phase are marked with red or green dots when punishment or reward has been delivered to that odour respectively. Thin lines show 3 representative samples of individual flies. Thick lines show the median over the simulated flies that visited both odours. Bottom row: the preference index (PI) to each odour extracted by the above cumulative times. **Figure 11–Figure supplement 1.** The mean KC→MBON synaptic weights over the simulated flies that visited both odours. **Figure 11–Figure supplement 2.** Behavioural summary of simulated flies grouped by the areas that they visited. **Figure 11–Figure supplement 3.** Behavioural summary of simulated flies when controlled by the different types of MBONs. **Figure 11–Figure supplement 4.** Paths of 100 simulated flies when using the dopaminergic plasticity rule and during all 10 repeats. **Figure 11–Figure supplement 5.** Behavioural summary of simulated flies when using the reward prediction error plasticity rule. **Figure 11–Figure supplement 6.** Paths of 100 simulated flies when using the RPE plasticity rule and during all 10 repeats. **Figure 11–Figure supplement 7.** The mean KC→MBON synaptic weights when using the RPE plasticity rule over the simulated flies that visited both odours.

In ***Figure 11***–***Figure Supplement 4***, we can see that most simulated flies do not visit any of the regions that an odour can be detected in the first repeats, and therefore, in ***Figure 11***B, we start seeing an effect in the averaged statistics after the second repeat of the experiment. However, in the first couple of repeats, the individual paths already show a small tendency to the expected behaviour of the flies: avoid the punished region and approach the rewarded one. Due to the unpredictable behaviour of the individual flies, in ***Figure 11***B we summarise only times from simulated flies that have visited both odours for at least one second. In later repeats of the experiment, the PI shows that (on average) flies prefer the non-punished and rewarded odours. When both of them are punished or rewarded they equally prefer none or both respectively. Note that the above result does not mean that each fly spends equal time in both odours, but that most probably some flies choose to spend more time with the one and some with the other (as show from the individual cumulative durations in ***Figure 11***B), but their population is equal. It is interesting that almost in every repeat the flies are neutral about the odours during pre-training (time-step before the reinforced one — marked with red or green), show a relatively small effect during training and bigger effect during post-training. This might be because in every repeat of the experiment they are initialised in the centre, so they spend some time randomly exploring before they detect an odour.

By looking at the PIs of ***Figure 11***B, we see a strong effect when electric shock is paired with odour A or B, but not very strong otherwise. We also see a smaller PI for flies experiencing sugar than the ones that experience electric shock, which is inline with experimental data (***Krashes and Waddell, 2011a***,b). When shock is paired with both odours we expect that the simulated flies will try to minimise the time spent exposed to any of them, which is precisely what we see in the coloured lines. In contrast, simulated flies seem to increase the time spend in both odours when paired with sugar with a slight preference towards the reinforced odour. In general, our results show that (in time) the simulated flies seem to develop some prior knowledge about both odours when experienced at least one of them with reinforcement (see ***Figure 11***B and ***Figure 11***–***Figure Supplement 2A***), which we suggest is because of their overlapping KCs associated with both odours. We believe that this leads into self-reinforcement, which means that when the animal experiences the non-reinforced odour it will automatically associate the reinforcement associated with the overlapping KCs to all the KCs associated with this odour, which is effectively a form of second-order conditioning.

From the summarised synaptic weights shown in ***Figure 11***–***Figure Supplement 1***, we can see that the susceptible MBONs immediately block the simulated flies from approaching the punishing odours, while they allow them approach the rewarding ones, which results in the smaller PI shown in sugar-related experiments compare to the shock-related ones, as discussed before. This is partially because of the lack of reciprocal connections between the opposing susceptible MBONs, and it can be verified through the appetitive conditioning, where the synaptic weights seem to change as the simulated flies now prefer the reinforced odour site. Susceptible MBONs convulsively break the balance between attraction and avoidance created by the restrained and LTM MBONs, also affecting their responses, and allowing STM and as a result LTM formation even without the presence of reinforcement. ***Figure 11***–***Figure Supplement 1*** also show that the restrained MBONs seem to play an important role during the first repeats (up to 5), but then they seem to reduce their influence giving up the control to the LTM MBONs, which seem to increase their influence with time. This is partially an effect of the MAM microcircuit, which verifies its function and the role of the restrained MBONs as storing short-term memories. ***Figure 11***–***Figure Supplement 3*** shows that the different types of MBONs alone are also capable of controlling the behaviour. However, they seem to better work when combined, as they complement one another in different stages, e.g., during early or late repeats and crucial times.

### Dopaminergic plasticity rule vs reward prediction-error

We have already shown that our novel dopaminergic plasticity rule (DPR) and the connectome of the incentive circuit build a powerful model for memory dynamics and behavioural control. In order to verify the importance of our DPR in the model, we run the same experiments by replacing it with the reward prediction error (RPE) plasticity rule (***Rescorla and Wagner, 1972***).

The idea behind RPE is that the fly learns to predict how rewarding or punishing a stimulus is by altering its prediction when this does not match the actual reward or punishment experienced (***Zhao et al., 2021***). This can be adapted to the mushroom body circuit by assuming for a given stimulus represented by KC activation, the MBON output is the prediction, and the KC→MBON synaptic weights should be altered (for the active KC) proportionally to the difference between the MBON output and the actual reinforcement signalled by the DAN. In The reward prediction error plasticity rule, we show how our DPR can be replaced with the RPE (as described above) in our model. Note that this rule allows updates to happen only when the involved KC is active, implying synaptic plasticity even without DAN activation but not without KC activation, which is in contrast with our DPR and recent findings (***Berry et al., 2018***; ***Hige et al., 2015***) [also in larva (***Schleyer et al., 2018***, ***2020***)].

This effect, i.e., learning when the KC is active even without DAN activation, is visible in ***Figure 9***–***Figure Supplement 2*** and ***Figure 9***–***Figure Supplement 3***, where we can see that, for the susceptible MBONs, the synaptic weights recover every time before the shock delivery, when the odour is presented alone, resulting in no meaningful learning and cancelling their susceptible property. Restrained MBONs look less affected (at least in this experimental set-up), while the long-term memory MBONs lose their charging momentum obtained by the saturation effect, resulting in more fragile memories. Furthermore, due to the KC (instead of dopamine) gating of this plasticity rule, the responses during the unpaired and extinction conditions look identical in all neurons, while the reversal makes a difference only on the responses to odour A. In general, the responses reproduced using the RPE plasticity rule have none of the properties of our model that have been shown earlier and also they cannot explain the dynamics of the responses recorded from animals.

In contrast to the responses, the behaviour of the simulated flies (as shown in ***Figure 11***–***Figure Supplement 5*** and ***Figure 11***–***Figure Supplement 6***) is less affected by the plasticity rule: we still see a preference to the non-punished or rewarded odours. However, there are some details in the behaviour that are different and some properties of the model that need to be mentioned. First, we see that the simulated flies now spend more time in the punished odours (compare to the non-punished ones), which might look as adaptation (in PI level), but it is actually forgetting about the odour. ***Figure 11***–***Figure Supplement 7*** shows that synaptic weights targeting the restrained and LTM MBONs are dramatically depressed during the first three repeats and are unable to recover whatsoever, which means that this part of the circuit is knocked out by then. Hence, the behaviour is controlled solely by the susceptible MBONs, which now look more like LTM MBONs that are not reciprocally connected. Furthermore, the synaptic weights associating the odours to both motivations seem to constantly decrease, which makes us believe that both susceptible MBONs will have the same future as the restrained and LTM ones, but it will just take longer. Therefore, we see that although the RPE predicts a reasonable behaviour for inexperienced (or minor experienced) simulated flies, it could gradually result in a meaningless behaviour for experienced flies.

## Discussion

We have shown that the combination of our novel’dopaminergic plasticity rule’ (DPR) with the incentive circuit (IC) of the mushroom body is able to generate similar neural responses and behaviours to flies in associative learning and forgetting paradigms. Regarding our model, we provide evidence for the existence of all hypothesised connections and suggest that at least 3 types of mushroom body output (susceptible, restrained and long-term memory) and 3 types of dopaminergic neurons (discharging, charging and forgetting) exist in the fruit fly brain, discriminated by their functionality. As we show, this forms a unified system for rapid memory acquisition and transfer from short-term to long-term memory, which could underly the ability to make exploration/exploitation trade-offs.

### Advantages of the dopaminergic plasticity rule

The proposed dopaminergic plasticity rule (DPR), while remaining very simple, allows the animal to express a variety of behaviours depending on their experience. The rule remains local to the synapse, that is, it depends only on information that is plausibly available in the presynaptic area of the KC axon (***Equation 1***): the activity of the KC, the level of DA, and the deviation of the current ‘weight’ from a set-point ‘resting weight’. We note that it was not possible to obtain good results without this third component to the rule, although the underying biophysical mechanism is unknown; we speculate that it could involve synapsin as it has a direct role in regulating the balance of reserve and release vesicle pools, and is required in the MB for associative learning (***Michels et al., 2011***). The rule also introduces a bidirectional ‘dopaminergic factor’ based on the results of ***Handler et al.*** (***2019***) who showed the combination of DopR1 and DopR2 receptor activity can result in depression or potentiation of the synapse. In our plasticity rule, a positive or negative dopaminergic factor combined with active or inactive KCs leads to four possible effects on the synapse: depression, potentiation, recovery and saturation. This allows substantial flexibility in the dynamics of learning in different MB compartments.

In particular, the saturation allows LTM MBONs to consolidate their memories and makes it very hard to forget. This only occurs for consistently experienced associations, which then become strongly embedded. Only a persistent change in the valence of reinforcement experienced with a given stimuli can reset the activity of LTM MBONs through the Reciprocal long-term memories microcircuit, which equips the circuit with flexibility even in the long-term memories. Further, the fact that the DPR allows short-term (restrained) and long-term memories to interact through the Memory assimilation mechanism, increases the capacity of the circuit. Whatever the restrained MBONs learn is eventually assimilated by the LTM MBONs, opening up space for the formation of new memories in the restrained MBONs. When combined with sparse coding of odours in a large number of KCs, the LTM MBONs can store multiple memories, for different odours. Short-term experience might occasionally affect the behaviour when the susceptible and restrained MBONs learn something new, and hence mask the LTM output, but eventually this will be smoothly integrated with the previous experience in the LTM MBONs. The DPR plays an important role in this mechanism, as we saw earlier, and the connectivity alone is not enough for it to work properly.

By contrast, the reward prediction-error (RPE) plasticity rule lacks this flexibility, and fails to maintain useful long-term memories when applied to the same circuit architecture. A literal interpretation of RPE for the MB would require that the difference (error) between the post-synaptic MBON activity and the DA level is somehow calculated in the pre-synaptic KC axon. This seems inconsistent with the observation that learning is generally unaffected by silencing the MBONs during acquisition (***Hige et al., 2015***; ***Krashes et al., 2007***; ***Dubnau et al., 2001***; ***McGuire et al., 2001***). Alternatively (and not directly requiring MBON activity in the KC plasticity rule) the RPE could be implemented by circuits (***Bennett et al., 2021***; ***Springer and Nawrot, 2021***; ***Eschbach et al., 2020***) in which DANs transmit an error signal computed by their input reinforcement plus the opposing feedback from MBONs (i.e., MBONs inhibit DANs that increase the KC→MBON synaptic weights, or they excite those that suppress the synaptic weights). However, although the evidence for MBON-DAN feedback connections is well-grounded, it is less clear that they are consistently opposing. For example, in the microcircuits we have described, based on neurophysiological evidence, some DANs that depress synaptic weights receive inhibitory feedback from MBONs (***Pavlowsky et al., 2018***) and some DANs that potentiate synaptic weights receive excitatory feedback from DANs (***Ichinose et al., 2015***). As we have shown, the DPR is able to operate with this variety of MBON-DAN connections. Note that, by using the appropriate circuit, i.e., positive MBON-DAN feedback to depressing DANs, our DPR could also have an RPE effect. Although the proposed incentive circuit does not include such connections, it is still possible that they exist.

### The conditioning effects of the model

During the past decades, a variety of learning effects have been investigated in flies, including forward and backward (relief) conditioning, first- and second-order conditioning and blocking, which we could potentially use to challenge our model. In Derivation of the dopaminergic plasticity rule, we demonstrate that our model supports the backward (or relief) conditioning results presented in ***Handler et al.*** (***2019***). Backward conditioning is when the reinforcement is delivered just before the odour presentation and it is based on the time-dependency between the two stimuli. ***Handler et al.*** (***2019***) suggest that the backward conditioning is a mechanism driven by ER-Ca^2+^ and cAMP in a KC→MBON synapse, when a single DAN releases DA on it. In our model, we assume that different time-courses in the response of DopR1 and DopR2 receptors cause the different patterns of ER-Ca^2+^ and cAMP, resulting in the formation of opposite associations for forward and backward conditioning. We note however that in our model, the effect also requires that the target MBON inhibits the respective DAN (as in our susceptible memory microcircuits) altering the time course of neurotransmitter release. This may suggests backward conditioning does not occur in all MB compartments. We believe this mechanism for backward conditioning is better supported than the hypothesised mechanism of post-inhibitory rebound in opposing valence DANs presented in ***Adel and Griffith*** (***2021***), although some role for both mechanisms is possible.

Backward conditioning can be distinguished from the unpaired conditioning effect; the latter involves the presentation of reinforcement and a specific odour in alternation with less temporal proximity. It has been observed ***Jacob and Waddell, 2020***; ***Schleyer et al., 2018***) that this procedure will produce a change in response to the odour that is opposite in valence to the reinforcement, e.g., approach to an odour that is ‘unpaired’ with shock. Note that this effect can be observed both in standard two odour CS+/CS- training paradigms (where an altered response to CS-, in the opposite direction to CS+, is often observed) but also in single odour unpaired paradigms. Surprisingly, our model also produces unpaired conditioning, notably through a different mechanism than backward conditioning. When DANs are activated by a reinforcement without KC activation, the weights of all KCs are potentially altered, e.g., restored towards their resting weight or slightly potentiated. This alteration means that subsequent presentation of odour alone can be accompanied by MBON-driven activation of DANs, resulting in specific alteration of the weights for the presented odour. In the example of ***Figure 12***, odour A starts to self-reinforce its attractive LTM when presented in alternation with shock, and will be preferred to an alternative odour B in subsequent testing. However, repeated presentation of other odours during testing, without further shock, might lead to generalisation (equal preference to all experienced odours).

**Figure 12.**
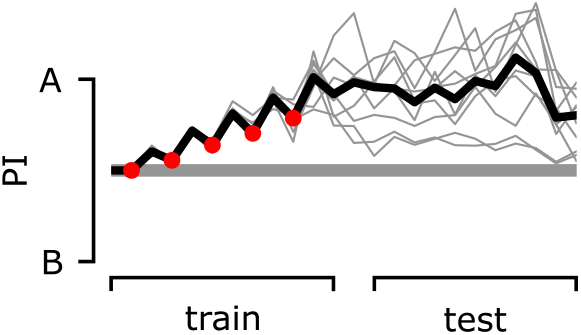
The preference index (PI) of the agent during the classic unpaired conditioning paradigm. During the training phase, we deliver electric shock or odour A alternately. During the test phase, we deliver odour A and odour B alternately. The preference index (PI) is calculated by using the MBON responses for each odour.

The self-reinforcing property of the positive feedback in the LTM microcircuit can also account for second-order conditioning. If a motivation has been associated to an odour, MBONs related to that motivation will have increased activity when the odour is delivered, even in the absence of reinforcement. In the LTM microcircuit, the positive MBON-DAN connection will consequently activate the charging DAN, so any additional cue (or KC activity) presented alongside the learned odour will also experience increase in the respective KC→MBON weights, creating a similar charging momentum and resulting in a second-order association. Perhaps surprisingly, this predicts that second-order conditioning might happen directly in the long-term memory microcircuit without being filtered by the susceptible and restrained memories first. This would be consistent with the observation that second-order conditioning in flies requires strong induction of the first-order memory and that first-order memory does not appear to be extinguished by the absence of reinforcement during second-order training (***Tabone and Belle, 2011***).

Finally, although we have not tested it explicitly here, it is clear that our plasticity rule (unlike RPE) would not produce blocking. The blocking effect, as described by ***Kamin*** (***1967***), is when the conditioning to one stimulus subsequently blocks any conditioning to other elements of a mixture including that stimulus. Under RPE learning, this is explained by the first stimulus already correctly predicting the reinforcer, so there is no error to drive a change in the weights. Using the DPR, the updates are local to the synapse and do not depend on a calculation of errors summarised across different odour identities, so blocking does not happen, which is consistent with the observed behaviour of fruit flies (***Young et al., 2011***; ***Brembs and Heisenberg, 2001***). Although presentation of a learned odour along with a novel odour might, through feedback from the MBONs, alter the DAN responses to the reinforcement, in our circuit this is not generally an opponent feedback so will not cancel the reinforcing effects for the novel odour. This also highlights the difference of our Susceptible & restrained memories and Long-term memory microcircuits from the RPE circuits described ***Bennett et al.*** (***2021***); ***Springer and Nawrot*** (***2021***); ***Eschbach et al.*** (***2020***) and ***Zhao et al.*** (***2021***). Nevertheless, as ***Wessnitzer et al.*** (***2012***) and later ***Bennett et al.*** (***2021***) suggest, the fact that blocking has not been observed in fruit flies could also be explained by the way that the mixture of odours is represented by the KCs, i.e., that it might not be simply the superposition of the activity patterns of the individual odours.

### Additional mushroom body connections

Our model suggests that only KC→MBON, MBON⊣DAN, MBON→DAN and DAN→MBON connections are essential for successful learning in the mushroom bodies. However, there are a number of additional known connections in the mushroom bodies, such as KC→APL, APL⊣KC, DAN→MBON, axoaxonic KC→KC and KC→DAN connections that have been neglected in this model, and need further consideration.

In the larval brain, there are two anterior paired lateral (APL) neurons, one for each mushroom body. They extend their dendrites to the lobes of the mushroom bodies and terminate their axons in the calyxes releasing the inhibitory GABA neurotransmitter (***Tanaka et al., 2008***). Although there are still two of them, in the adult brain both their dendrites and axons are innervating the calyx and the lobes (***Wu et al., 2013***), suggesting that they function as both global and local inhibitory circuits. Moreover, DAN⊣APL (***Liu and Davis, 2009***) and APL⊣DAN (***Wu et al., 2012***) connections have been proposed, but there is no clear description of what their function is. Several previous models (***Peng and Chittka, 2017***; ***Delahunt et al., 2018***) have demonstrated that a potential function for this global/local inhibition network is gain control such that the total number of KCs firing to different stimuli remains similar, and indeed that the same effect can be implemented using a flexible threshold for KC firing (***Saumweber et al., 2018***; ***Zhu et al., 2020***; ***Zhao et al., 2020***). In our model we have simplified the KC input, representing just two odours as different patterns across a small number of KCs with a fixed number of them being active at all times, so the hypothesised gain control function of the APL is not useful here. However it remains an interesting question whether there is learning between the KC and APL in the lobes (***Zhou et al., 2019***), or between the APL and KC in the calyx, and what role this might play in the overall dynamics of memory acquisition.

In addition, ***Eichler et al.*** (***2017***) suggest that most of the KCs input, i.e., 60%, is from other KCs. We suggest these connections (together with the ones from the APL) might create local winner-takes-all (WTA) networks that force a limited number of KCs per compartment to be active at one time. This predicts that it is possible for the same KC axon to be active in one compartment but inactive in another [consistent with recent data from ***Bilz et al.*** (***2020***)], and that an almost fixed number of KCs might be active at all times, even when no odour is delivered (e.g., fresh air only) enabling acquisition and forgetting at all times. ***Ito et al.*** (***2008***) shows that KCs can be active even in the absence of odours but with no consistent spiking, which is a characteristic of WTA networks when the underlying distribution of spikes across the neurons is almost uniform.

***Eichler et al.*** (***2017***) also observed (from electron microscopy reconstruction in larva) that within a compartment, in a ‘canonical microcircuit’, KCs make direct synapses to the axons of DANs, and that DAN pre-synapses often simultaneously contact KCs and MBONs. The same connections have been observed in adult Drosophila by (***Takemura et al., 2017***). The extent to which KCs (and thus odour inputs) might be directly exciting DANs remains unclear. ***Cervantes-Sandoval et al.*** (***2017***) show that stimulating KCs result in increased DAN responses and that DANs are activated through the ACh neurotransmitter. However, we note that in our model, such an effect could be explained without assuming a direct connection. For example, in the ‘long-term memory’ microcircuit, activating the KCs result in increased activity of the LTM MBON which excites the respective charging DAN. The DAN that ***Cervantes-Sandoval et al.*** (***2017***) provide evidence from is PPL1-α2α’2, which gets excited by MBON-α2α’2 neurons, as it is characterised by the ACh neurotransmitter (***Aso et al., 2014a***). In our terms, this could be an LTM MBON that excites its respective charging DAN, PPL1-α2α’2 (***Li et al., 2020***), and provide the source of ACh detected on it. More generally, the altered activity of DANs in response to odours that has been observed during learning can be also observed in our model, without requiring direct KC→DAN connections or their modification. Nevertheless, such connections may possibly play a role in enhancing the specificity of dopamine-induced changes in KC→MBON connectivity. Interestingly, the depression of KC→DAN synapses, in parallel with KC→MBON synapses, could provide an alternative mechanism for implementing RPE learning (***Takemura et al., 2017***).

***Takemura et al.*** (***2017***) demonstrate that the direct synapses observed from DANs to MBONs are functional in altering the MBON post-synaptic current to DAN activation, independently of KCs. This could be a mechanism by which learnt responses to reinforcements are coordinated with the current presence or absence of the reinforcement (***Schleyer et al., 2020***, ***2011***; ***Gerber and Hendel, 2006***). Another possibility is that post-synaptic as well as pre-synaptic changes might be involved in learning at the KC→MBON synapse (***Pribbenow et al., 2021***).

### Beyond attraction and aversion

The incentive circuit (IC) consists of 6 MBONs and 6 DANs that link a pair of antagonistic motivations, attraction and avoidance. However, there are ~ 34 MBONs and ~ 130 DANs in the mushroom body of the adult fruit fly brain, within which the IC is an identifiable motif. We suggest the possibility that this motif could be repeated, representing additional opposing motivations, with some neurons having multiple roles depending on the motivational context as proposed by ***Cohn et al.*** (***2015***), working either as restrained MBONs and discharging DANs, or as LTM MBONs and forgetting DANs depending on the reinforcer identity. We have illustrated this concept of a unified system of motivations as The incentive wheel. This could explain how PAM-β2β’2a [i.e., MB301B (***May et al., 2020***)] is a sugar-encoding discharging DAN in the appetitive olfactory conditioning context, but it also is an avoidance-driving forgetting DAN in a different context (e.g., aversive olfactory conditioning). In addition, 2 MBONs of the incentive circuit do not interact with the α’/β’ KCs of the mushroom body. MBON-γ4>γ1γ2 and MBON-γ1pedc>α/β, are part of two autonomous microcircuits, i.e., the susceptible memories (SM), and are working under the context provided by the ~ 675 γ-KCs relative to the task. This makes possible that the KCs from the γ lobe connect to all the susceptible memories of the flies for the ~ 8 available motivations illustrated in the The incentive wheel.

From a functional point of view, the mushroom bodies seem to be involved in the motivation and behaviour of the animal, especially when it comes to behaviours essential for survival. In the mammalian brain, this function is subserved by the limbic system, which is composed of a set of complicated structures, such as the thalamus, hypothalamus, hippocampus and amygdala (***Dalgleish, 2004***; ***Roxo et al., 2011***). According to ***Papez*** (***1937***), sensory (and mostly olfactory) input comes in the limbic system through the thalamus, which connects to both the cingulate cortex (through the sensory cortex) and the hypothalamus (***Roxo et al., 2011***; ***Dalgleish, 2004***). Responses in the cingulate cortex are guiding the emotions, while the ones in the hypothalamus are guiding the behaviour (bodily responses). Finally, the hypothalamus connects with the cingulate cortex through the anterior thalamus (forward) and the hippocampus (backward stream). ***MacLean*** (***1949***) augmented this model, by adding the amygdala and PFC structures that encode primitive emotions (e.g., anger and fear) and connect to the hypothalamus (***Roxo et al., 2011***; ***Dalgleish, 2004***). We suggest that some of the functions we have identified in the MB incentive circuit could be mapped to limbic system structures (see ***Figure 13***).

**Figure 13.**
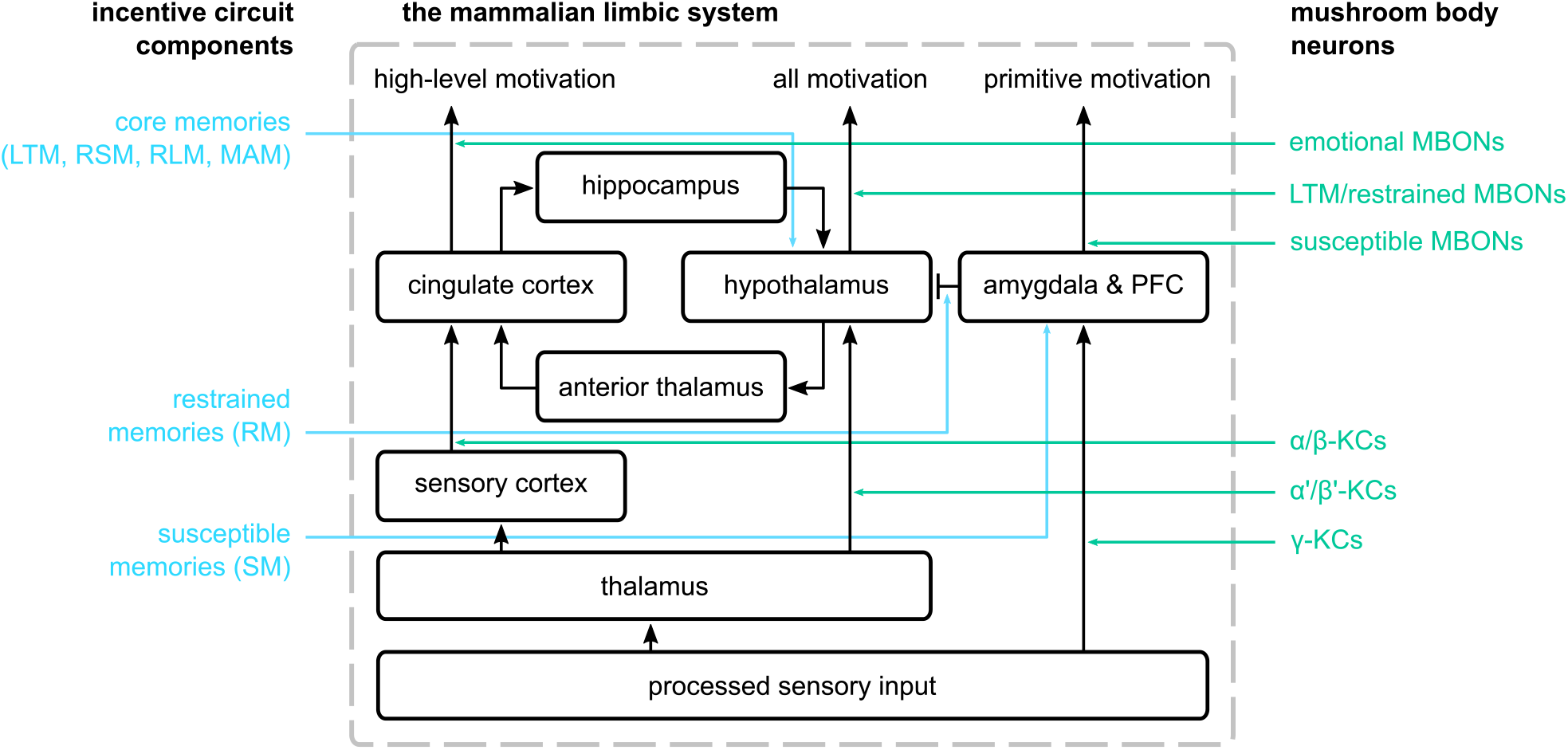
The mammalian limbic system as described by ***Papez*** (***1937***) and ***MacLean*** (***1949***) and the suggested parallels in the proposed incentive circuit. On the left, we show the mushroom body microcircuits that correspond to the different structures in the mammalian limbic system. In the centre, we have the connections among the different structures of the limbic system. On the right, we show the groups of mushroom body neurons we suggest that have similar function to the ones in the limbic system.

More specifically, the α’/β’-KCs could have similar role to the neurons in the thalamus, α/β-KCs represent a higher abstraction of the input stimuli and have a similar role to the ones in the sensory cortex, while the γ-KCs represent relatively unprocessed stimuli. This would make the susceptible MBONs a parallel to neurons in the amygdala, creating responses related to primitive motivations and connecting to (inhibiting) the restrained MBONs, which we would compare to the hypothalamus as providing the main control of behaviour. As we suggest that the same MBONs could fulfil a role as LTM or restrained in different circuits (see The incentive wheel), the LTM would also correspond to hypothalamus, with input from the α’/β’-KCs, and thus the RSM, RLM, LTM and MAM microcircuits are assumed to correspond to hypothalamus functions. Following this analogy we predict that the function of the cingulate cortex then is represented by the α/β MBONs, encoding the ‘emotions’ of the animal towards reinforced stimuli, potentially controlling more sophisticated decision making. This mapping would suggest the connections amongst the restrained/LTM (α’/β’) MBONs and the ‘emotional’ (α/β) MBONs are similar to the hippocampus and anterior thalamus pathways.

#### Box 1. Summary of predictions.

The model yields predictions that can be tested using established experimental protocols:

1. MBON-γ2α’1 and MBON-γ5β’2a should exhibit short-term memories, while MBON-α’1 and MBON-β2β’2a long-term memories. MBON-γ1pedc>α/β and MBON-γ4>γ1γ2 should exhibit susceptible memories. Restrained and susceptible MBONs should show more consistent responses across flies. LTM MBONs should have more variable responses because they encode all previous experiences of the animal.
2. Activating MBON-γ2α’1 or MBON-β2β’2a should increase the responses rate of PAM-β’2a, and similarly activating MBON-γ2β’2a or MBON-α’1 should excite PPL1-γ2α’1. This would verify the excitatory STM reciprocal and LTM feedback connections of the circuit. By activating the LTM MBONs (e.g., MBON-α’1 and MBON-β2β’2a) should also excite the forgetting DANs (e.g., PAM-β2β’2a and PPL1-γ2α’1 respectively). This would verify the excitatory LTM reciprocal connections of the circuit.
3. By consistently activating one of the LTM MBONs while delivering a specific odour, the LTM MBON should show increased response to that odour even without the use of a reinforcement. This would verify the saturation effect of the DPR and the charging momentum hypothesis. On the other hand, if we observe reduced response rate, this would show that MBON-DAN feedback connection is inhibitory and that RPE is implemented by the circuit.
4. Blocking the output of charging DANs (i.e., PPL1-γ2α’1 and PAM-β’2a) could reduce the acquisition rate of LTM MBONs, while blocking the output of LTM MBONs would prevent memory consolidation. Blocking the reciprocal connections of the circuit should prevent generalising amongst opposing motivations (unable to make short- or long-term alteration of responses to odours once memories have formed). Blocking the output of forgetting DANs would additionally lead to hyper-saturation of LTMs, which could cause inflexible behaviour.
5. Activation of the forgetting DANs should depress the KC-MBON synaptic weights of the restrained and LTM MBONs of the same and opposite valence respectively, and as a result suppress their response to KC activation. Activation of the same DANs should cause increased activity of these MBONs for silenced KCs at the time.
6. Unpaired conditioning should involve the LTM circuit (or at least some microcircuit within the MB where the MBON excites a DAN). Second-order conditioning should involve the LTM circuit, and might not require the susceptible and restrained memory circuits. Backward conditioning might not occur in all compartments as in our model it required that the target MBON inhibits its respective DAN (susceptible memory microcircuit) and to date has only been demonstrated for microcircuits with this property.
7. DANs that innervate more than one compartment may have different functional roles in each compartment.

While it might seem startling to suggest that a compact circuit of single identified neurons in the insect MB mimics in miniature these far larger and more complex structures in the mammalian brain, the justification comes from the similarity in the behavioural demands common to all animals: surviving and adapting in a changing world.

## Methods

### Implementation of the incentive circuit

We represent the connections between neurons by using synaptic-weight matrices and non-linearly transform the information passing from one neuron to another by using an activation function. Next, we define these parameters and some properties of our computational model, which are not a result of unconstrained optimisation and are consistent throughout all our experiments.

#### Parameters of the model

We assume that the odour identity passes through the projection neurons (PNs) into the mushroom body and its Kenyon cells (KCs). It is not in the scope of this work to create a realistic encoding of the odour in the PNs, so we assume that the odour signal is represented by *n*_p_ = 2 PNs, one for each odour, and that these project to form distinct activations in a set of *n*_k_ = 10 KCs in the MB, i.e, subset of KCs that respond to the specific odours used in the experiments. Therefore, the vector **p**_A_ = [1, 0] represents the activity of the PNs when odour A is detected, **p**_B_ = [0, 1] when odour B, **p**_AB_ = [1, 1] when both odours and **p**_Ø_ = [0, 0] when none of them is detected. The responses of the KCs are calculated by

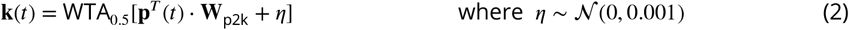

*η* is some Gaussian noise, 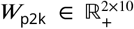 is the weights matrix that allows the transformation of the 2-dimensional odour signal into the 10-dimensional KC responses, and *t* is the current time-step. The WTA_0.5_[*x*] is an activation function that keeps the top 50% of KCs active, based on the strength of their activity. Note that the number of neurons we are using for PNs and KCs is not very important and we could use any combination of PN and KC populations. However, the bigger the KC population the smaller percentage of them should be active. The PN→KC synaptic weights used are shown below

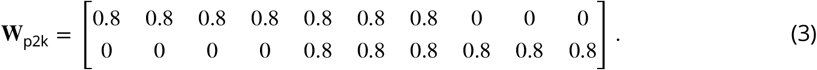

The odours are represented by different firing patterns across 10 KCs: 4 fire only for A, and 3 fire only for B, while the remaining 3 fire to either odour. This is to show the effects of the DPR when we have overlap in the KCs that respond to the two odours used in the conditioning paradigm. This assumption also created the best fit with the data, suggesting that there might be overlapping KCs encoding the real odours tested in the fly experiments.

We transform the reinforcement (US), **u**(*t*) ∈ {0, 1}^2^, delivery into an input for the DANs by using the weights matrix 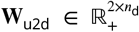. We represent the activity of the DANs, 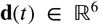, with a 6-dimensional vector, where each dimension represents a different neuron in our model. Specifically,

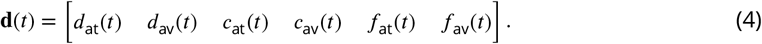

The US is represented by a 2-dimensional vector where the 1^st^ dimension denotes rewarding signal and the 2^nd^ dimension denotes punishment: **u**_sugar_ = [1, 0] and **u**_shock_ = [0, 1]; and the contribution of this vector to the responses of the DANs is is given by

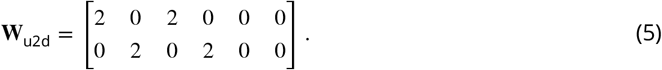

In line with the DANs vector representation, we have a similar vector for MBONs, 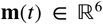, where each dimension represents the response of a specific neuron in time *t* as is shown in the equation below

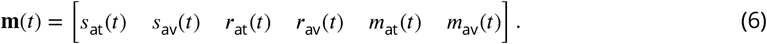

The weight matrix that encodes the contribution of KCs to the MBON responses, 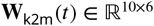, is initialised as

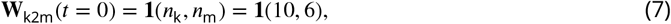

which effectively is a 10 × 6 matrix of ones. In other words, all KCs connect to all MBONs, and their initial weight is positive and the same for all connections. As these are plastic weights, their value depends on the time-step, and therefore we provide time, *t*, as a parameter. Note that also *w*_rest_ = 1, which initially results in the absence of memory, 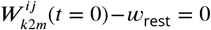. Thus, any deviation of the synaptic weights from their resting value represents a stored memory with strength 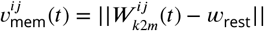.

There are also MBON→DAN, 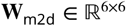, and MBON→MBON connections, 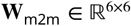, which are given by

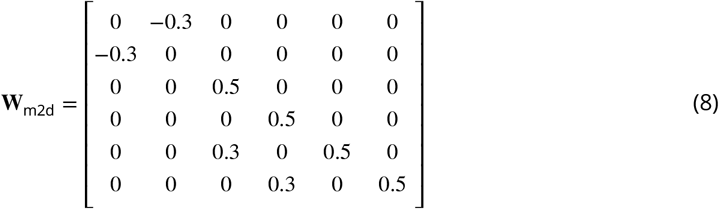

and

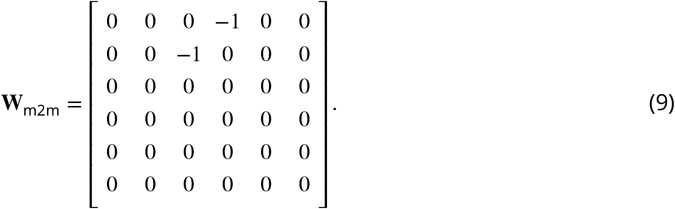

The above matrices summarise the excitatory (positive) and inhibitory (negative) connections between MBONs and DANs or other MBONs as defined in the incentive circuit (***Figure 3***, see also ***Figure 14***). The sign of the weights was fixed but the magnitude of the weights was hand-tuned in order to get the desired result, given the constraint that equivalent types of connections should be same weight (e.g., in the reciprocal microcircuits). The magnitude of the synaptic weights specify the effective strength of each of the described microcircuits in the overall circuit. We also add some bias to the responses of DANs, **b**_d_, and MBONs, **b**_m_, which is fixed as

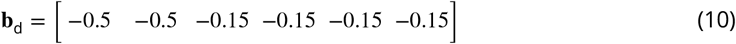

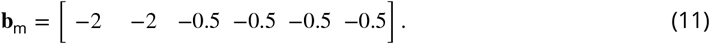

**Figure 14.**
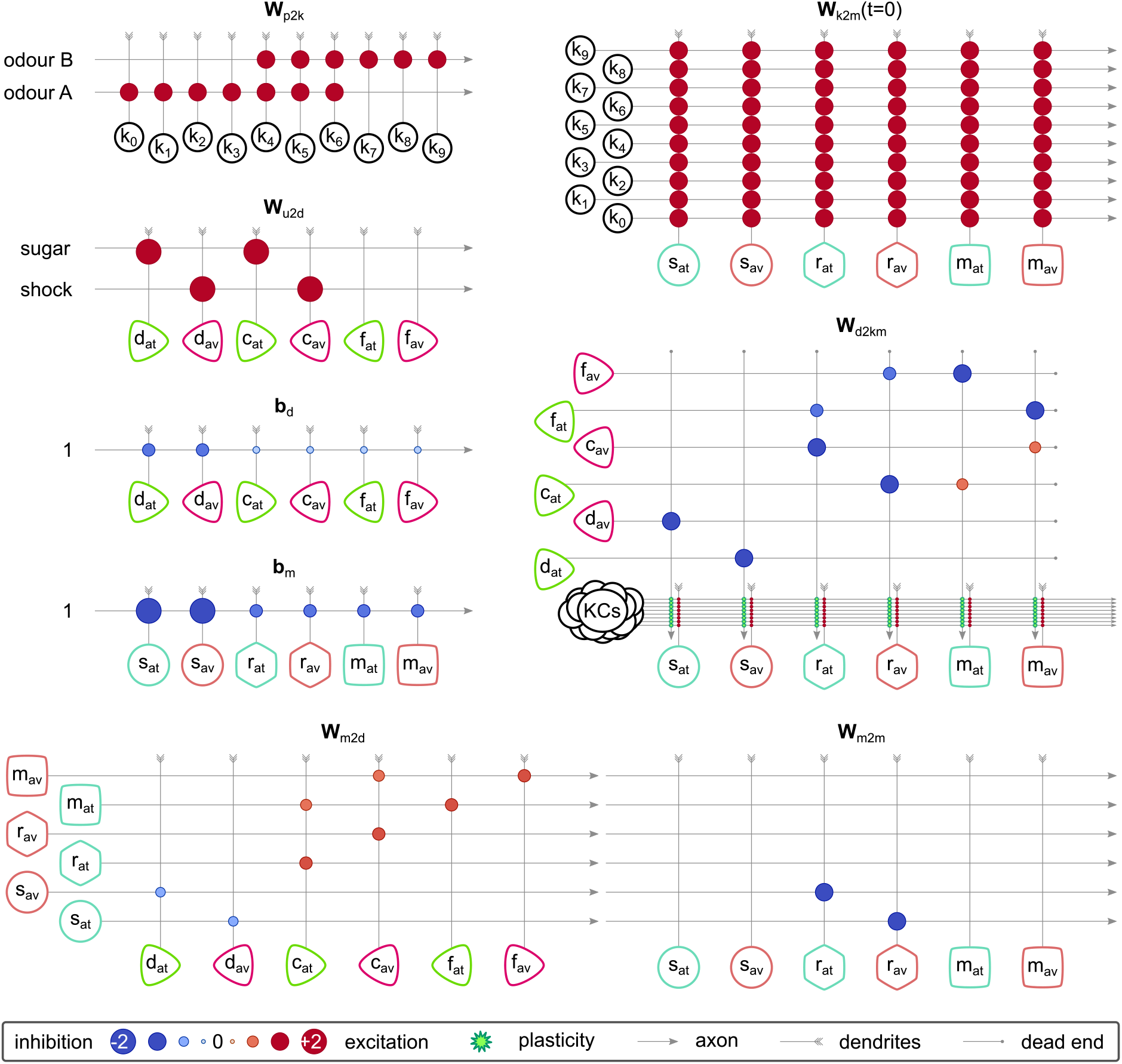
The synaptic weights and connections among the neurons of the incentive circuit (IC). Each panel corresponds to a different synaptic-weights-matrix of the circuit. The size of the circles when the pre-synaptic axon crosses the post-synaptic dendrite shows how strong a connection is, and the colour shows the sign (blue for inhibition, red for excitation). Light-green stars show where the synaptic plasticity takes place and how the DANs modulate the synaptic weights between KCs and MBONs. **Figure 14–Figure supplement 1.** The responses of the DANs and MBONs of the circuit when altering the pre-synaptic strengths of MBONs. **Figure 14–Figure supplement 2.** The responses of the DANs and MBONs of the circuit when altering the DA modulation strengths. **Figure 14–Figure supplement 3.** The responses of the DANs and MBONs of the circuit when altering the DAN and MBON biases.

This bias can be interpreted as the resting value of the neurons, or some external input from other neurons that are not included in our model.

Finally, we define the DAN-function matrix, 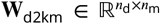, which transforms the responses of the DANs into the dopamine factor that modulates the **W**_k2m_(*t*) synaptic weights, and it is given below

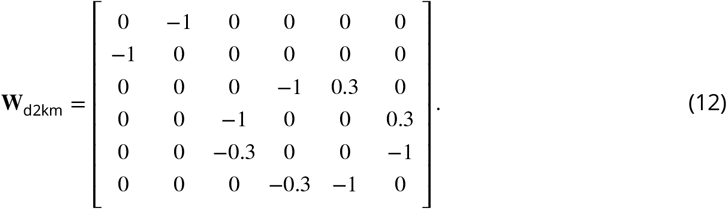

All the parameters described above are illustrated in ***Figure 14***. ***Figure 14***–***Figure Supplement 1***, ***Figure 14***–***Figure Supplement 2*** and ***Figure 14***–***Figure Supplement 3*** show how each of these parameters affect the responses of the neurons in the incentive circuit. The last thing left to describe is the activation function, which is used in order to generate the DAN and MBON responses. This is

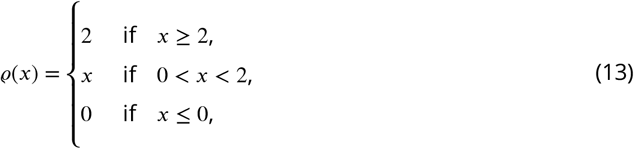

which is the *rectified linear unit* (ReLU) function, bounded in *ϱ*(*x*) ∈ [0, 2]. The reason why we bound the activity is in order to avoid having extremely high values that explode during the charging of the LTM.

#### Forward propagation

For each time-step, *t*, we read the environment and propagate the information through the model in order to update the responses of the neurons and the synaptic weights. This process is called *forward propagation* and we repeat it as long as the experiment runs.

First, we read the CS, **p**(*t*), and US, **u**(*t*), from the environment and we calculate the KC responses by using ***Equation 2***. In order to calculate the DANs and MBONs update we define the differential equations as follows

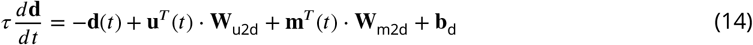

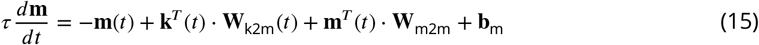

where *τ* = 3 is a time-constant that is defined by the number of time-steps associated in each trial and *T* denotes the transpose operation of the matrix or vector. Using the above differential equations, we calculate the updated responses (i.e., responses in the next time-step, *t* + 1) as

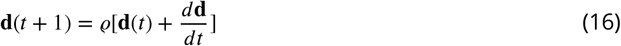

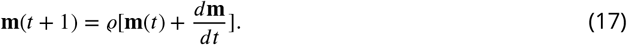

Finally, we calculate the dopaminergic factor, 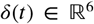, and update the KC→MBON synaptic weights as

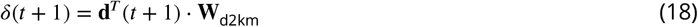

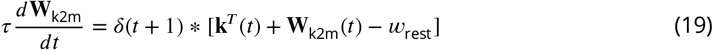

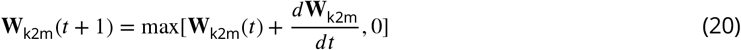

where denotes the element-wise multiplication and *w*_rest_ = 1 is the resting value of the weights. Note that element-wise multiplication means that each element of the *δ*(*t*) vector will be multiplied with each column of the **W**_k2m_(*t*) matrix. Also, the element-wise addition of the transposed vector, **k**^*T*^(*t*), to the **W**_k2m_(*t*) matrix, means that we add each element of **k**(*t*) to the corresponding row of **W**_k2m_(*t*). We repeat the above procedure as many times as it is required in order to complete the running experimental paradigm routine.

#### Modelling the neural responses

To emulate the acquisition and forgetting paradigms used for flies, we run the simulated circuit in an experiment that consists of *T* = 73 time-steps. Each time-step actually comprises four repeats of the forward propagation update described above, to smooth out any bias due to the order of computations (value vs weights update). After the initialisation time-step at *t* = 0 there are 24 *trials* where each trial consists of 3 *in-trial time-steps*.

Within each trial, the first time-step has no odour, and in the second and third time steps, odour is presented: odour Aon even trials and odour B on odd trials. Atrial can have no shock (***Figure 15***A), unpaired shock presented in the first time step (***Figure 15***B), or paired shock presented in the third time step (***Figure 15***C). The first 2 trials compose the ‘pre-training phase’, where we expose the model to the two odours alternately (i.e., odour A in trial 1 and odour B in trial 2) without shock delivery. Then we have the acquisition phase, where we deliver shock paired with odour B for 10 trials (5 trials per odour; ***Figure 15***D). Before we proceed to the forgetting phases, we leave 2 empty trials (1 per odour), which we call the *resting trials*. The forgetting phases last for another 10 trials (5 trials per odour; ***Figure 15***E, F and G). During the extinction phase, no shock is delivered while we continue alternating the odours (see ***Figure 15***E); during the unpaired phase, shock is delivered unpaired from odour A (see ***Figure 15***F); while at the reversal phase shock is paired with odour A (***Figure 15***G).

**Figure 15.**
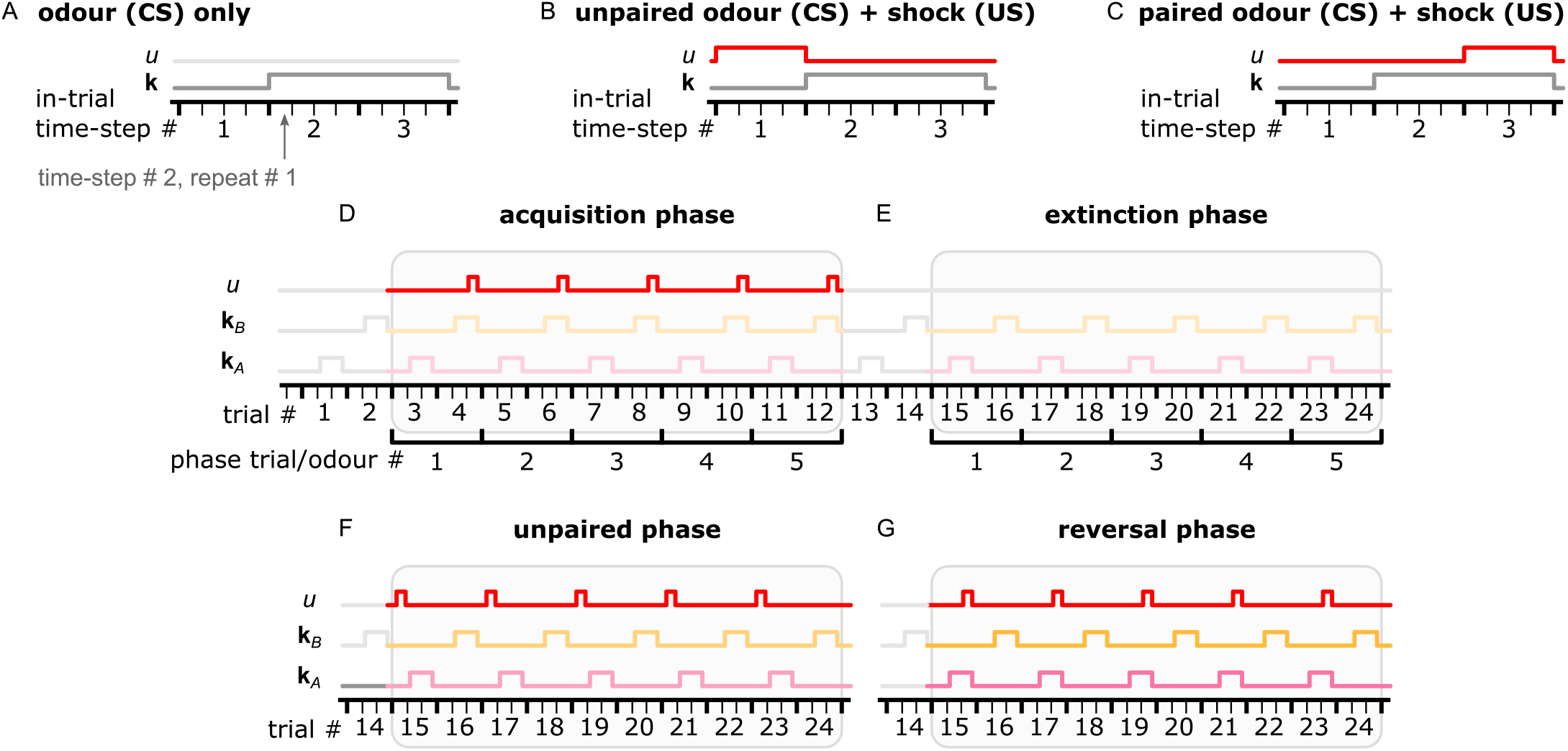
Description of the simulation process from our experiments. A single trial is composed of 3 in-trial time-steps and each time-step by 4 repeats. Odour is provided only during the 2^nd^ and 3^rd^ in-trial time-steps, while shock delivery is optional. (A) In an extinction trial only odour (CS) is delivered. (B) During an unpaired trial, shock is delivered during in-trail time-step 1. (C) During a paired trial, shock is delivered along with odour delivery and during in-trial time-step 3. (D) The acquisition phase has 5 odour A only trials and 5 paired odour B trials alternating. (E) The extinction phase has 5 odour A only trials and 5 odour B only trials alternating. (F) The unpaired phase has 5 odour A unpaired trials and 5 odour B only trials alternating. (G) The reversal phase has 5 odour A paired trials and 5 odour B only trials alternating. The colours used in this figure match the ones in ***Figure 4***.

#### The classic unpaired conditioning paradigm

In this case, during the acquisition phase we deliver only electric shock in odd trials (omission of odour B), followed by an extinction phase as described above.

#### Modelling the behaviour

The experiments last for 100 sec each and they are split in 3 phases as shown in ***Figure 11***A. In *pre-training*, the flies are placed in the centre of arena and explore freely for 20 sec. In *training*, either shock or sugar is associated with the region 30cm around odour A, odour B, or around both sources for 30 sec. In *post-training*, we remove the reinforcement and let the flies express their learnt behaviour for another 50 sec creating an extinction forgetting condition. ***Figure 11***B shows the normalised cumulative time spent experiencing each odour and the odour preference of the flies during the different phases for each of the six training conditions, and for 10 repeats of the experiment, when their behaviour is controlled by a combination of the attractive and repulsive forces on the two odours. The actual paths of the flies for all the 10 repeats are illustrated in ***Figure 11***–***Figure Supplement 4***.

In practice, in order to create the experiences of *n*_fly_ = 100 flies, we have created another routine that embeds the simulation of their motion and environment. We represent the position of each fly, 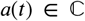, and the sources of the odours in the arena, 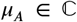 and 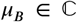 for odours A and B respectively, in the 2D space as complex numbers in the form *x* + *iy*. Therefore, the flies are initialised in *a*(*t* = 0) = 0 and the sources of the odours are placed in *μ_A_* = −0.6 and *μ_B_* = 0.6. The standard deviation of the odour distributions are *σ_A_* = *σ_B_* = 0.3.

We get the odour intensity in each time-step by using the Gaussian density functions of the two odours and the position of the fly in the arena

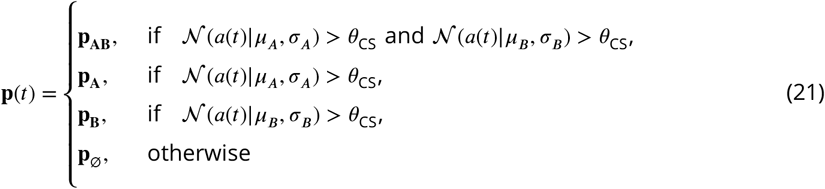

where **p**_*A*_, **p**_*B*_, **p**_*AB*_ and **p**_Ø_ are the identities of odours A, B, “A and B” and none of them respectively in the PNs as described in Parameters of the model, and *θ*_CS_ = 0.2 is the detection threshold forthe odour. Note that PN responses depend only on the fact that an odour has been detected or not and it is not proportional to the detected intensity. The reinforcement is applied to the simulated fly when the position of the agent is inside a predefined area around the odour, i.e., ∥*μ*_CS_ − *a*(*t*)∥ < *ρ*_US_, where *ρ*_US_ = 0.3 is the radius of the reinforced area. Note that the radius of the area where the odour is detectable is roughly *ρ*_CS_ ≃ 0.58, which is larger than the reinforced area. Then we run a Forward propagation using the above inputs.

From the updated responses of the MBONs, we calculate the *attraction force*, **v**(*t*), for the mixture of odours which modulates the velocity of the fly. This force is calculated by taking the difference between the responses of the MBONs that drive the behaviour:

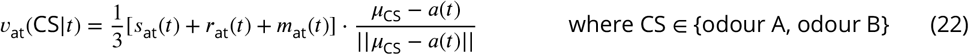

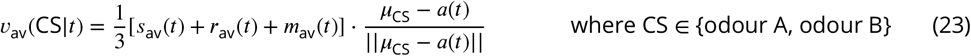

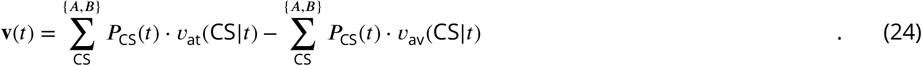

where *μ*_CS_ is the position of the odour source and *P*_CS_(*t*) is the probability of being closer to the specific CS source calculated using the Gaussian distribution function and the Bayesian theorem. For example, given that the prior probability of being closer to odour A and odour B is equal at any time, i.e., *P*(*A*) = *P*(*B*) = 0.5, the probability of being closer to odour A is given by

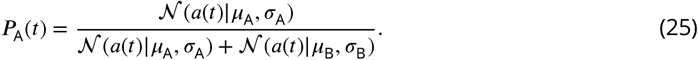

The velocity of the simulated fly is updated as follows

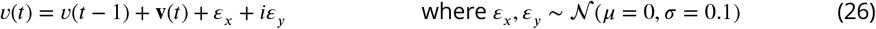

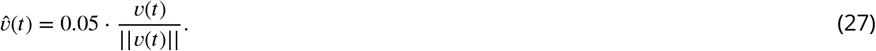

We normalise the velocity so that we keep the direction but replace the step size with 0.05 m/sec. The noise added to the velocity is introduced in order to enable the flies to move in 2 dimensions and not just between the two odour sources. Also, when the attraction force is **v**(*t*) = 0, then the noise along with the previous velocity is the one that drives the flies.

We repeat the above process for *T* = 100 time-steps with 1 Hz (1 time-step per second), and we provide shock or sugar (when appropriate) between time-steps 20 and 50, otherwise we use a zero-vector as US input to the DANs.

#### Calculating the normalised cumulative exposure and the preference Index

In ***Figure 11***B, for each phase (i.e., pre-training, training and post-training) we report the normalised cumulative exposure of the flies in each odour and their preference index between them. The normalised cumulative exposure is calculated by

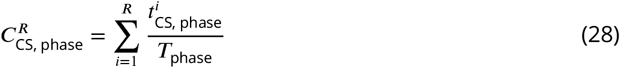

where *R* is the repeat of the experiment, *i* is the iterative repeat, *T*_phase_ is the number of time-steps for the specific phase, 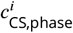, is the number of time-steps spend exposed in the specific CS ∈ {A, B}, phase and repeat.

The preferences index for every repeat is calculated using the above quantities

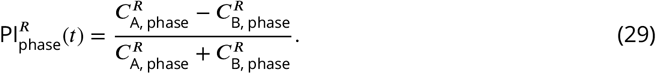

#### The reward prediction error plasticity rule

In ***Figure 9***–***Figure Supplement 2***, ***Figure 9***–***Figure Supplement 3***, ***Figure 11***–***Figure Supplement 5*** and ***Figure 11***–***Figure Supplement 6*** we present the responses and synaptic weights of the incentive circuit neurons, and the behaviour of the simulated flies using the RPE plasticity rule. This was done by replacing our plasticity rule in ***Equation 19*** with the one below

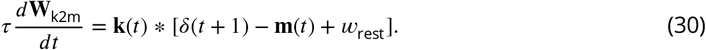

### Derivation of the dopaminergic plasticity rule

***Handler et al.*** (***2019***) suggest that ER-Ca^2+^ and cAMP playa decisive role in the dynamics of forward and backward conditioning. More specifically, they suggest that the KC→MBON synaptic change, 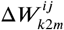 is proportional to the combined ER-Ca^2+^ and cAMP levels, which can be written formally as

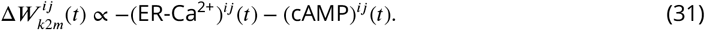

We assume that ER-Ca^2+^ and cAMP levels are determined by information available in the local area of the target KC axon (pre-synaptic terminal): the dopamine (DA) level emitted by the DANs to the KC synapses of the respective (*j*^th^) MBON, *D^j^*(*t*) ≥ 0; the activity of the (*i*^th^) pre-synaptic KC, *k^i^*(*t*) ≥ 0; the respective KC→MBON synaptic weight; 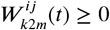 (assumed always positive, exciting the MBON) and the resting synaptic weights, 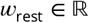, which we assume are a constant parameter of the synapse. Tuning the above quantities in order to reproduce the ER-Ca^2+^ and cAMP levels, we postulate a mathematical formulation of the latter as a function of the available information

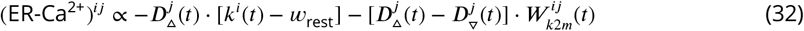

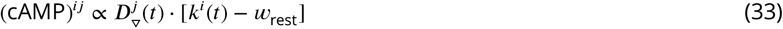

where 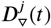 and 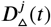 are the depression and potentiation components of the DA respectively [assumed to correspond to DopR1 and DopR2 receptors (***Handler et al., 2019***), or potentially to involve co-transmitters released by the DAN such as Nitric Oxide (***Aso et al., 2019***)]. We assume two types of DAN terminals: the depressing and potentiating terminals. In depressing terminals (arrow down), 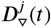 makes a higher peak in its activity followed by a faster diffusion than 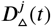, which seems to be the key for the backward conditioning. The opposite happens in potentiating DAN terminals. ***Figure 16*** shows the ER-Ca^2+^ and cAMP levels during forward and backward conditioning for a depressing DAN [see ***Figure 16***–***Figure Supplement 1*** for the responses of all the terms used including 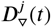 and 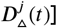, which are comparable to the data shown in ***Handler et al.*** (***2019***) (also ***Figure 16*** — shown in grey). Note that here we are more interested in the overall effects of learning shown in ***Figure 16***A rather than the detailed responses of ***Figure 16***B.

**Figure 16.**
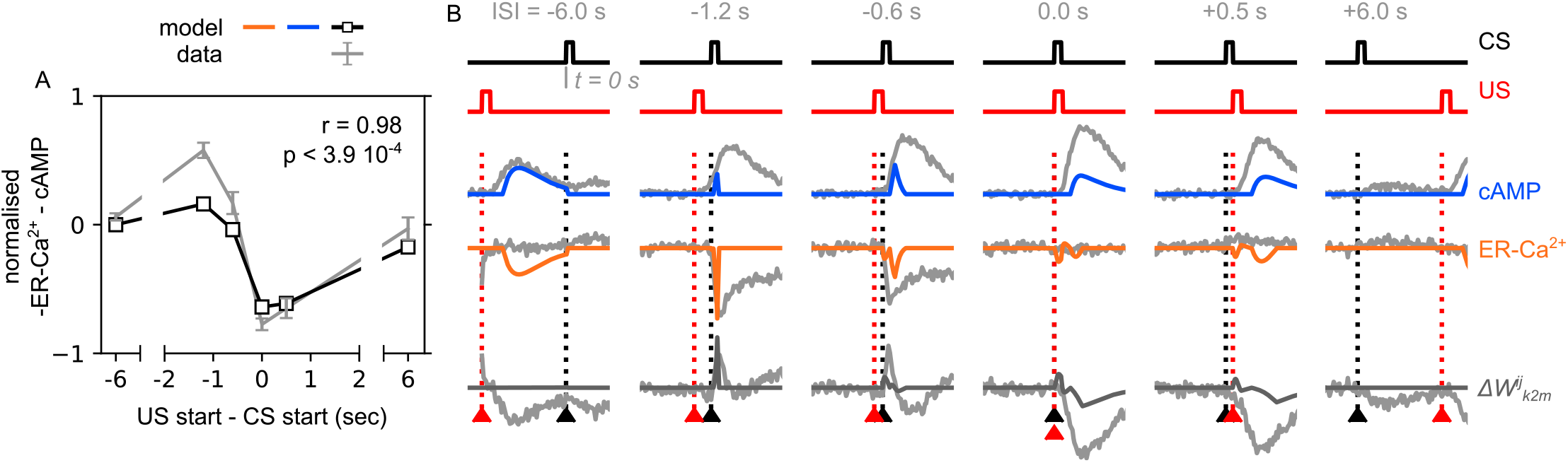
The effect of the ER-Ca^2+^ and cAMP based on the order of the conditional (CS) and unconditional stimuli (US). (A) Normalised mean change of the synaptic weight plotted as a function of the Δ*s* (US start - CS start), similar to ***Handler et al.*** (***2019***) - ***Figure 5***F (blue line). For ease of comparison, the predicted mean values are drawn on the top of the data (mean ±SEM) from the original paper (***Handler et al., 2019***) - grey lines and error-bars. (B) Detailed ER-Ca^2+^ and cAMP responses reproduced for the different Δ*s*, and their result synaptic-weight change. Black arrowhead marks time of the CS (duration 0.5 sec); red arrowhead marks time of the US (duration 0.6 sec), similar to ***Handler et al.*** (***2019***) - ***Figure 5***D. For ease of comparison, the predicted responses are drawn on the top of the data from the original paper (***Handler et al., 2019***) - grey lines. **Figure 16–Figure supplement 1**. All the chemical levels and neural activities calculated based on the order of the conditional (CS) and unconditional stimuli (US). **Figure 16–Figure supplement 2**. Parameter exploration of the *τ*_short_ and *τ*_long_ in ***Equation 35*** and ***Equation 36***.

By replacing ***Equation 32*** and ***Equation 33*** in ***Equation 31***, we can rewrite the update rule as a function of known quantities, forming our *dopaminergic plasticity rule* (DPR) of ***Equation 1***, which we rewrite of convenience

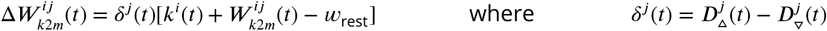

The *dopaminergic factor*, *δ^j^*(*t*), is the difference between the 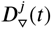 and 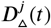 levels, and it can be positive 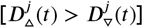 or negative 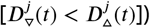. Combined with the state of the KC activity results in the four different weight modulation effects: *depression, potentiation, recovery* and *saturation*.

In ***Figure 16***B (where we assume a depressing DAN terminal), all four effects occur in four out of the six cases creating complicated dynamics that allow forward and backward learning. Similarly, a potentiating terminal might trigger all the effects in a row but in different order and duration. Note that in the simulations run for the results of this paper, we simplify the dopaminergic factor to have a net positive or negative value for the time-step in which it influences the synaptic weight change, as the time-steps used are long enough (e.g., around 5 sec; see Implementation of the incentive circuit) and we assume less complicated interchange among the effects.

In ***Figure 16***A, we report the normalised mean change of the synaptic weight calculated using the computed ER-Ca^2+^ and cAMP levels and the formula below

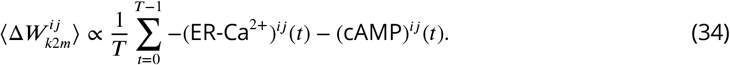

#### Decomposing the dopaminergic factor

In ***Equation 18***, the dopaminergic factor, *δ*(*t*) is derived from the matrix ***Equation 12*** which captures in abstracted and time-independent form the effects of dopamine release. To model these more explicitly, as described in Derivation of the dopaminergic plasticity rule, the dopaminergic factor can be decomposed as *δ*(*t*) =**D**_▵_(*t*)−**D**_▿_(*t*) where each component has a time dependent form given by the differential equations:

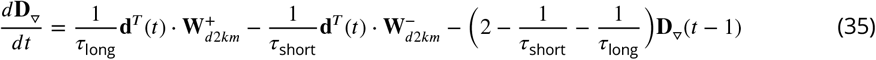

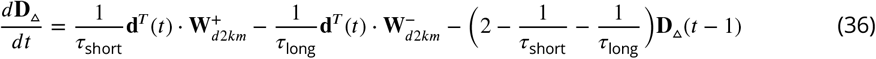

where 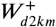 and 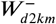 represent the positive-only (potentiation/saturation) and negative-only (depression/recovery) dopaminergic effects; *τ*_short_ and *τ*_long_ are time-constants that define the short (main) and long (secondary) duration of the dopamine effect. The longer the time-constant, the slower the diffusion but also the lower the peak of the effect. Note the two time-constants must satisfy the constraint 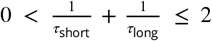 in order for the above differential equations to work properly.

In ***Figure 16***, where we are interested in more detailed dynamics of the plasticity rule, and the sampling frequency is high, i.e., 100Hz, we use *τ*_short_ = 60 and *τ*_long_ = 104, which we choose after a parameter exploration available in ***Figure 16***–***Figure Supplement 2***. This essentially means that **D**_▿_(*t*) and **D**_▵_(*t*) are expressed as time-varying functions following DAN spike activity. Note that for the specific (susceptible) type of MBON examined there, the DAN causes depression of the synapse, so there is no positive dopaminergic effect, i.e., 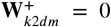. By setting 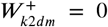 in equations ***Equation 35*** and ***Equation 36***, we have the fast update with the high peak for **D**_▿_(*t*) (0.5 sec for a full update) and a slower update with lower peak for **D**_▵_(*t*) (1 sec for a full update), as described in the Derivation of the dopaminergic plasticity rule.

For the experiments in ***Figure 4*** and ***Figure 11***, we use *τ*_short_ = 1 and *τ*_long_ = +∞, which removes the dynamics induced by the relation between **D**_▿_(*t*) and **D**_▵_(*t*), and ***Equation 18*** emerges from

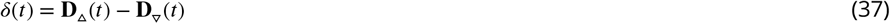

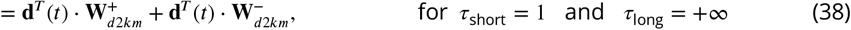

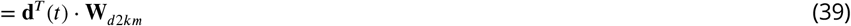

This essentially means that each update represents a time-step that is longer than the effective period of backward conditioning for the responses of the Microcircuits of the mushroom body and Modelling the behaviour sections (where sampling frequency is low, i.e., ≤ 0.5Hz and 1 Hz respectively) and therefore we use the same time-constants that result in the simplified ***Equation 18***.

### Data collection

In order to verify the plausibility of the incentive circuit, we recorded the neural activity in genetically targeted neurons during aversive olfactory conditioning which is described in more detail in ***McCurdy et al.*** (***2021***). We simultaneously expressed the green GCaMP_6_f Ca^2+^ indicator and red Ca^2+^-insensitive tdTomato in neurons of interest to visualise the Ca^2+^ changes which reflect the neural activity. We collected data from 357 5- to 8-days-old female flies (2-14 per neuron; 8 flies on average) and for 43 neurons, which can be found in ***Figure 4***–***source data 1*** (also illustrated in ***Figure 4***–***Figure Supplement 1***).

Each fly was head-fixed for simultaneous delivery of odours and electric shock while recording the neural activity. Their proboscis was also glued, while their body and legs were free to move (see ***Figure 4***A). The flies were allowed to recover from the gluing process for 15 min before placing them under the microscope. We used green (555 nm) and blue (470 nm) lights to record GCaMP and Tomato signals. We also used 0.1% 3-octanol (OCT) and 0.1% 4-methylcyclohexanol (MCH) for odour A and B respectively, and the flow rate was kept constant at 500 mL/min for each odour. The flies were allowed to acclimate to the airflow for at least one minute before starting of the experiment.

During the experiments, we alternate trials where 5 sec of each odour is presented 5 sec after the (green or red) light is on. We start with 2 pre-training trials (1 per odours) followed by 5 acquisition trials per odour. During acquisition, flies receive alternating 5 sec pulses of OCT (odour A) and MCH (odour B) paired with electric shock, repeated for 5 trials. During reversal, OCT is presented with shock and MCH without, repeated for 2 trials. On trials where electric shock was delivered, it was presented 4 sec after odour onset for 100 ms at 120V.

### Calculating off- and on-shock values

From the data collection process described above we get trials of 100 time-steps and at 5 Hz (20 sec each). Odour is delivered between time-steps 25 and 50 (between 5 sec and 10 sec), and shock is delivered during time-step 45 (at 9 sec). In this work, we report 2 values for each trial: the *off-shock* and *on-shock* values, which represent the average response to the odour before and during the period in which shock delivery could have occurred (even if shock is not delivered).

For the off-shock value, from each data-stream of activity from the target neuron, we collect the values from time-steps between 28 (5.6 sec) and 42 (8.4 sec). This gives us a matrix of *n*_fly_ × 15 values, whose average and standard deviation is the reported off-shock value. Similarly, for the on-shock values, we collect the values in time-steps between 44 (8.6 sec) and 48 (9.6 sec), which gives a matrix of *n*_fly_ ×5 values, whose average and standard deviation is the on-shock value. We define ‘on-shock’ as the time window from 8.6 sec to 9.6 sec, where shock onset occurs at *t* = 9 sec.

## Acknowledgements

We are grateful to Bertram Gerber for his useful comments on earlier drafts of the manuscript. We also thank James Bennett for discussion on their data and experiments and Vanessa Ruta for kindly providing their data for validating the dopaminergic plasticity rule. We also thank the Insect Robotics group for helpful critique on the figures and the reviewers on earlier revisions for their fruitful comments.

## Appendix 1 The incentive wheel

We have shown that the incentive circuit (IC) is able to explain classical conditioning experiments that have been done with adult fruit flies, and its neurons can replicate the responses of the mushroom bodies in the flies’ brain. We have also seen that three types of memories are stored in this model (i.e., susceptible, restrained and long-term) for each of the two represented motivations of the animal (i.e., attraction or avoidance) guided by reinforcements (i.e., reward or punishment). Although this model is sufficient to explain the behaviour of the animals in the laboratory, where the animal is exposed in controlled portions of chemicals and the results are translated into a simple attraction to or avoidance from a source, in the wild there are more than two motivations that modulate the behaviour of the animal either synergistically or opponently.

**Appendix 1 Figure 1.**
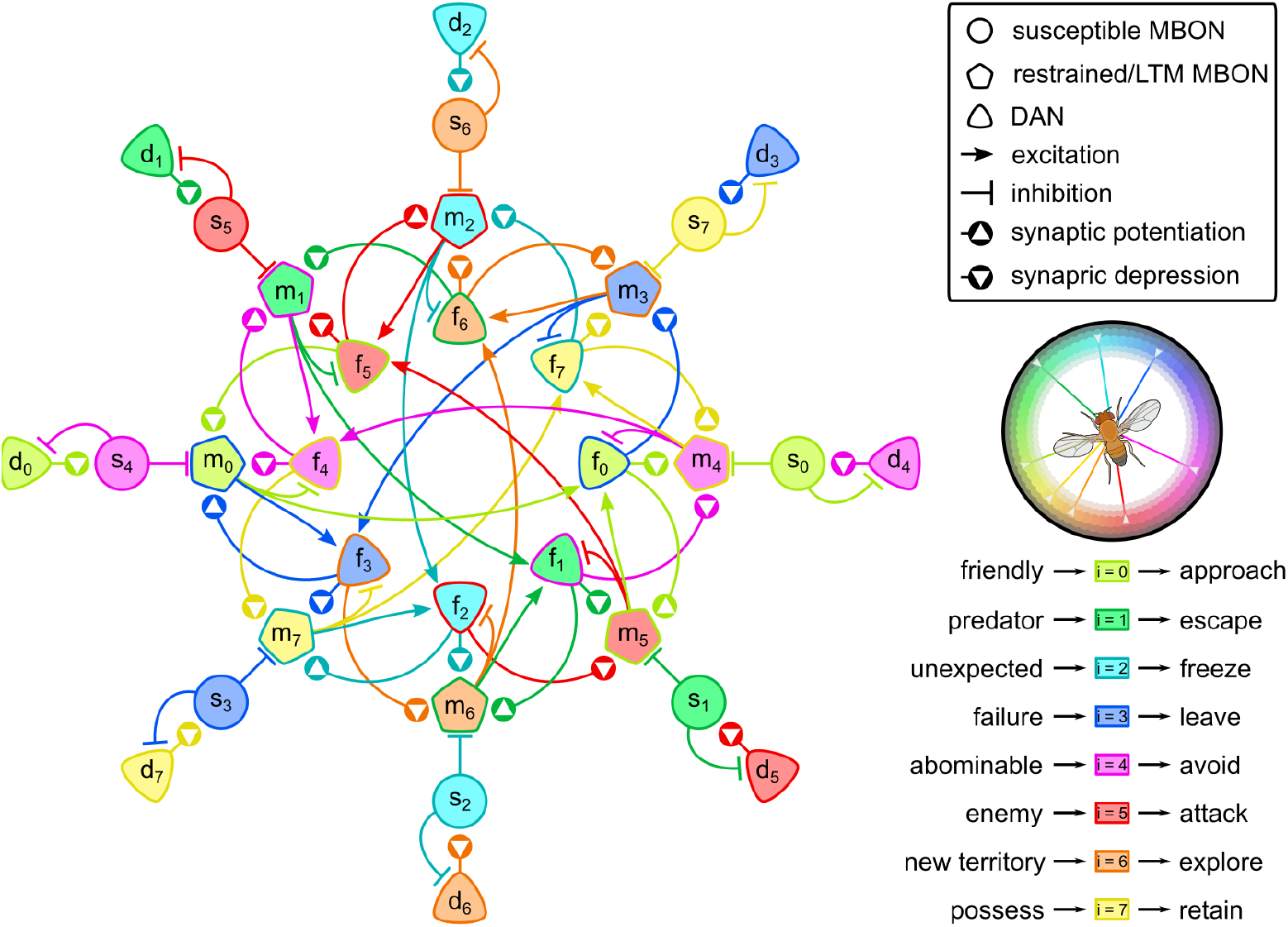
The “incentives wheel” model. This model supports that reinforcement is not binary but it draws its values from a spectrum. Different types of reinforcements trigger different dopaminergic neurons (DANs) that enable learning in different parts of the mushroom body. Colours show the variety of motivations that the model can encode associated with humans’‘wheel of emotions’ (***Plutchik, 2001***); e.g., <u>light green</u>: trust, <u>green</u>: fear, <u>light blue</u>: surprise, <u>blue</u>: sadness, <u>pink</u>: disgust, <u>red</u>: anger, <u>orange</u>: anticipation, <u>yellow</u>: joy. Neurons of more than one colours are part of multiple circuits that contribute in different motivations.

Real-life experiences are complicated and rich in information. This could produce a whole spectrum of reinforcements and motivations that guide the behaviour of animals. Data show that animals respond differently in different reinforcements, which cannot be represented just by the magnitude of a single variable, e.g., more/less rewarding/punishing. For example, different concentration of salt (***Zhang et al., 2013***), or sugar (***Colomb et al., 2009***) might be combined with the sated state of the animal, activate different subsets of DANs and trigger different behaviours, such as feeding or escaping. When the male fruit fly is exposed to female pheromones, courtship behaviour is triggered through P1 neurons (***Kallman et al., 2015***; ***Sten et al., 2020***), which can be translated to attraction, but has nothing to do with the appetite of the animal. On the other hand, other male pheromones trigger avoidance which suggests that a similar to the incentive circuit could explain this behaviour. The mushroom body has been proved to contribute in many behaviours other than olfactory classical conditioning, including visual navigation, and its output neurons encode richer information that is very close to humans’ decision making (***Heisenberg, 2003***).

It is reasonable to think that the ~ 34 output and ~ 130 dopaminergic neurons interacting with the mushroom bodies in the brain of fruit flies are not all used in order to discriminate odours and assign a positive or negative reinforcement to them driving attraction and avoidance. For this reason, we believe that the different MBONs do not represent different odours like it has been proposed before (***Huerta et al., 2004***) neither are split into two groups [e.g., attraction or avoidance, (***Schwaerzel et al., 2003***; ***Schroll et al., 2006***; ***Waddell, 2010***)], but rather represent different motivations of the animal that all together guide its overall behaviour (***Heisenberg, 2003***; ***Krashes et al., 2009***). These motivations are associated to different contexts, which are represented by the responses of the KCs as it has been proposed by ***Cohn et al.*** (***2015***), who showed that the same output neurons in the γ compartment respond differently when a different context is given. This context enables or disables different microcircuits of the mushroom body — similar to the ones described in Microcircuits of the mushroom body —which result in the activation of a subset of MBONs that represent different motivations, while the overlapping microcircuits result in what we sometimes call “noisy” or “insignificant” changes in the behaviour.

Appendix 1 ***Figure 1*** illustrates such a model, which we call the ‘incentive wheel’ (IW). In this model, we use 4 identical incentive memories (C0/4, C1/5, C2/6 and C3/7), where the Reciprocal short-term memories microcircuit of the one is the Reciprocal long-term memories microcircuit of another. As the structure of the RLM microcircuit is identical to the RSM one, we assume that the RLM of circuit C0/4 is the RSM of circuit C1/5, and the RLM of circuit C1/5 is the RSM of circuit C2/6 and so on. This way, we weave the different circuits into an incentive wheel with opposing motivations. The reinforcements that trigger the DANs in this model are drawn from a spectrum and the output of the MBONs of the model triggers different motivations. The LTMs and restrained memories can both exist in the same neurons of the core of the model, representing different motivations in different contexts. This might cause changes in the behaviour of the circuits that is irrelevant to the associated reinforcement, but relevant to a neighbouring reinforcement of the spectrum.

The ‘incentive wheel’ is an example of how the incentive circuit can be part of a bigger circuit that can provide a variety of motivations to the animal. An extension of it could have susceptible MBONs, *s_i_*, connecting to other susceptible MBONs from a parallel IW model with higher-order motivations. In Appendix 1 ***Figure 1***, we have associated the different motivations with the primary human emotions from the ‘wheel of emotions’ (***Plutchik, 2001***). Higher-order motivations could exist by combining primary motivations as if they were emotions and result in more complicated behaviours for the animal.

## Appendix 2

**Appendix 2 Table 1.** Additional information on the mushroom body neurons used for the incentive circuit, sorted by their short name. Information about the neurons has been collected from ***McCurdy et al.*** (***2021***) and ***Aso et al***. (***2014a***).

| Driver | Cluster | Neuron name | #cells | Short name | Alternative names |
| --- | --- | --- | --- | --- | --- |
| MB011B | M4/M6 | MBON-$\gamma 5\beta'2a$ | 1 | MBON-01 | MB-M6 |
| MB399B | M4/M6 | MBON-$\beta 2\beta'2a$ | 1 | MBON-02 | |
| MB298B | MV2 | MBON-$\gamma 4 > \gamma 1\gamma 2$ | 1 | MBON-05 | |
| MB112C | | MBON-$\gamma 1pedc > \alpha/\beta$ | 1 | MBON-11 | MB-MVP2 |
| MB077B | V3/V4 | MBON-$\gamma 2\alpha'1$ | 2 | MBON-12 | |
| MB050B | V2 | MBON-$\alpha'1$ | 3 | MBON-15 | |
| MB109B | PAM | PAM-$\beta'2a$ | 6-9 | PAM-02 | |
| MB301B | PAM | PAM-$\beta 2\beta'2a$ | > 3 | PAM-04 | subset of MB-M8 |
| MB312B | PAM | PAM-$\gamma 4 < \gamma 1\gamma 2$ | 13-17 | PAM-07 | subset of MB-AIM? |
| MB320C | PPL1 | PPL1-$\gamma 1ped$ | 1 | PPL1-01 | MB-MP1, MP |
| MB296B<sub>1</sub> | PPL1 | PPL1-$\gamma 2\alpha'1_1$ | 1 | PPL1-03 | MB-MV1 |
| MB296B<sub>2</sub> | PPL1 | PPL1-$\gamma 2\alpha'1_2$ | 1 | PPL1-03 | MB-MV1 |

---

**Figure 3–Figure supplement 1.**
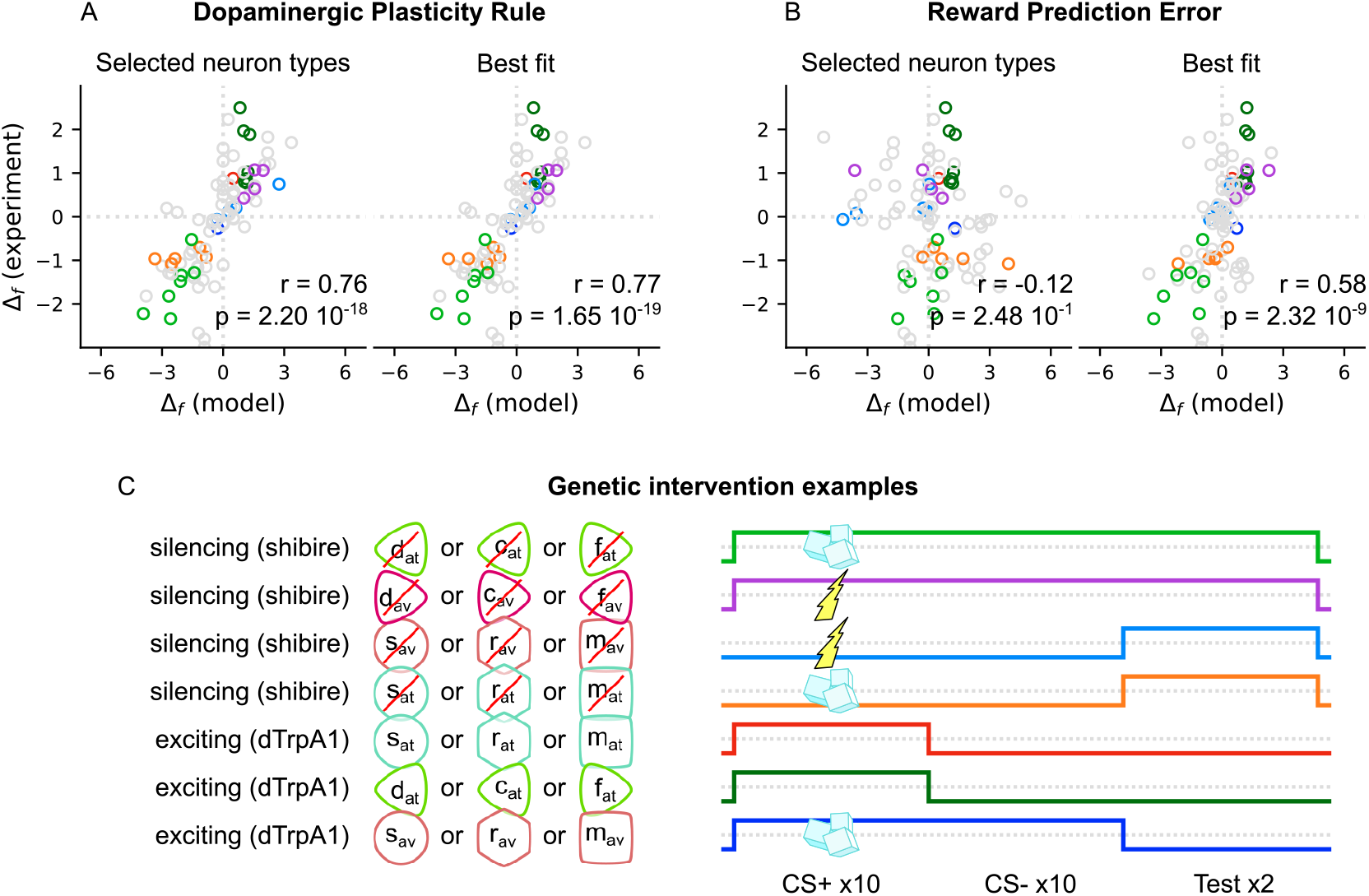
Testing the performance of the incentive circuit in the experiments of ***Bennett et al.*** (***2021***) — **Figure 5**. ***Bennett et al.*** (***2021***) collected behavioural data (Δ*f* measure) from 92 experiments, summarised in ***Figure 3***–***source data 1***, and calculate their correlation coefficient to the behaviour produced by their model (VSλmodel: *r* = 0.68, *p* < 10^−4^; MV model: *r* = 0.65, *p* < 10^−4^). The behavioural data involve intervention (e.g., activation or silencing) in different MBONs or DANs, which are grouped by colour codes. Here we use the same colour codes as in the original paper for convenience. For the details of this analysis please refer to ***Bennett et al.*** (***2021***). We test the behaviour of the incentive circuit when using two different learning rules: (A) the DPR and (B) the RPE. As we do not know what type the intervened MBON or DAN is, we test for all the types (and groups of them) and we report the ones with the highest correlation under the “Best fit” plot. We also try to guess the type by the identity of the neuron (or group of neurons) intervened and we report the correlation coefficient under the “Selected neuron types” plot. The types selected for each experiment can also be found in ***Figure 3***–***source data 1***. (C) Examples of the genetic intervention as coloured in A and B. We run 10 trials of odour A + shock or sugar (indicated with a thunder or cubes respectively) or without reinforcement (absence of thunder and sugar cubes), followed by 10 trials of odour B without reinforcement (acquisition phase). Then proceed with two trials of testing odour A vs odour B (extinction phase). In addition, we model genetic intervention by targeting selected neurons of our model and silence them via the shibire blockage or excite them through the dTrpA1 channel (timing of the intervention is shown by the coloured lines). The colours of the lines correspond to the different examples to the coloured samples of A and B.

**Figure 4–Figure supplement 1.**
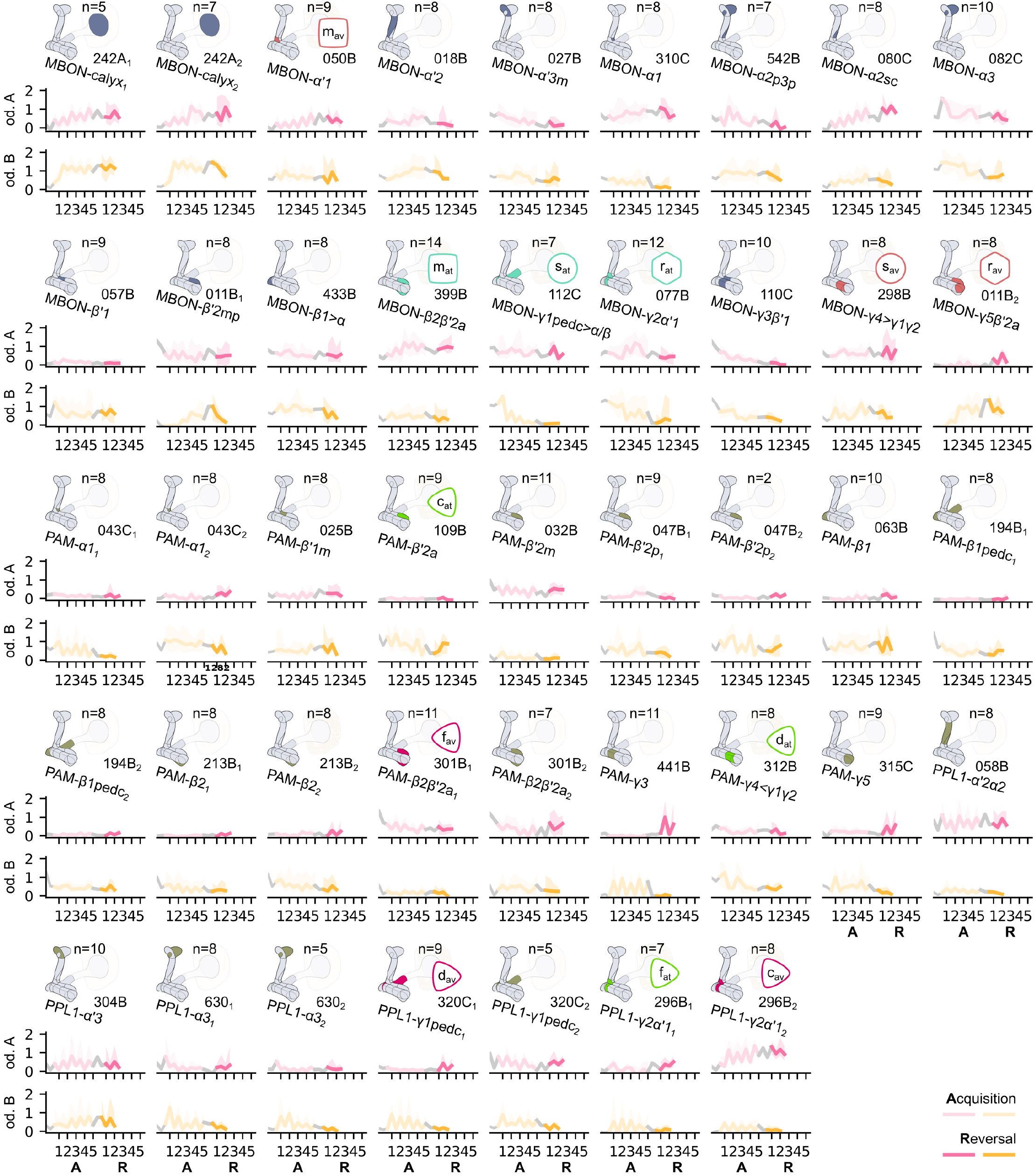
The responses from all the recorded neurons in the *Drosophila melanogaster* mushroom body during the experiment described in **Figure 4**B. In each row we present the driver of the recorded neurons and a schematic representation of where they innervate the mushroom body;the median responses of the neuron for each odour (coloured as pink for odour A or yellow for odour B) over a number of flies, denoted as *n*, and the 25% and 75% quantiles marked by the coloured region.

**Figure 5–Figure supplement 1.**
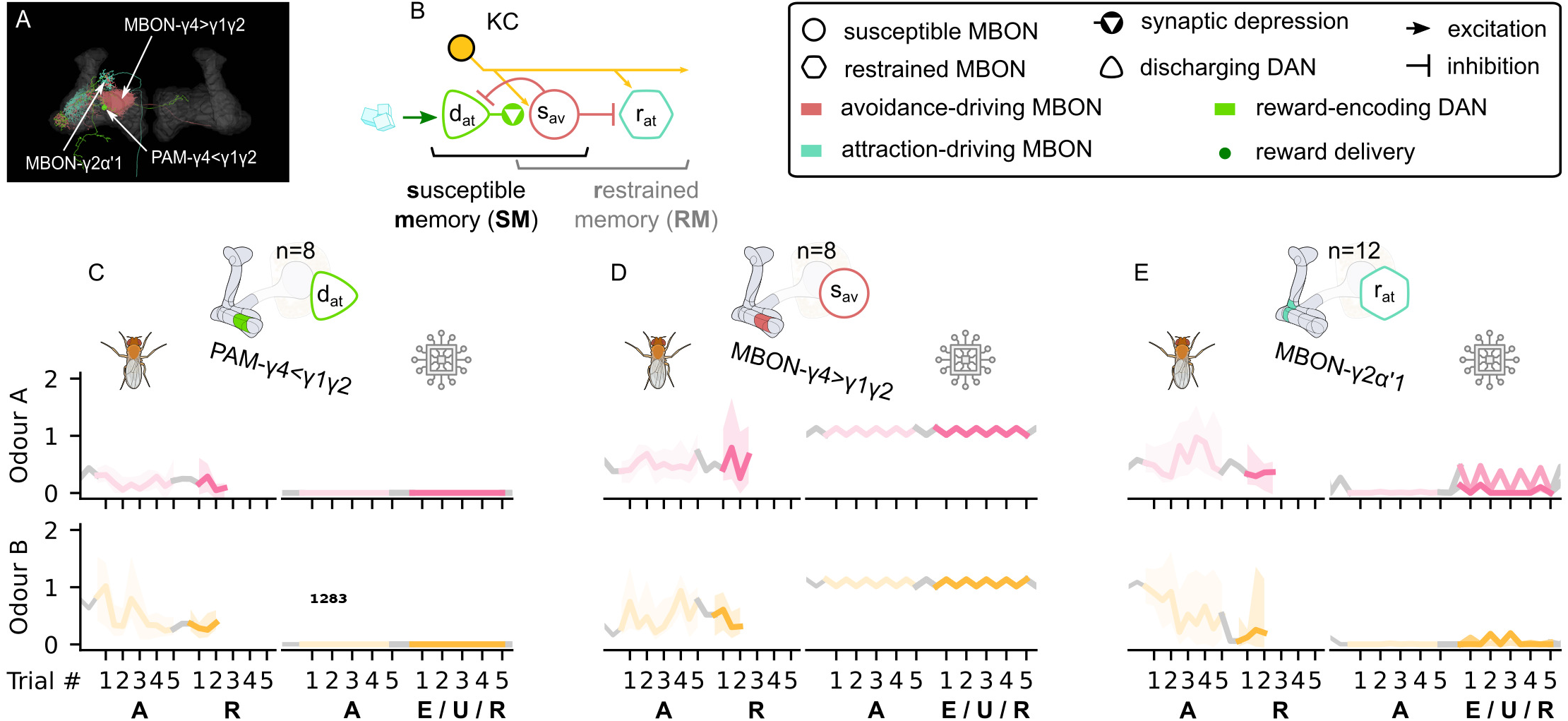
The susceptible and restrained microcircuits of the mushroom body. (A) Image of the avoidance-driving susceptible and attraction-driving restrained memory microcircuits made of the PAM-γ4<γ1γ2, MBON-γ4>γ1γ2 and MBON-–2α’ 1 neurons - created using the Virtual Fly Brain software (***Milyaev et al., 2012***). (B) Schematic representation of the Susceptible & restrained memories microcircuits connected via the susceptible MBON. The responses of (C) the reward-encoding discharging DAN, *d*_at_, (D) the avoidance-driving susceptible MBON, *s*_av_, and (E) the attraction-driving restrained MBON, *r*_at_, generated by experimental data (left) and the model (right) during the olfactory conditioning paradigms of **Figure 4**D. Lightest shades denote the extinction, mid shades the unpaired and dark shades the reversal phase. For each trial we report two consecutive time-steps: the off-shock (i.e., odour only) followed by the on-shock (i.e., paired odour and shock) when available (i.e., odour B in acquisition and odour A in reversal phase) otherwise a second off-shock time-step (i.e., all the other phases).

**Figure 5–Figure supplement 2.**
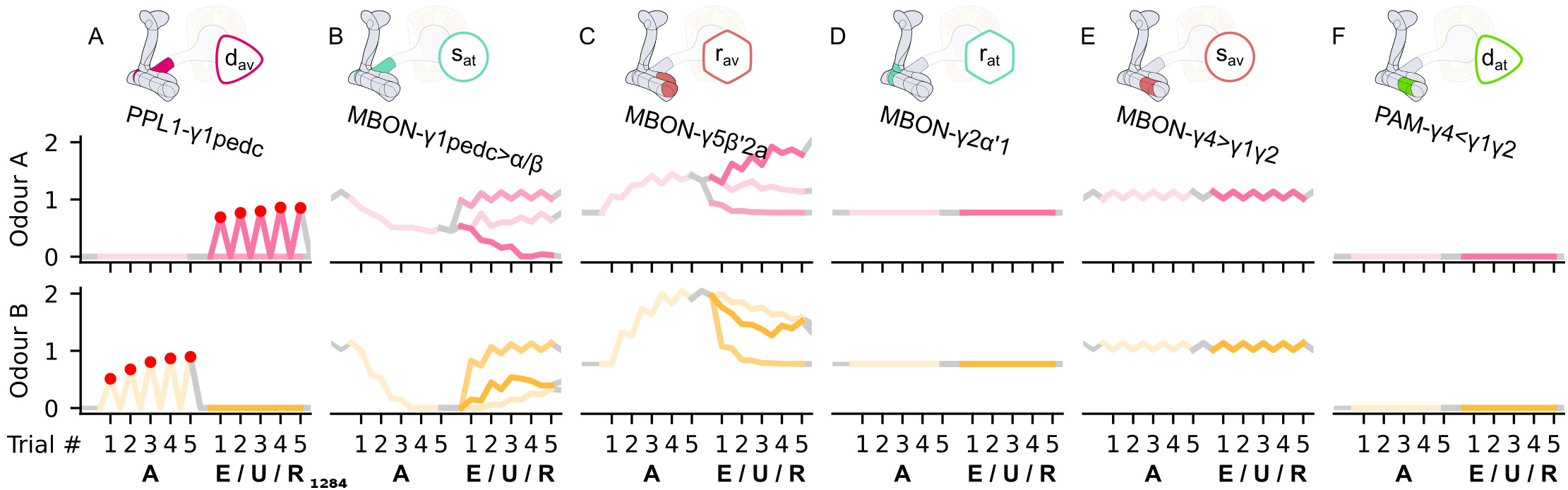
The responses of the neurons of the incentive circuit using only the connections of the SM and RM microcircuits. The responses of (A) the punishment-encoding discharging DAN, *d*_av_, (B) the attraction-driving susceptible MBON, *s*_at_, (C) the avoidance-driving restrained MBON, *r*_av_, (D) the attraction-driving restrained MBON, *r*_at_, (E) the avoidance-driving susceptible MBON, *s*_av_, and (F) the reward-encoding discharging DAN, *d*_at_, generated by experimental data (left) and the model (right) during the olfactory conditioning paradigms of **Figure 4**D. Lightest shades denote the extinction, mid shades the unpaired and dark shades the reversal phase. The first row of responses (coloured pink) corresponds to responses associated with odour A, and the second (coloured yellow) to those associated with odour B. For each trial we report two consecutive time-steps: the off-shock (i.e., odour only) followed by the on-shock (i.e., paired odour and shock) when available (i.e., odour B in acquisition and odour A in reversal phase) otherwise a second off-shock time-step (i.e., all the other phases).

**Figure 5–Figure supplement 3.**
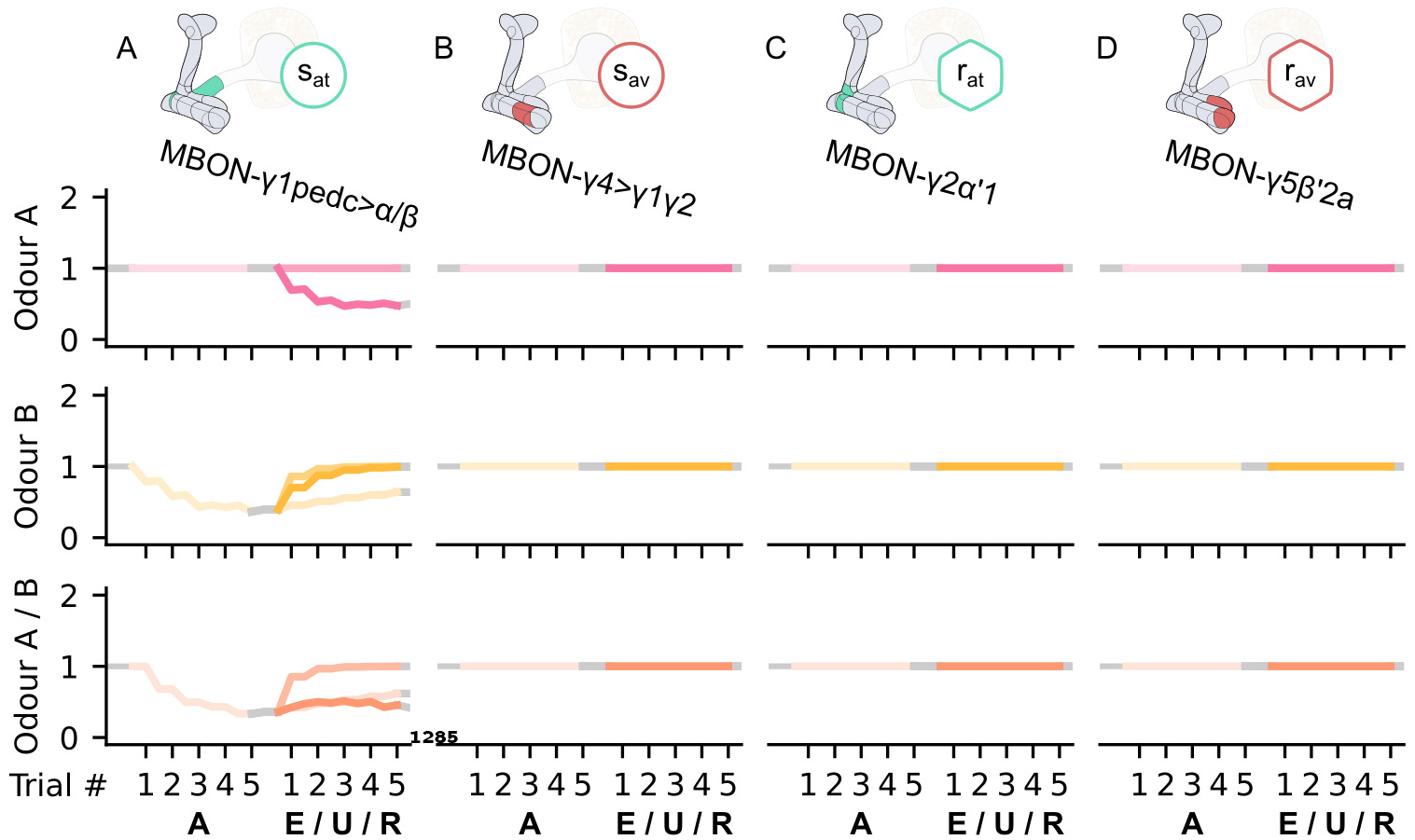
The KC→MBON synaptic weights of the neurons of the incentive circuit using only the connections of the SM and RM microcircuits. The synaptic weights of (A) the attraction-driving susceptible MBON, *s*_at_, (B) the avoidance-driving susceptible MBON, *s*_av_, (C) the attraction-driving restrained MBON, *r*_at_, and (D) the avoidance-driving restrained MBON, *r*_av_, generated by experimental data (left) and the model (right) during the olfactory conditioning paradigms of **Figure 4**D. Lightest shades denote the extinction, mid shades the unpaired and dark shades the reversal phase. The first row of weights (coloured pink) corresponds to KCs associated with odour A, the second (coloured yellow) to KCs associated with odour B and the third (coloured orange) to KCs associated with both odours. For each trial we report two consecutive time-steps: the off-shock (i.e., odour only) followed by the on-shock (i.e., paired odour and shock) when available (i.e., odour B in acquisition and odour A in reversal phase) otherwise a second off-shock time-step (i.e., all the other phases).

**Figure 6–Figure supplement 1.**
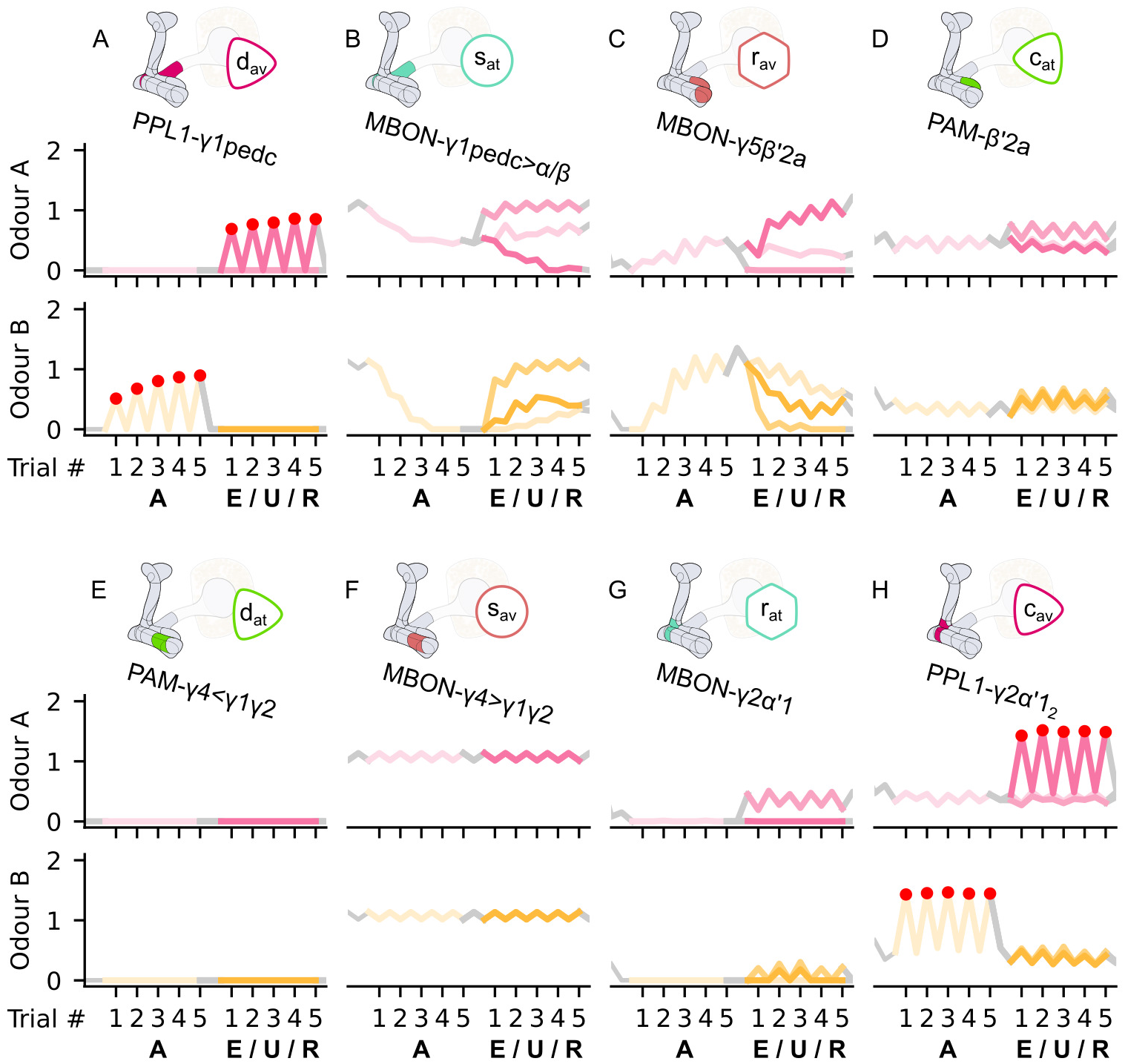
The responses of the neurons of the incentive circuit using only the connections of the SM, RM and RSM microcircuits. The responses of (A) the punishment-encoding discharging DAN, *d*_av_, (B) the attraction-driving susceptible MBON, *s*_at_, (C) the avoidance-driving restrained MBON, *r*_av_, (D) the reward-encoding charging DAN, *c*_at_, (E) the reward-encoding discharging DAN, *d*_at_, (F) the avoidance-driving susceptible MBON, *s*_av_, (G) the attraction-driving restrained MBON, *r*_at_, and (H) the punishment-encoding charging DAN, *c*_av_, generated by experimental data (left) and the model (right) during the olfactory conditioning paradigms of **Figure 4**D. Lightest shades denote the extinction, mid shades the unpaired and dark shades the reversal phase. The first row of responses (coloured pink) corresponds to responses associated with odour A, and the second (coloured yellow) to those associated with odour B. For each trial we report two consecutive time-steps: the off-shock (i.e., odour only) followed by the on-shock (i.e., paired odour and shock) when available (i.e., odour B in acquisition and odour A in reversal phase) otherwise a second off-shock time-step (i.e., all the other phases).

**Figure 6–Figure supplement 2.**
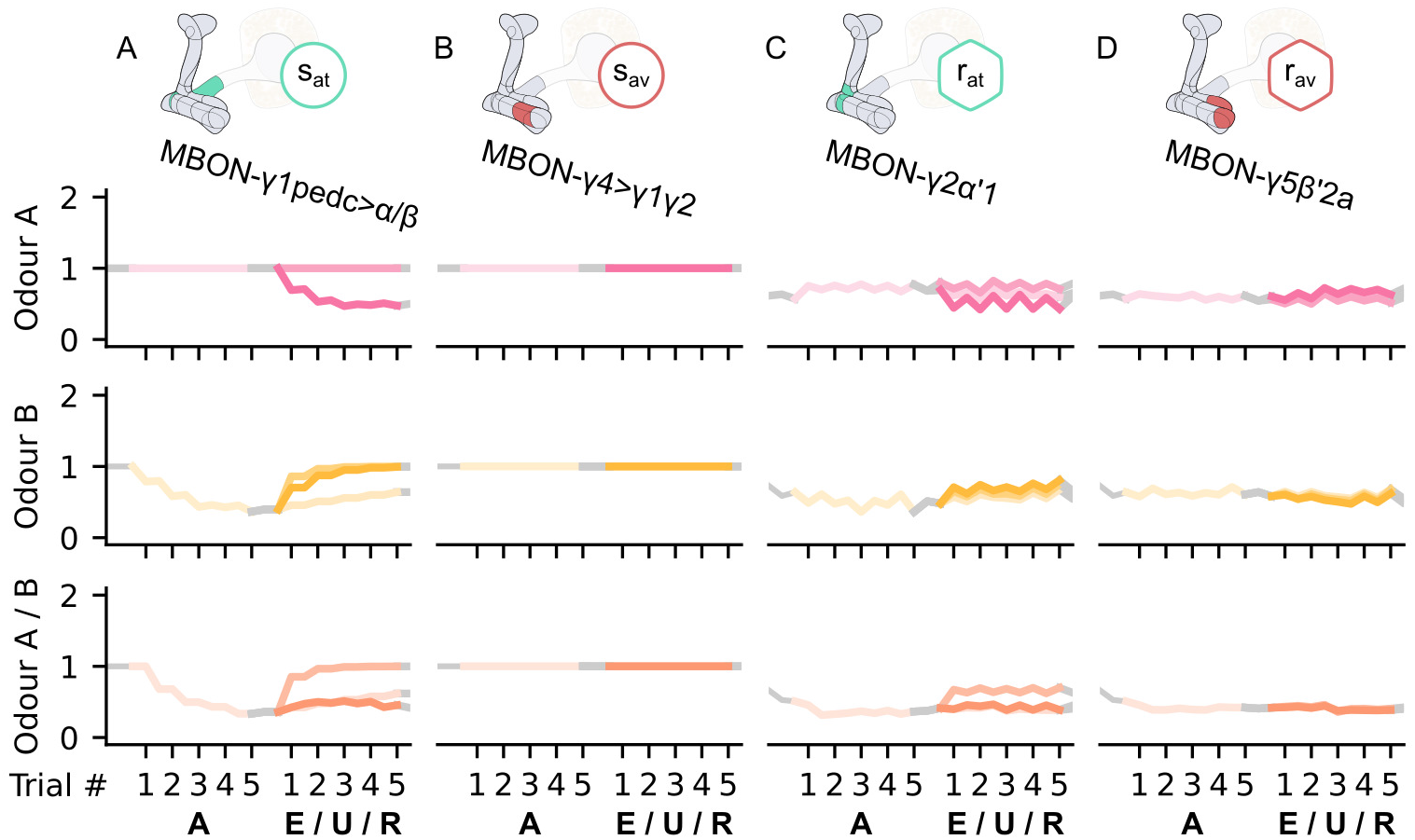
The KC→MBON synaptic weights of the neurons of the incentive circuit using only the connections of the SM, RM and RSM microcircuits. The synaptic weights of (A) the attraction-driving susceptible MBON, *s*_at_, (B) the avoidance-driving susceptible MBON, *s*_av_, (C) the attraction-driving restrained MBON, *r*_at_, and (D) the avoidance-driving restrained MBON, *r*_av_, generated by experimental data (left) and the model (right) during the olfactory conditioning paradigms of **Figure 4**D. Lightest shades denote the extinction, mid shades the unpaired and dark shades the reversal phase. The first row of weights (coloured pink) corresponds to KCs associated with odour A, the second (coloured yellow) to KCs associated with odour B and the third (coloured orange) to KCs associated with both odours. For each trial we report two consecutive time-steps: the off-shock (i.e., odour only) followed by the on-shock (i.e., paired odour and shock) when available (i.e., odour B in acquisition and odour A in reversal phase) otherwise a second off-shock time-step (i.e., all the other phases).

**Figure 7–Figure supplement 1.**
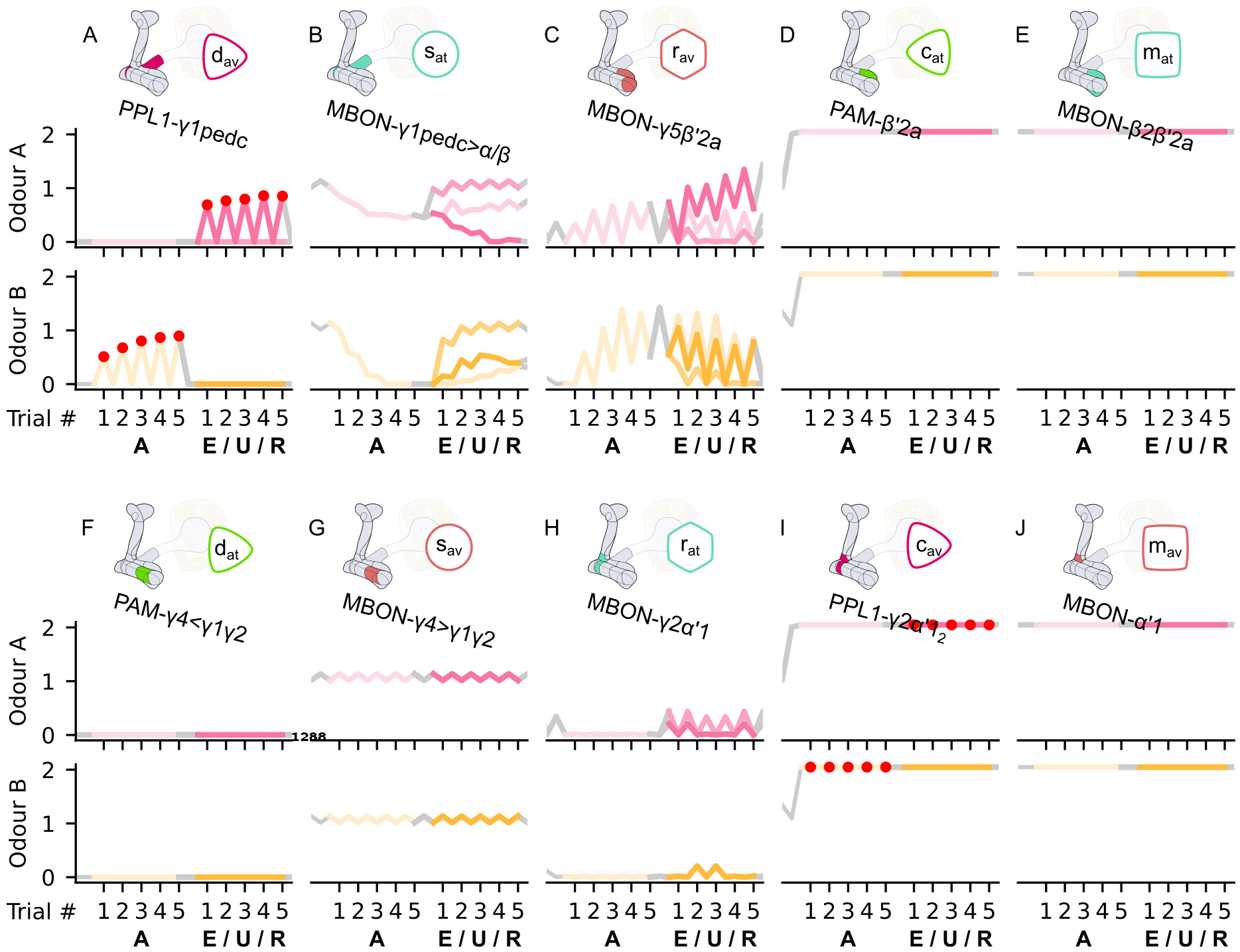
The responses of the neurons of the incentive circuit using only the connections of the SM, RM, RSM and LTM microcircuits. The responses of (A) the punishment-encoding discharging DAN, *d*_av_, (B) the attraction-driving susceptible MBON, *s*_at_, (C) the avoidance-driving restrained MBON, *r*_av_, (D) the reward-encoding charging DAN, *c*_at_, (E) the attraction-driving long-term memory MBON, *m*_at_, (F) the reward-encoding discharging DAN, *d*_at_, (G) the avoidance-driving susceptible MBON, *s*_av_, (H) the attraction-driving restrained MBON, *r*_at_, (I) the punishment-encoding charging DAN, *c*_av_, and (J) the avoidance-driving long-term memory MBON, *m_av_*, generated by experimental data (left) and the model (right) during the olfactory conditioning paradigms of **Figure 4**D. Lightest shades denote the extinction, mid shades the unpaired and dark shades the reversal phase. The first row of responses (coloured pink) corresponds to responses associated with odour A, and the second (coloured yellow) to those associated with odour B. For each trial we report two consecutive time-steps: the off-shock (i.e., odour only) followed by the on-shock (i.e., paired odour and shock) when available (i.e., odour B in acquisition and odour A in reversal phase) otherwise a second off-shock time-step (i.e., all the other phases).

**Figure 7–Figure supplement 2.**
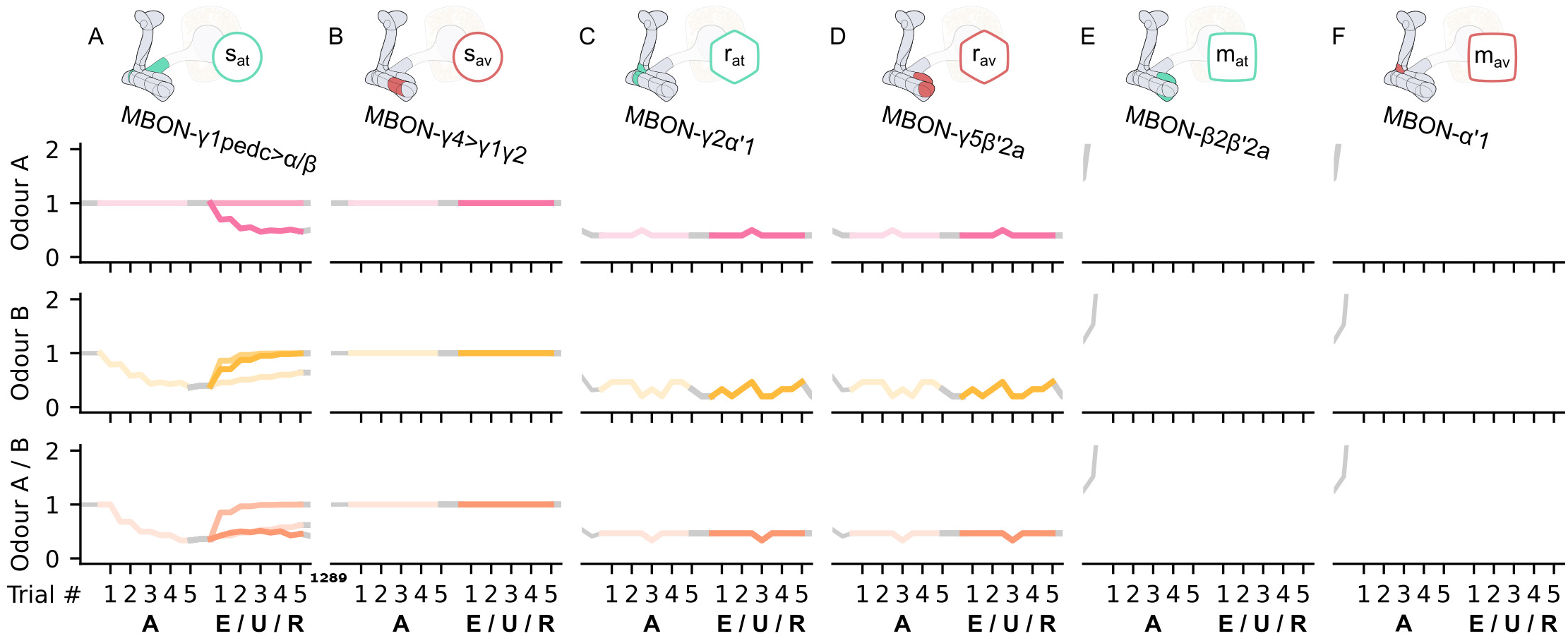
The KC→MBON synaptic weights of the neurons of the incentive circuit using only the connections of the SM, RM, RSM and LTM microcircuits. The synaptic weights of (A) the attraction-driving susceptible MBON, *s*_at_, (B) the avoidance-driving susceptible MBON, *s*_av_, (C) the attraction-driving restrained MBON, *r*_at_, (D) the avoidance-driving restrained MBON, *r*_av_, (E) the attraction-driving long-term memory MBON, *m*_at_, and (F) the avoidance-driving long-term memory MBON, *m*_av_, generated by experimental data (left) and the model (right) during the olfactory conditioning paradigms of **Figure 4**D. Lightest shades denote the extinction, mid shades the unpaired and dark shades the reversal phase. The first row of weights (coloured pink) corresponds to KCs associated with odour A, the second (coloured yellow) to KCs associated with odour B and the third (coloured orange) to KCs associated with both odours. For each trial we report two consecutive time-steps: the off-shock (i.e., odour only) followed by the on-shock (i.e., paired odour and shock) when available (i.e., odour B in acquisition and odour A in reversal phase) otherwise a second off-shock time-step (i.e., all the other phases).

**Figure 8–Figure supplement 1.**
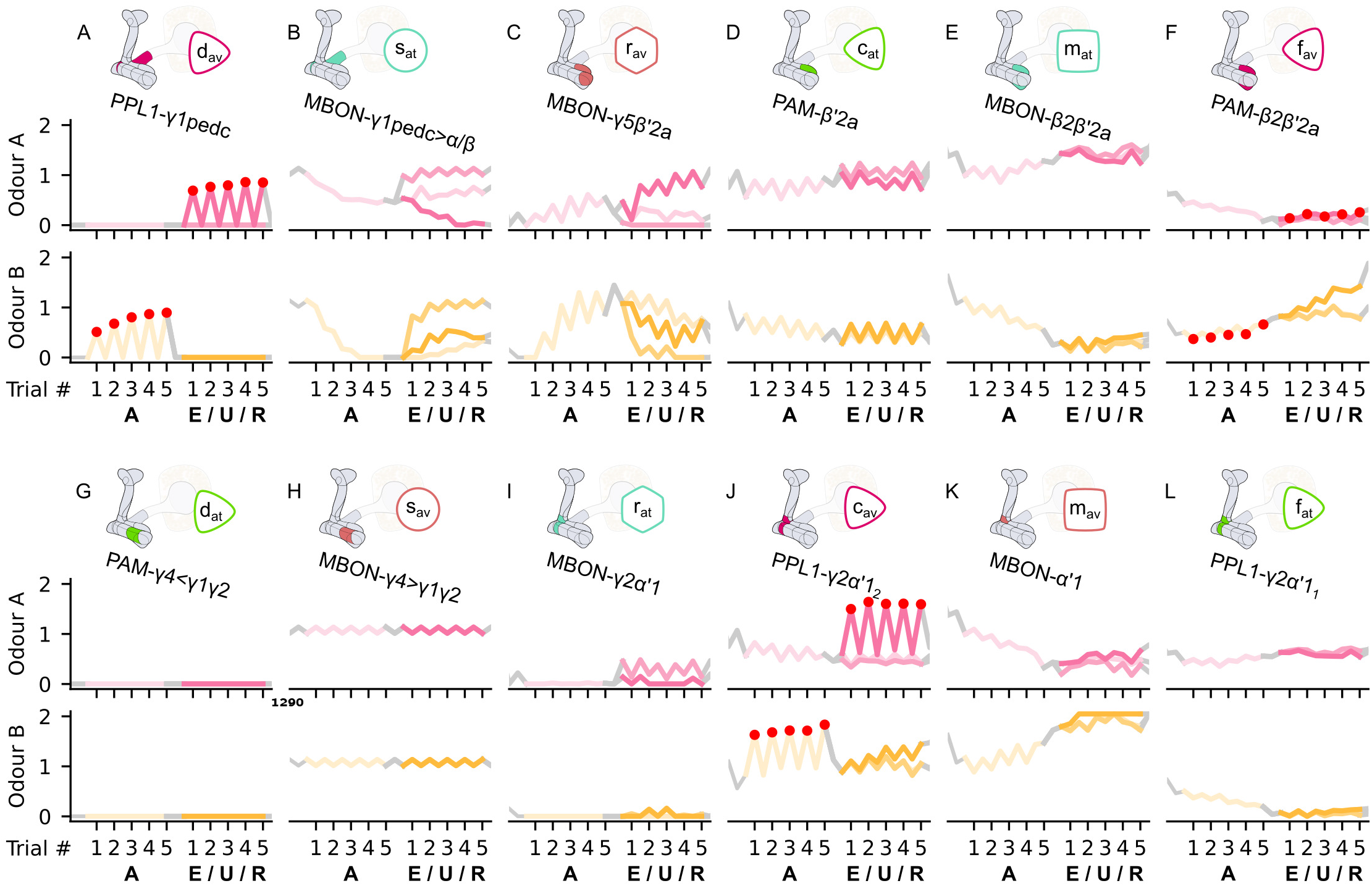
The responses of the neurons of the incentive circuit using only the connections of the SM, RM, RSM, LTM and RLM microcircuits. The responses of (A) the punishment-encoding discharging DAN, *d*_av_, (B) the attraction-driving susceptible MBON, *s*_at_, (C) the avoidance-driving restrained MBON, *r*_av_, (D) the reward-encoding charging DAN, *c*_at_, (E) the attraction-driving long-term memory MBON, *m*_at_, (F) the avoidance-driving forgetting DAN, *f*_av_, (G) the reward-encoding discharging DAN, *d*_at_, (H) the avoidance-driving susceptible MBON, *s*_av_, (I) the attraction-driving restrained MBON, *r*_at_, (J) the punishment-encoding charging DAN, *c*_av_, (K) the avoidance-driving long-term memory MBON, *m*_av_, and (L) the attraction-driving forgetting DAN, *f*_at_, generated by experimental data (left) and the model (right) during the olfactory conditioning paradigms of **Figure 4**D. Lightest shades denote the extinction, mid shades the unpaired and dark shades the reversal phase. The first row of responses (coloured pink) corresponds to responses associated with odour A, and the second (coloured yellow) to those associated with odour B. For each trial we report two consecutive time-steps: the off-shock (i.e., odour only) followed by the on-shock (i.e., paired odour and shock) when available (i.e., odour B in acquisition and odour A in reversal phase) otherwise a second off-shock time-step (i.e., all the other phases).

**Figure 8–Figure supplement 2.**
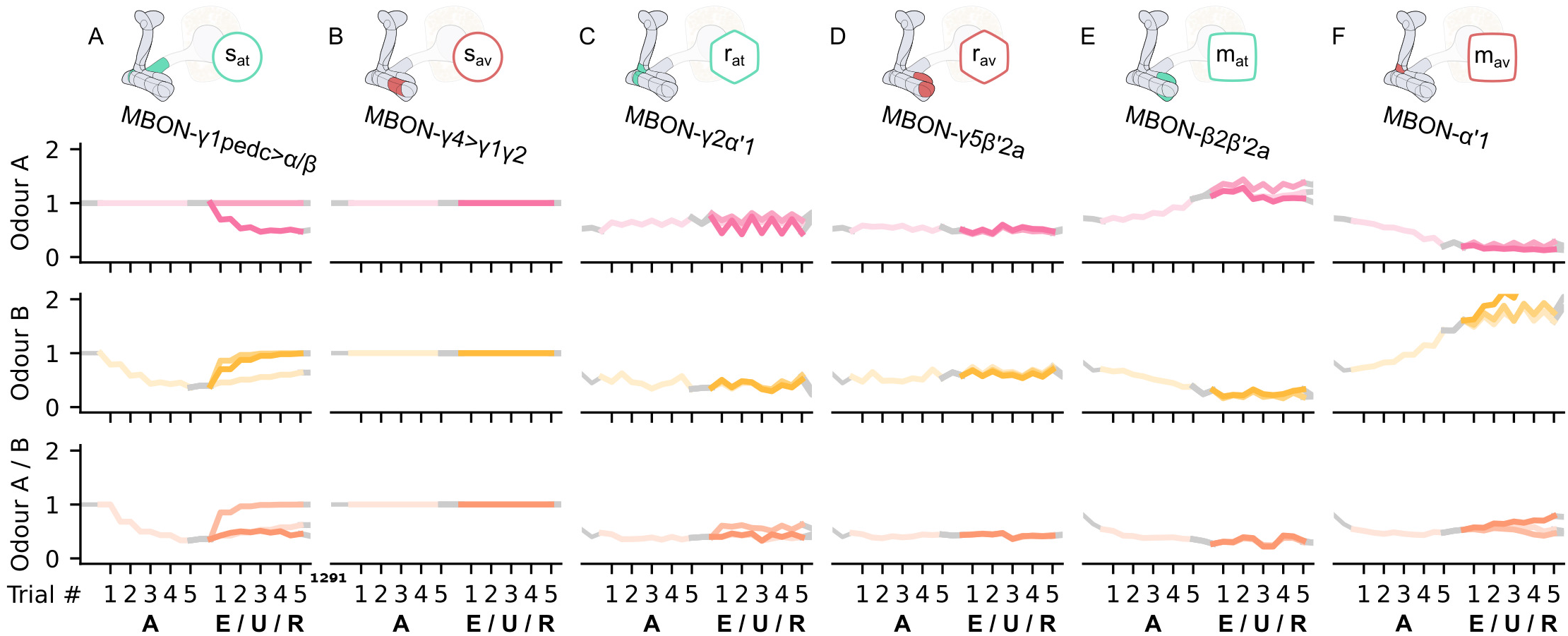
The KC→MBON synaptic weights of the neurons of the incentive circuit using only the connections of the SM, RM, RSM, LTM and RLM microcircuits. The synaptic weights of (A) the attraction-driving susceptible MBON, *s*_at_, (B) the avoidance-driving susceptible MBON, *s*_av_, (C) the attraction-driving restrained MBON, *r*_at_, (D) the avoidance-driving restrained MBON, *r*_av_, (E) the attraction-driving long-term memory MBON, *m*_at_, and (F) the avoidance-driving long-term memory MBON, *m*_av_, generated by experimental data (left) and the model (right) during the olfactory conditioning paradigms of **Figure 4**D. Lightest shades denote the extinction, mid shades the unpaired and dark shades the reversal phase. The first row of weights (coloured pink) corresponds to KCs associated with odour A, the second (coloured yellow) to KCs associated with odour B and the third (coloured orange) to KCs associated with both odours. For each trial we report two consecutive time-steps: the off-shock (i.e., odour only) followed by the on-shock (i.e., paired odour and shock) when available (i.e., odour B in acquisition and odour A in reversal phase) otherwise a second off-shock time-step (i.e., all the other phases).

**Figure 9–Figure supplement 1.**
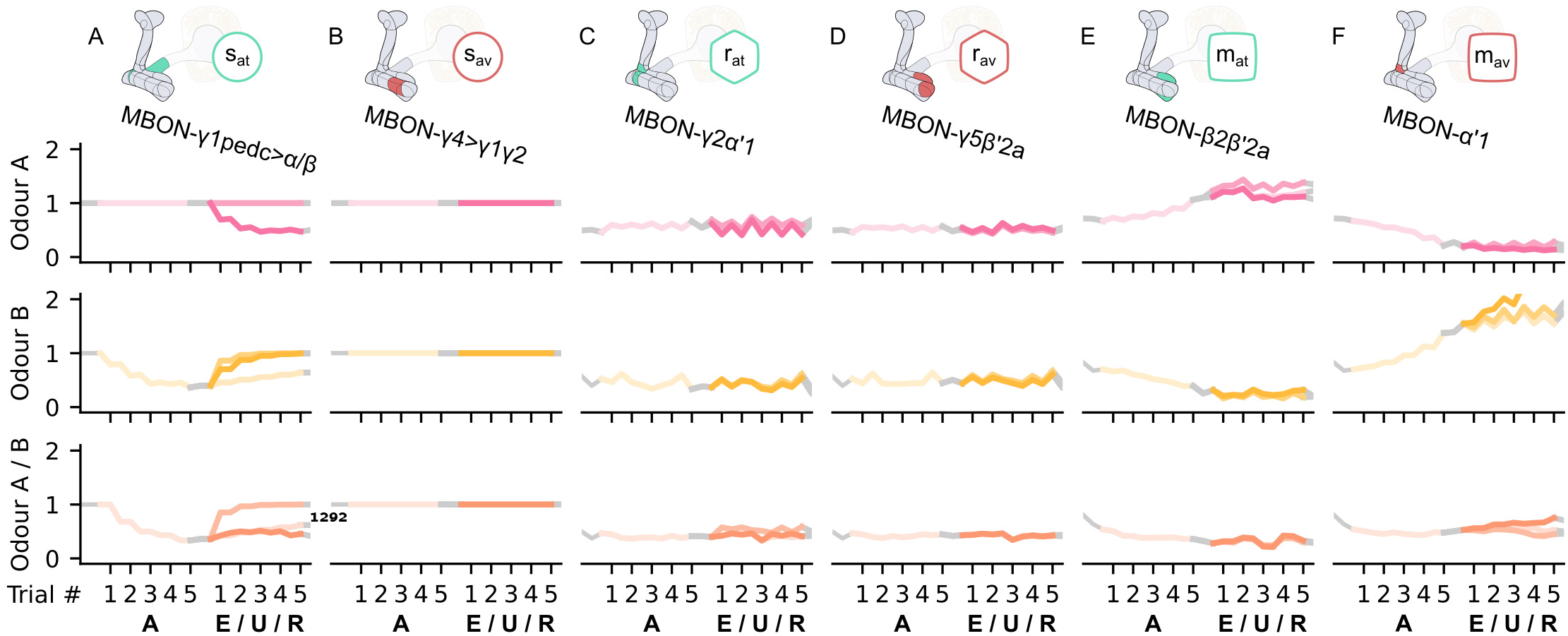
The KC→MBON synaptic weights of the neurons of the incentive circuit using all its connections. The synaptic weights of (A) the attraction-driving susceptible MBON, *s*_at_, (B) the avoidance-driving susceptible MBON, *s*_av_, (C) the attraction-driving restrained MBON, *r*_at_, (D) the avoidance-driving restrained MBON, *r*_av_, (E) the attraction-driving long-term memory MBON, *m*_at_, and (F) the avoidance-driving long-term memory MBON, *m*_av_, generated by experimental data (left) and the model (right) during the olfactory conditioning paradigms of **Figure 4**D. Lightest shades denote the extinction, mid shades the unpaired and dark shades the reversal phase. The first row of weights (coloured pink) corresponds to KCs associated with odour A, the second (coloured yellow) to KCs associated with odour B and the third (coloured orange) to KCs associated with both odours.

**Figure 9–Figure supplement 2.**
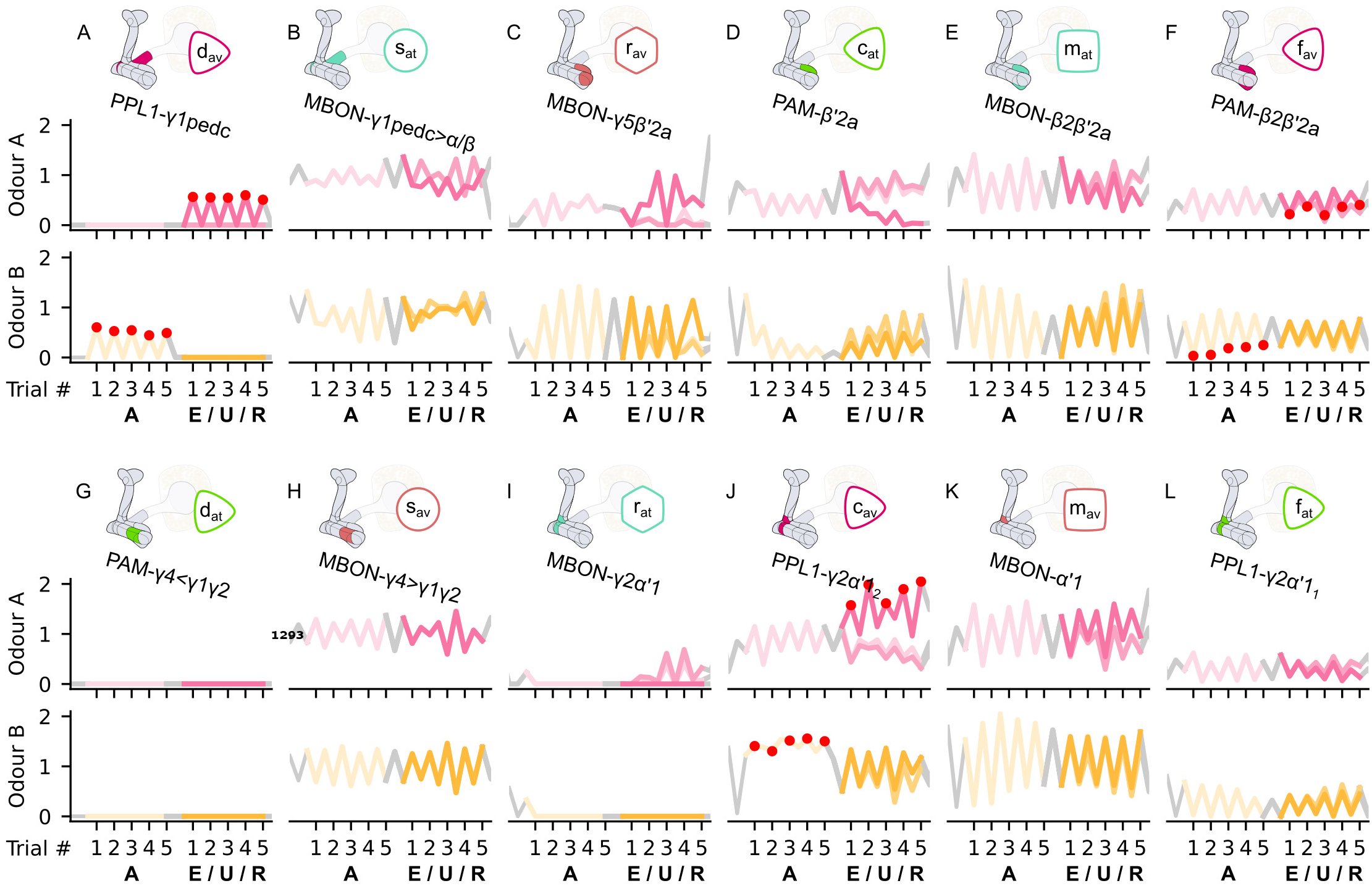
The reconstructed responses of the neurons of the incentive circuit using the reward prediction error learning rule. The reconstructed responses of (A) the punishment-encoding discharging DAN, *d*_av_, (B) the attraction-driving susceptible MBON, *s*_at_, (C) the avoidance-driving restrained MBON, *r*_av_, (D) the reward-encoding charging DAN, *c*_at_, (E) the attraction-driving long-term memory MBON, *m*_at_, (F) the punishment-encoding forgetting DAN, *f*_av_, (G) the reward-encoding discharging DAN, *d*_at_, (H) the avoidance-driving susceptible MBON, *s*_av_, (I) the attraction-driving restrained MBON, *r*_at_, (J) the punishment-encoding charging DAN, *c*_av_, (K) the avoidance-driving long-term memory MBON, *m*_av_, and (L) the reward-encoding forgetting DAN, *f*_at_, generated by experimental data (left) and the model (right) during the olfactory conditioning paradigms of **Figure 4**D. Lightest shades denote the extinction, mid shades the unpaired and dark shades the reversal phase.

**Figure 9–Figure supplement 3.**
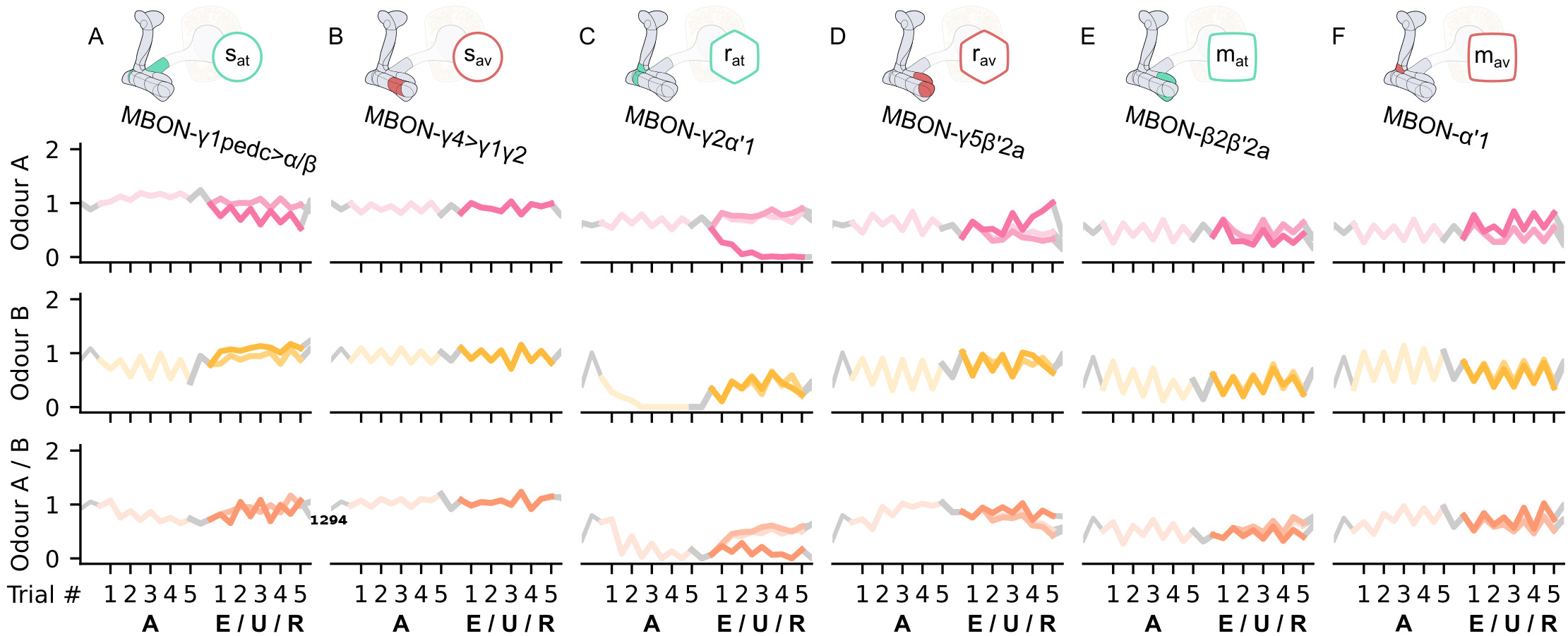
The KC→MBON synaptic weights of the neurons of the incentive circuit using the reward prediction error learning rule. The synaptic weights of (A) the attraction-driving susceptible MBON, *s*_at_, (B) the avoidance-driving susceptible MBON, *s*_av_, (C) the attraction-driving restrained MBON, *r*_at_, (D) the avoidance-driving restrained MBON, *r*_av_, (E) the attraction-driving long-term memory MBON, *m*_at_, and (F) the avoidance-driving long-term memory MBON, *m*_av_, generated by experimental data (left) and the model (right) during the olfactory conditioning paradigms of **Figure 4**D. Lightest shades denote the extinction, mid shades the unpaired and dark shades the reversal phase. The first row of weights (coloured pink) corresponds to KCs associated with odour A, the second (coloured yellow) to KCs associated with odour B and the third (coloured orange) to KCs associated with both odours.

**Figure 11–Figure supplement 1.**
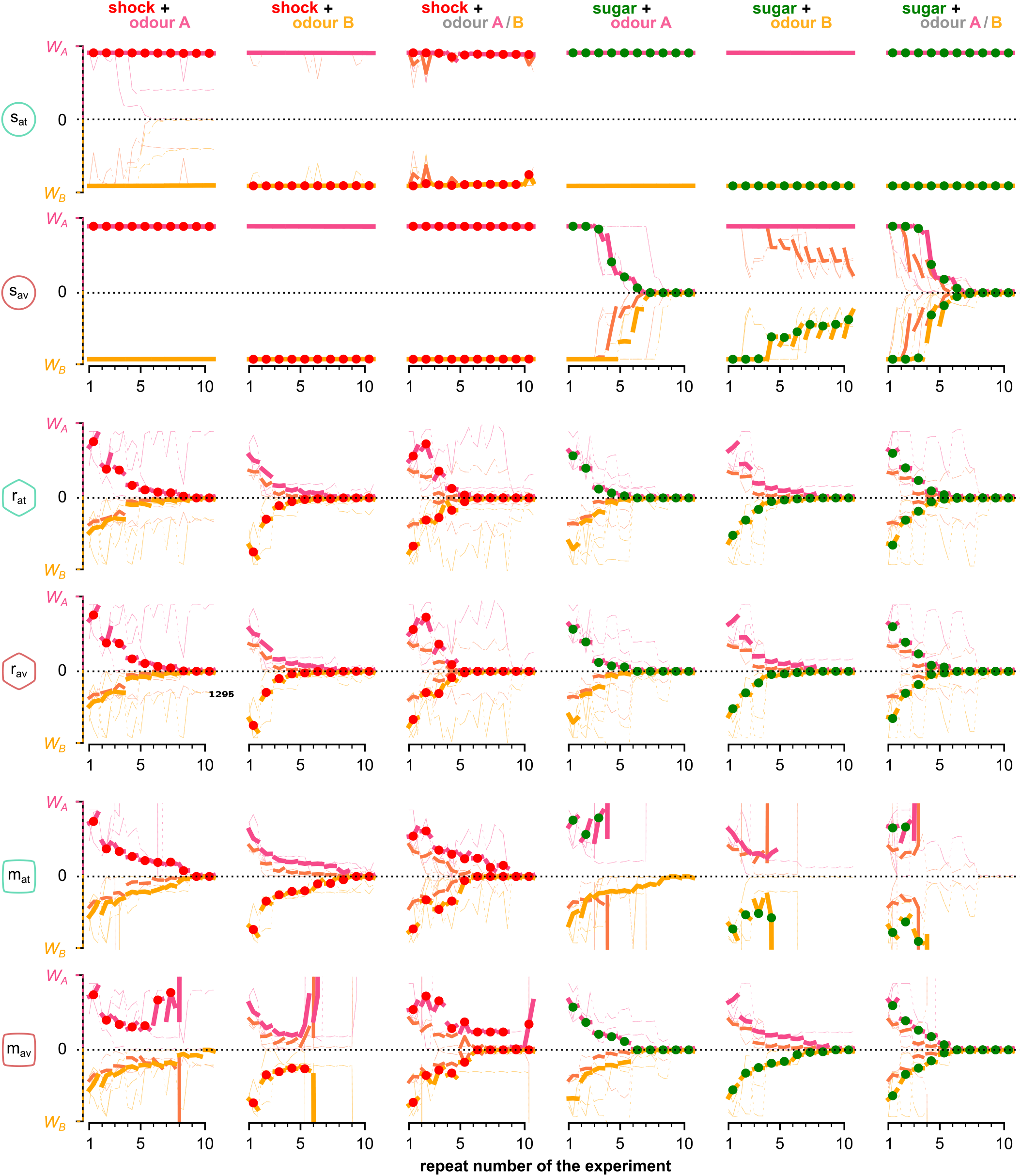
The mean KC→MBON synaptic weights over the simulated flies that visited both odours and for each neuron, phase and repeat of the experiment. Different rows correspond to KC→MBON weights associated with a different type of MBON, from top to bottom: attraction- and avoidance driving susceptible, restrained and long-term memory MBONs. Pink and yellow lines show synaptic weights associated with odour A and B respectively, dashed lines show weights associated with both odours. Thin lines show 3 representative examples of synaptic weights in single simulated flies. Thick lines show the median of the synaptic weight over all the simulated flies that visited both odours.

**Figure 11–Figure supplement 2.**
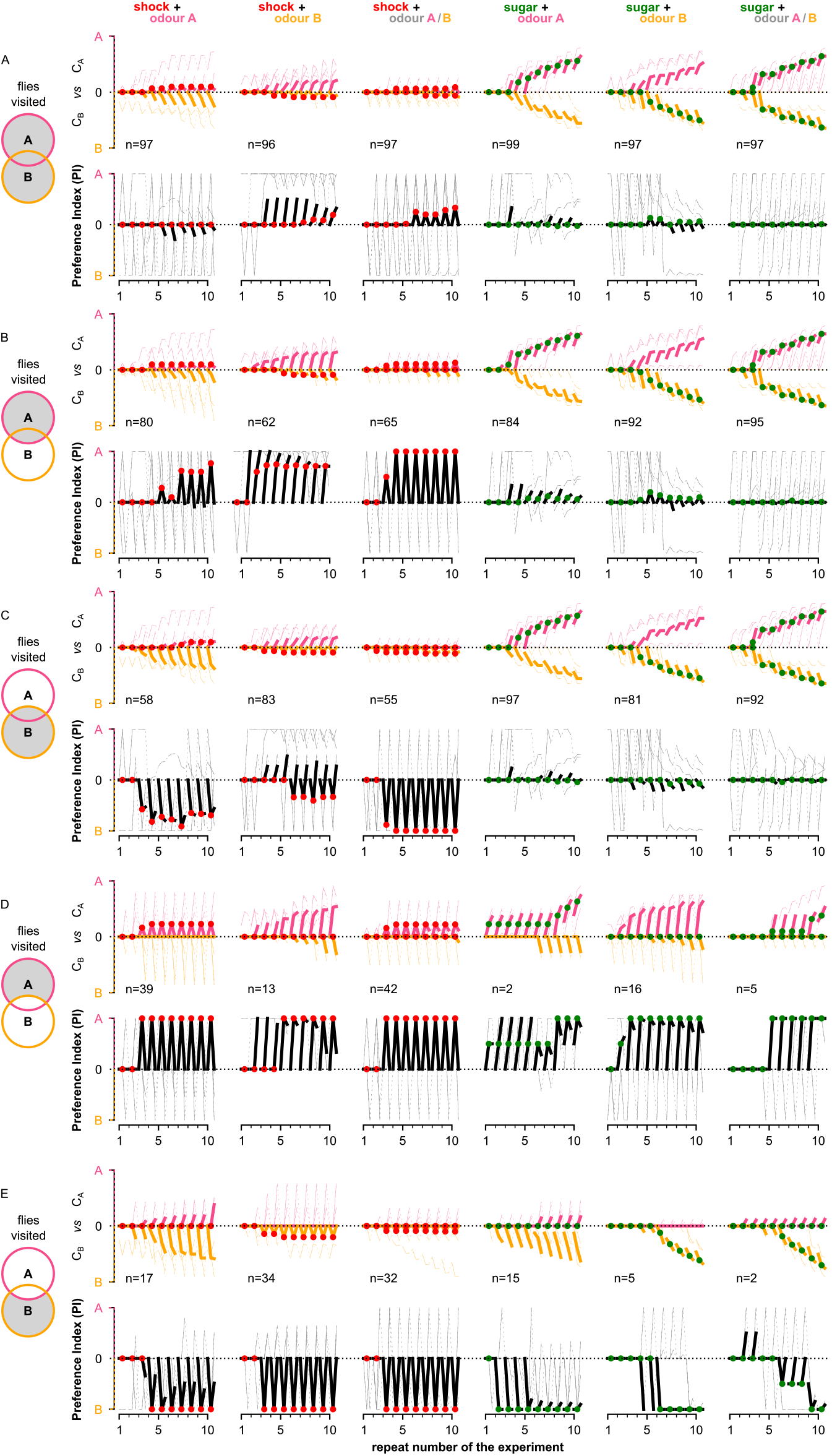
Behavioural summary of simulated flies grouped by the areas that they visited: (A) at least one of the two odours, (B) odour A, (C) odour B, (D) only odour A and (E) only odour B. In each panel, columns show the different conditions and the population for each group. Top row: the normalised cumulative time spend exposed in odour A (pink lines) or odour B (yellow lines - note this line is reversed). For each repeat we present 3 values (average over all the pre-training, training and post-training time-steps respectively) where the values associated with the training phase are marked with red or green dots when punishment or reward has been delivered to that odour respectively. Thin lines show 3 representative samples of individual flies. Thick lines show the median over the simulated flies that visited both odours. Bottom row: the preference index (PI) to each odour extracted by the above cumulative times.

**Figure 11–Figure supplement 3.**
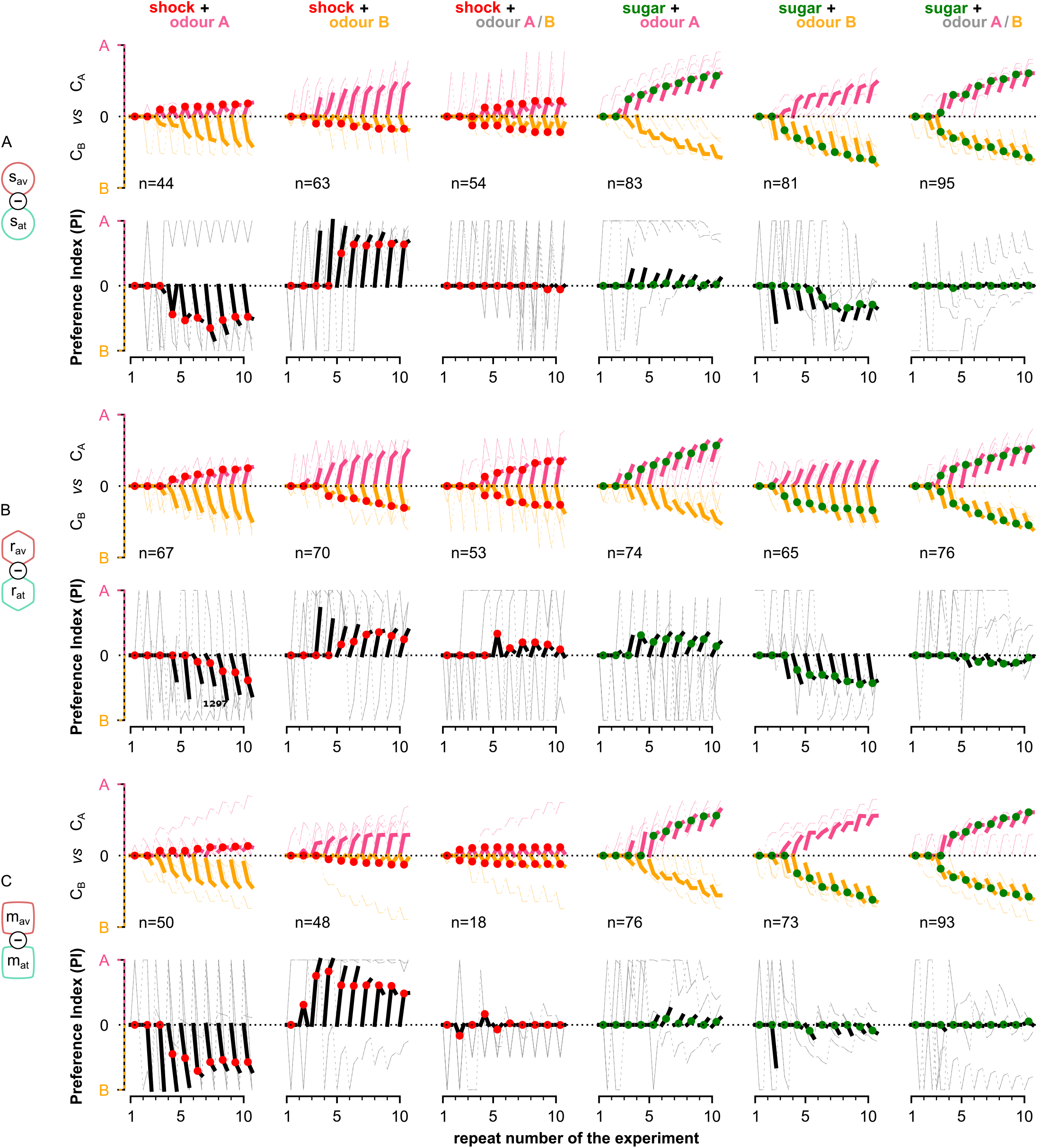
Behavioural summary of simulated flies when controlled by: (A) the susceptible, (B) restrained or (C) LTM MBONs separately. In each panel, columns show the different conditions and the population for each group. Top row: the normalised cumulative time spend exposed in odour A (pink lines) or odour B (yellow lines - note this line is reversed). For each repeat we present 3 values (average over all the pre-training, training and post-training time-steps respectively) where the values associated with the training phase are marked with red or green dots when punishment or reward has been delivered to that odour respectively. Thin lines show 3 representative samples of individual flies. Thick lines show the median over the simulated flies that visited both odours. Bottom row: the preference index (PI) to each odour extracted by the above cumulative times.

**Figure 11–Figure supplement 4.**
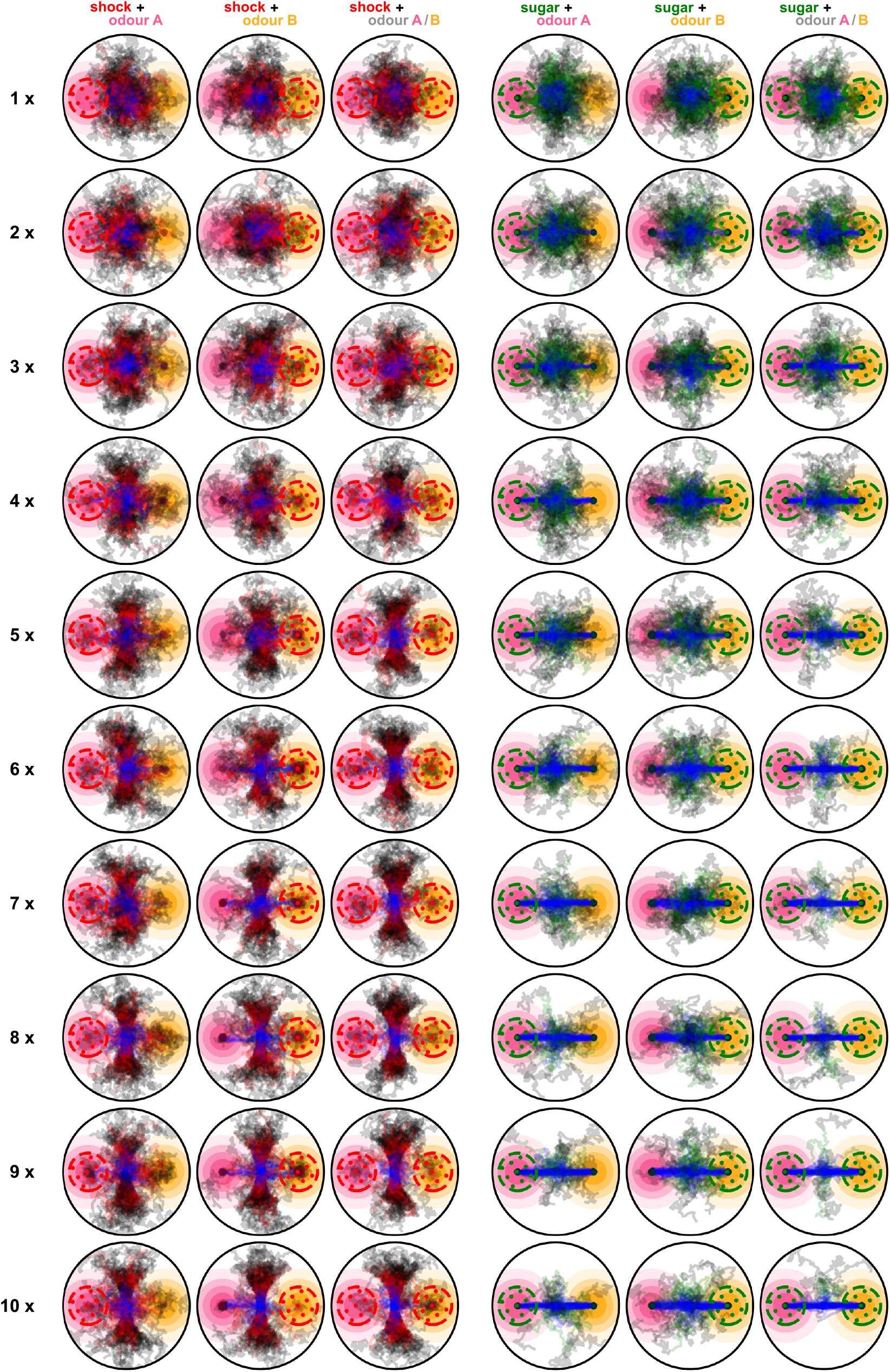
Paths of the flies when using the dopaminergic plasticity rule and during all 10 repeats of the experiment. Blue segments show the paths of the flies during the pre-training, red and green segments show the paths during the training phase for punishment and reward conditions respectively, and black segments show the paths during the post-training phase. Rows are for the different repeats and columns for the different conditions.

**Figure 11–Figure supplement 5.**
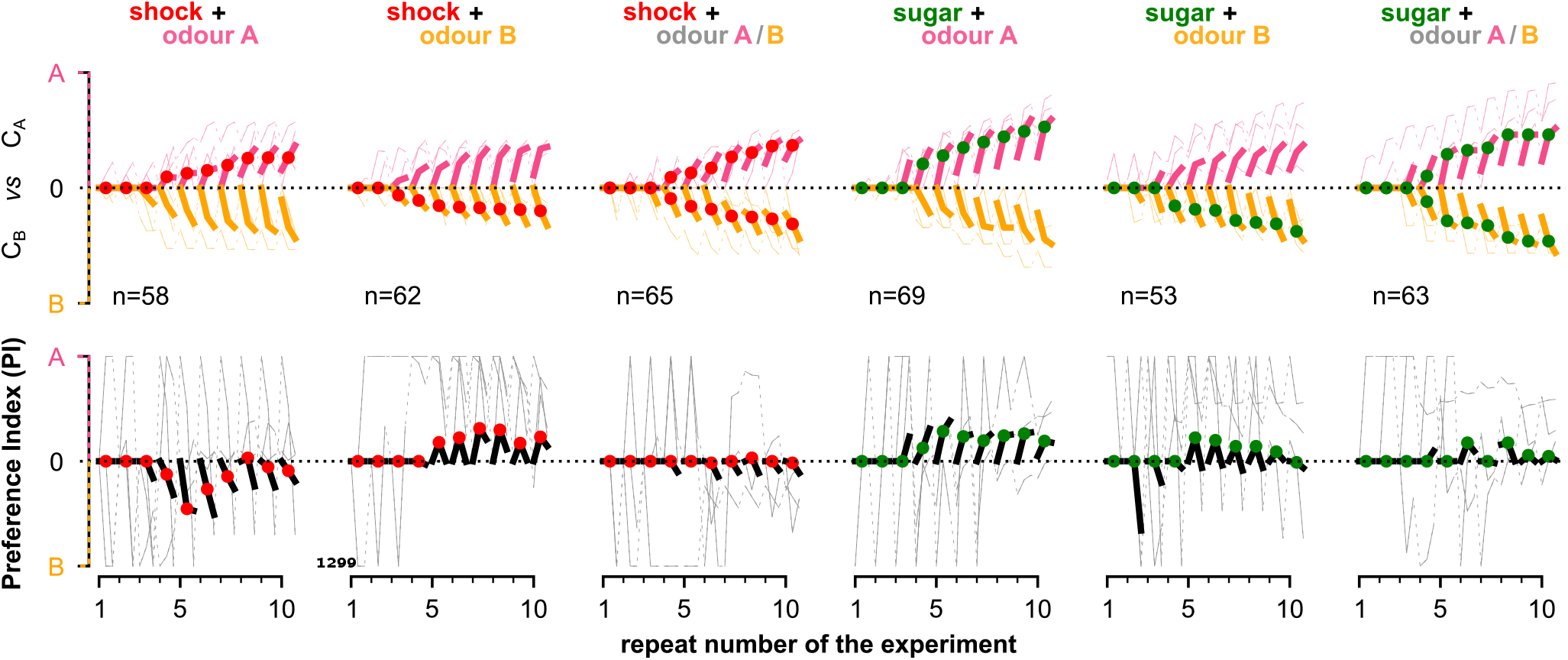
Behavioural summary of a subset of simulated flies that visited both odours at any time during the 10 repeats and the plasticity rule of their neurons was replaced by the reward prediction error plasticity rule. Columns show the different conditions and the population that was recorded visiting both odours. Top row: the normalised cumulative time spend exposed in odour A (pink lines) or odour B (yellow lines - note this line is reversed). For each repeat we present 3 values (average over all the pre-training, training and post-training time-steps respectively) where the values associated with the training phase are marked with red or green dots when punishment or reward has been delivered to that odour respectively. Thin lines show 3 representative samples of individual flies. Thick lines show the median over the simulated flies that visited both odours. Bottom row: the preference index (PI) to each odour extracted by the above cumulative times.

**Figure 11–Figure supplement 6.**
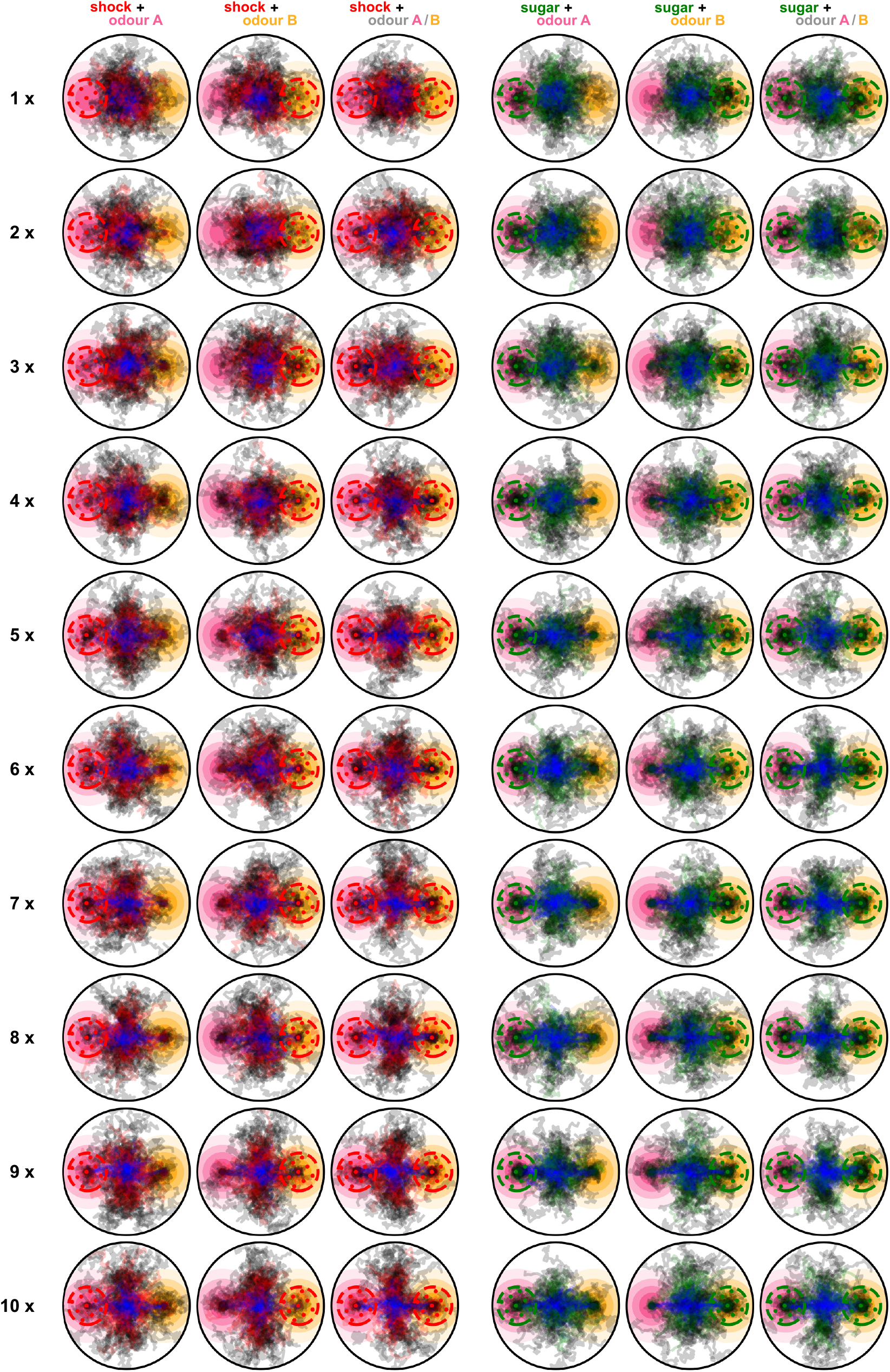
Paths of 100 simulated flies when using the reward prediction error plasticity rule and during all 10 repeats of the experiment. Blue segments show the paths of the flies during the pre-training, red and green segments show the paths during the training phase for punishment and reward conditions respectively, and black segments show the paths during the post-training phase. Rows are for the different repeats and columns for the different conditions.

**Figure 11–Figure supplement 7.**
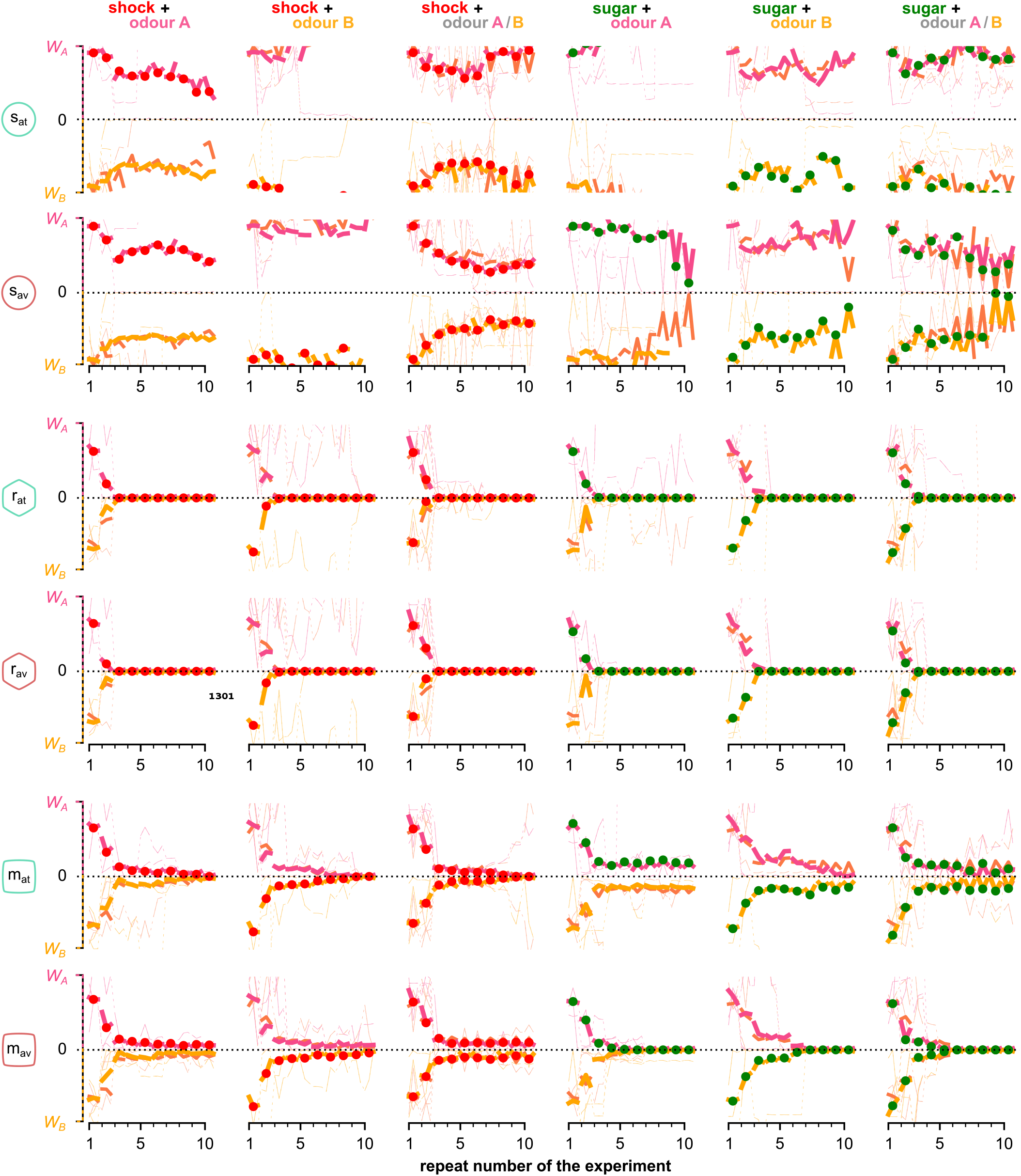
The mean KC→MBON synaptic weights when using the RPE plasticity rule over the simulated flies that visited both odours and for each neuron, phase and repeat of the experiment. Different rows correspond to KC→MBON weights associated with a different type of MBON, from top to bottom: attraction- and avoidance driving susceptible, restrained and long-term memory MBONs. Pink and yellow lines show synaptic weights associated with odour A and B respectively, dashed lines show weights associated with both odours. Thin lines show 3 representative examples of synaptic weights in single simulated flies. Thick lines show the median of the synaptic weight over all the simulated flies that visited both odours.

**Figure 14–Figure supplement 1.**
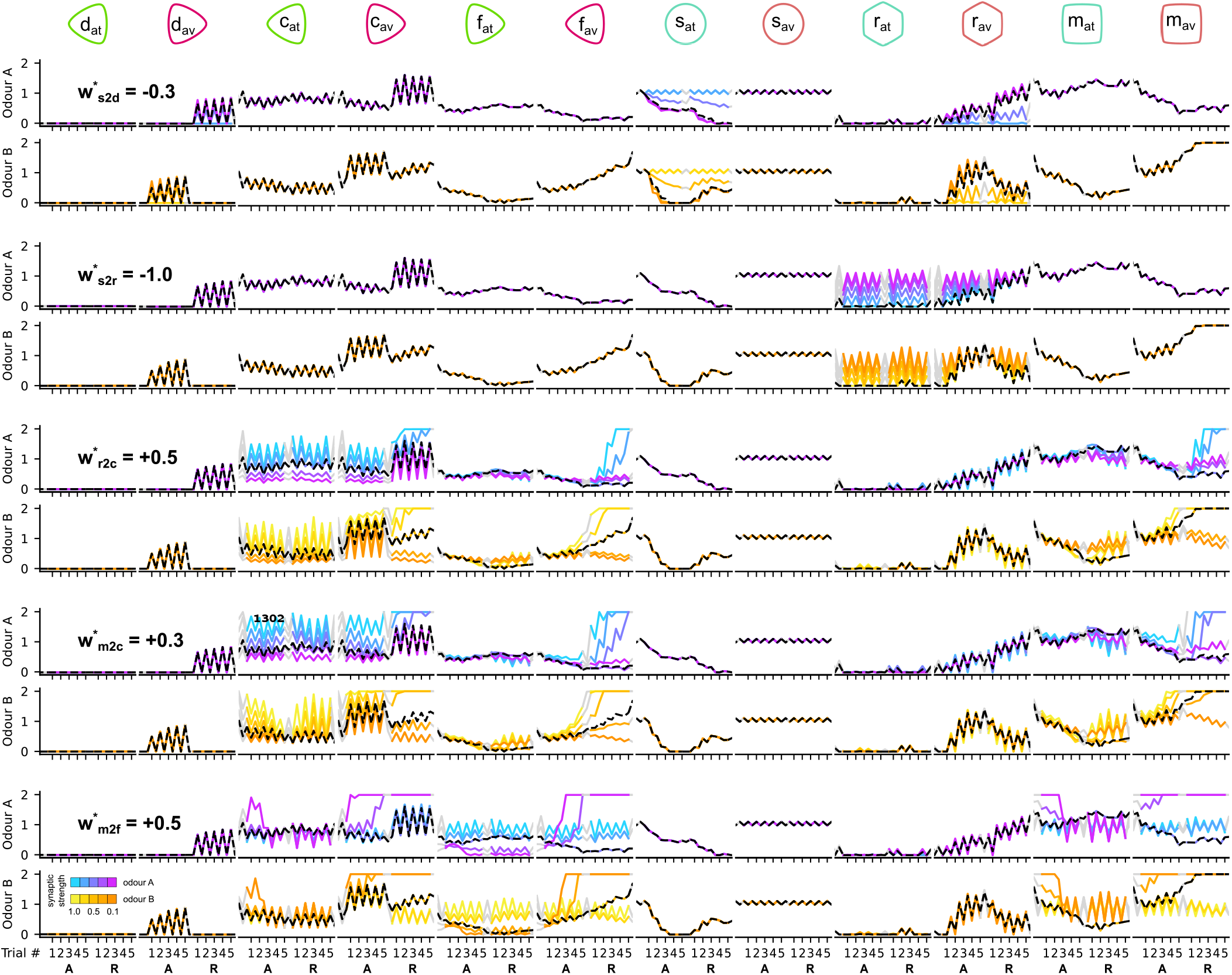
The responses of the DANs and MBONs of the circuit when altering the pre-synaptic strengths of MBONs during the reversal condition. Each column corresponds to the responses of a different neuron. Odd and even rows show the responses of the neurons to odour A and B respectively. Pairs of rows (consecutive odour A and odour B) show the responses of the neurons for the different values of the target parameter (indicated in the first column). Black dashed line shows the responses of the neurons for the chosen (*) parameter. The different colour codes show the responses of the neurons for the different (absolute) values of the target parameter as indicated in the colour-bar on the bottom left.

**Figure 14–Figure supplement 2.**
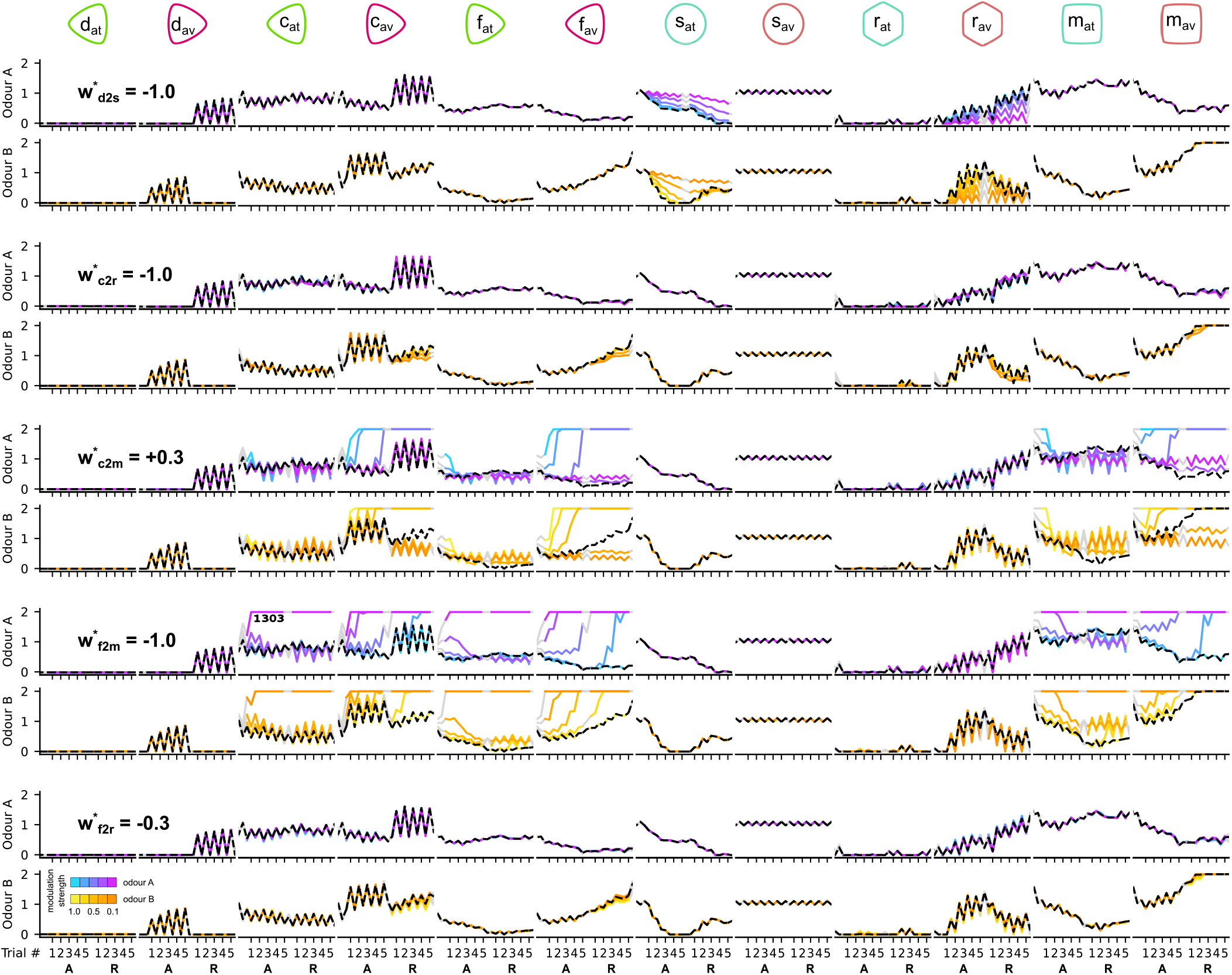
The responses of the DANs and MBONs of the circuit when altering the DA modulation strengths during the reversal condition. Each column corresponds to the responses of a different neuron. Odd and even rows show the responses of the neurons to odour A and B respectively. Pairs of rows (consecutive odour A and odour B) show the responses of the neurons for the different values of the target parameter (indicated in the first column). Black dashed line shows the responses of the neurons for the chosen (*) parameter. The different colour codes show the responses of the neurons for the different values of the target parameter as indicated in the colour-bar on the bottom left.

**Figure 14–Figure supplement 3.**
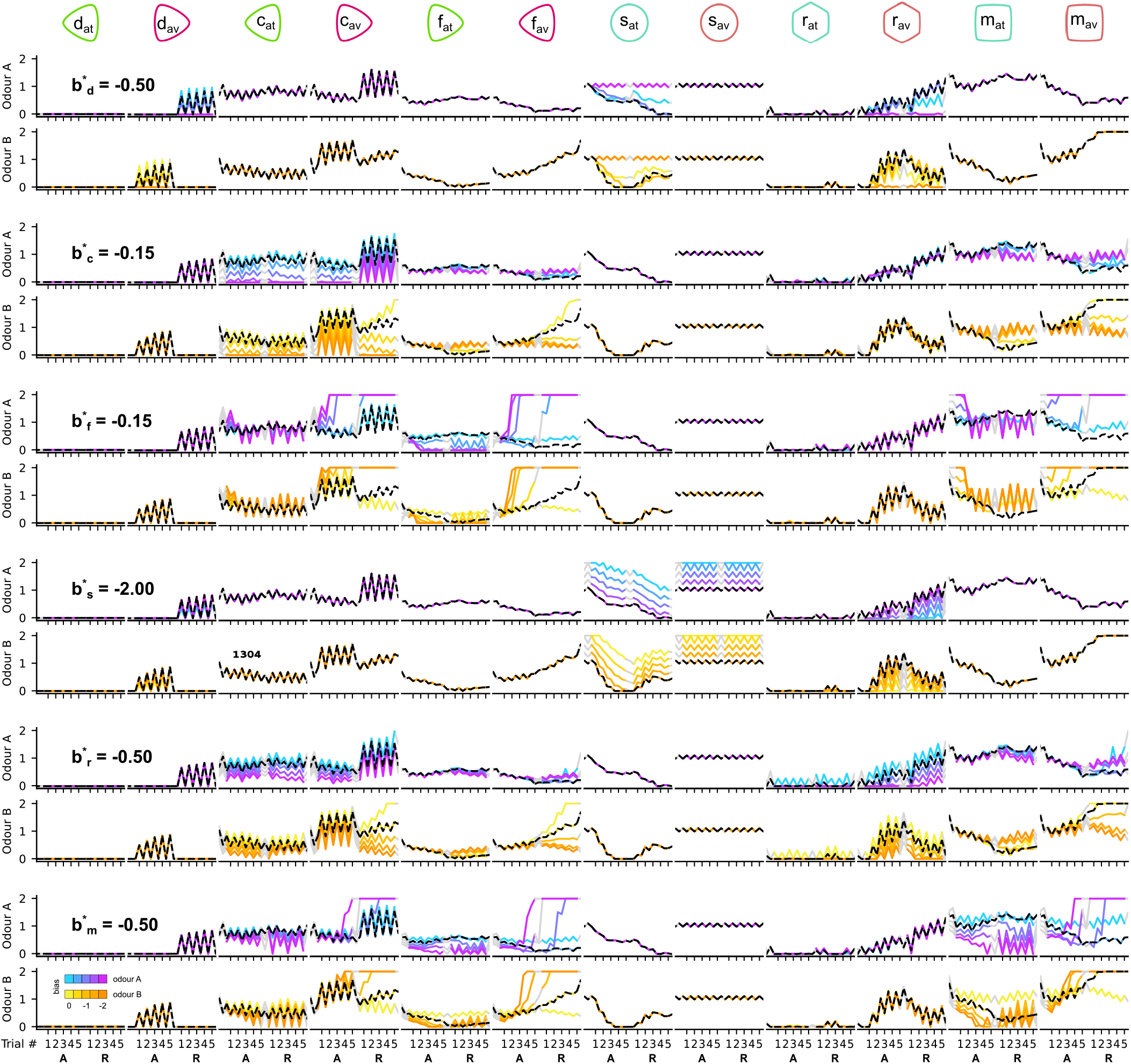
The responses of the DANs and MBONs of the circuit when altering the DAN and MBON biases during the reversal condition. Each column corresponds to the responses of a different neuron. Odd and even rows show the responses of the neurons to odour A and B respectively. Pairs of rows (consecutive odour A and odour B) show the responses of the neurons for the different values of the target parameter (indicated in the first column). Black dashed line shows the responses of the neurons for the chosen (*) parameter. The different colour codes show the responses of the neurons for the different values of the target parameter as indicated in the colour-bar on the bottom left.

**Figure 16–Figure supplement 1.**
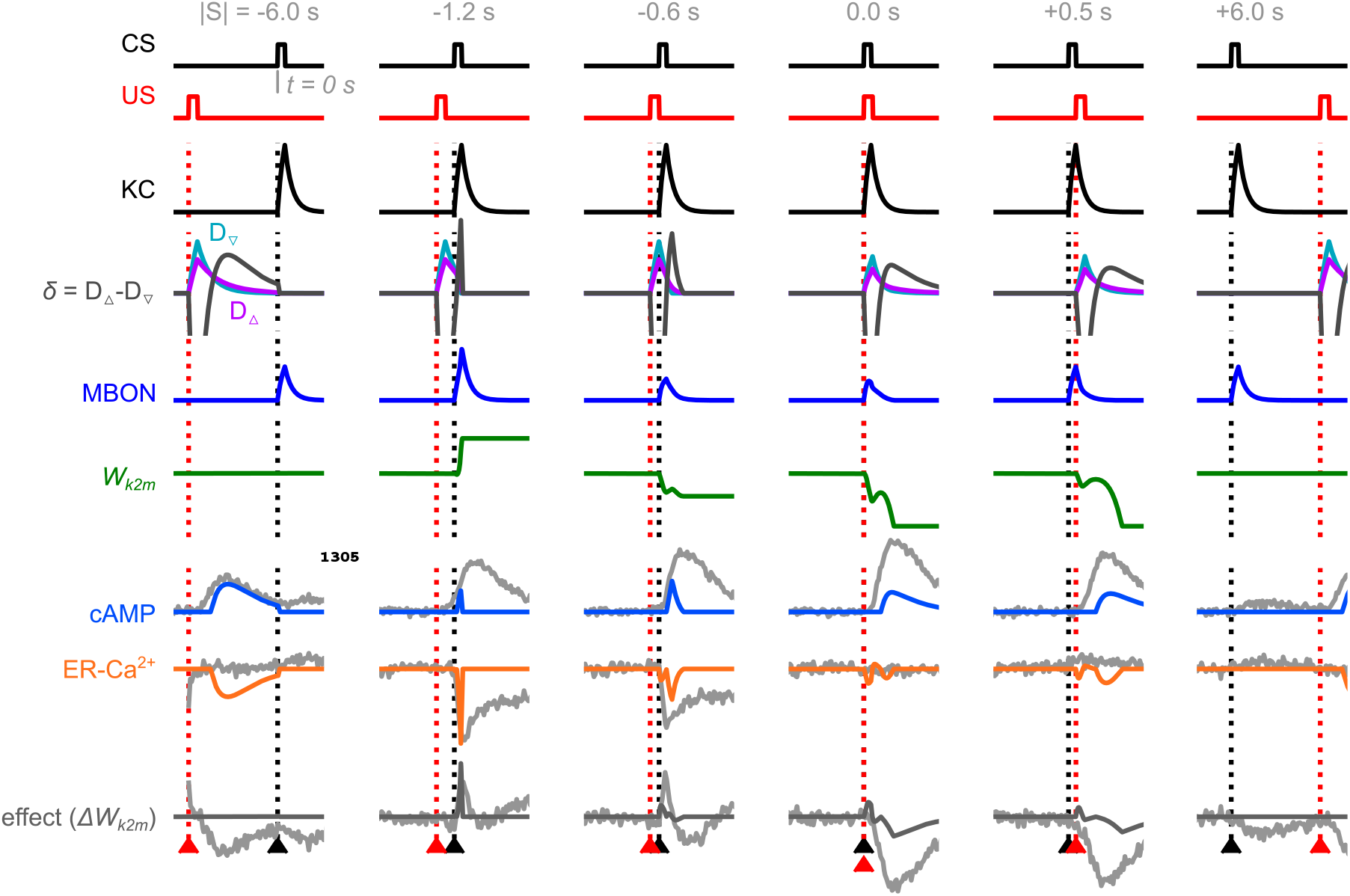
All the chemical levels and neural activities calculated based on the order of the conditional (CS) and unconditional stimuli (US). Black arrowhead marks time of the CS (duration 0.5 sec); red arrowhead marks time of the US (duration 0.6 sec), similar to ***Handler et al.*** (***2019***) - Figure 5D. Predicted ER-Ca^2+^, cAMP and the plasticity effect responses are drawn on top of the data from the original paper (***Handler et al., 2019***) - grey lines.

**Figure 16–Figure supplement 2.**
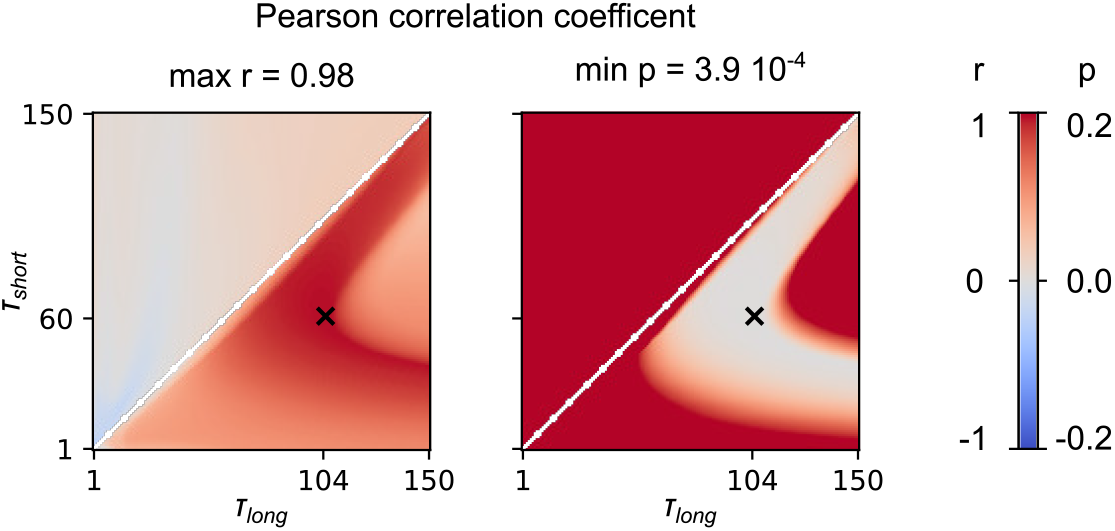
Parameter exploration of the *τ*_short_ and *τ*_long_ in ***Equation 35*** and ***Equation 36***. For the combinations of *τ*_short_ ∈ [1, 150] and *τ*_long_ ∈ [1, 150], we calculate the Pearson correlation coefficient between the predicted normalised mean change of the synaptic weight and the one extracted from the data of ***Handler et al.*** (***2019***), and we report the r and p values. With ‘x’ we mark the pair of parameters with the highest correlation.

## References

Adel M, Griffith LC. The Role of Dopamine in Associative Learning in Drosophila: An Updated Unified Model. Neuroscience Bulletin. 2021; p. 1–22. doi: 10.1007/s12264-021-00665-0.

Ardin P, Peng F, Mangan M, Lagogiannis K, Webb B. Using an Insect Mushroom Body Circuit to Encode Route Memory in Complex Natural Environments. PLOS Computational Biology. 2016; 12(2):e1004683. doi: 10.1371/journal.pcbi.1004683.

Arena P, Patané L, Stornanti V, Termini PS, Zäpf B, Strauss R. Modeling the insect mushroom bodies: Application to a delayed match-to-sample task. Neural Networks. 2013; 41:202–211. doi: 10.1016/j.neunet.2012.11.013.

Aso Y, Hattori D, Yu Y, Johnston RM, Iyer NA, Ngo TT, Dionne H, Abbott L, Axel R, Tanimoto H, Rubin GM. The neuronal architecture of the mushroom body provides a logic for associative learning. eLife. 2014; 3:e04577. doi: 10.7554/elife.04577.

Aso Y, Ray RP, Long X, Bushey D, Cichewicz K, Ngo TT, Sharp B, Christoforou C, Hu A, Lemire AL, Tillberg P, Hirsh J, Litwin-Kumar A, Rubin GM. Nitric oxide acts as a cotransmitter in a subset of dopaminergic neurons to diversify memory dynamics. eLife. 2019; 8:e49257. doi: 10.7554/elife.49257.

Aso Y, Rubin GM. Dopaminergic neurons write and update memories with cell-type-specific rules. eLife. 2016; 5:e16135. doi: 10.7554/elife.16135.

Aso Y, Sitaraman D, Ichinose T, Kaun KR, Vogt K, Belliart-Guérin G, Plaçais PY, Robie AA, Yamagata N, Schnaitmann C, Rowell WJ, Johnston RM, Ngo TTB, Chen N, Korff W, Nitabach MN, Heberlein U, Preat T, Branson KM, Tanimoto H, et al. Mushroom body output neurons encode valence and guide memory-based action selection in Drosophila. eLife. 2014; 3:e04580. doi: 10.7554/elife.04580.

Aso Y, Siwanowicz I, Bräcker L, Ito K, Kitamoto T, Tanimoto H. Specific Dopaminergic Neurons for the Formation of Labile Aversive Memory. Current Biology. 2010; 20(16):1445–1451. doi: 10.1016/j.cub.2010.06.048.

Baddeley B, Graham P, Husbands P, Philippides A. A Model of Ant Route Navigation Driven by Scene Familiarity. PLoS Computational Biology. 2012; 8(1):e1002336. doi: 10.1371/journal.pcbi.1002336.

Balkenius A, Kelber A, Balkenius C. From Animals to Animats 9, 9th International Conference on Simulation of Adaptive Behavior, SAB 2006, Rome, Italy, September 25-29, 2006. Proceedings. Lecture Notes in Computer Science. 2006; p. 422–433. doi: 10.1007/11840541\_35.

Bazhenov M, Huerta R, Smith BH. A Computational Framework for Understanding Decision Making through Integration of Basic Learning Rules. The Journal of Neuroscience. 2013; 33(13):5686–5697. doi: 10.1523/jneurosci.4145-12.2013.

Bell AJ, Sejnowski TJ. An Information-Maximization Approach to Blind Separation and Blind Deconvolution. Neural Computation. 1995; 7(6):1129–1159. doi: 10.1162/neco.1995.7.6.1129.

Bennett JEM, Philippides A, Nowotny T. Learning with reinforcement prediction errors in a model of the Drosophila mushroom body. Nature Communications. 2021; 12(1):2569. doi: 10.1038/s41467-021-22592-4.

Berry JA, Phan A, Davis RL. Dopamine Neurons Mediate Learning and Forgetting through Bidirectional Modulation of a Memory Trace. Cell Reports. 2018; 25(3):651–662.e5. doi: 10.1016/j.celrep.2018.09.051.

Bilz F, Geurten BRH, Hancock CE, Widmann A, Fiala A. Visualization of a Distributed Synaptic Memory Code in the Drosophila Brain. Neuron. 2020; 106(6):963–976.e4. doi: 10.1016/j.neuron.2020.03.010.

Brembs B, Heisenberg M. Conditioning with compound stimuli in Drosophila melanogaster in the flight simulator. The Journal of experimental biology. 2001; 204(Pt 16):2849–59.

Burke CJ, Huetteroth W, Owald D, Perisse E, Krashes MJ, Das G, Gohl D, Silies M, Certel S, Waddell S. Layered reward signalling through octopamine and dopamine in Drosophila. Nature. 2012; 492(7429):433–437. doi: 10.1038/nature11614.

Busto GU, Cervantes-Sandoval I, Davis RL. Olfactory Learning in Drosophila. Physiology. 2010; 25(6):338–346. doi: 10.1152/physiol.00026.2010.

Campbell RAA, Honegger KS, Qin H, Li W, Demir E, Turner GC. Imaging a Population Code for Odor Identity in the Drosophila Mushroom Body. The Journal of Neuroscience. 2013; 33(25):10568–10581. doi: 10.1523/jneurosci.0682-12.2013.

Cervantes-Sandoval I, Phan A, Chakraborty M, Davis RL. Reciprocal synapses between mushroom body and dopamine neurons form a positive feedback loop required for learning. eLife. 2017; 6:e23789. doi: 10.7554/elife.23789.

Claridge-Chang A, Roorda RD, Vrontou E, Sjulson L, Li H, Hirsh J, Miesenböck G. Writing Memories with Light-Addressable Reinforcement Circuitry. Cell. 2009; 139(2):405–415. doi: 10.1016/j.cell.2009.08.034.

Cleland TA. Inhibitory glutamate receptor channels. Molecular Neurobiology. 1996; 13(2):97–136. doi: 10.1007/bf02740637.

Cohn R, Morantte I, Ruta V. Coordinated and Compartmentalized Neuromodulation Shapes Sensory Processing in Drosophila. Cell. 2015; 163(7):1742–1755. doi: 10.1016/j.cell.2015.11.019.

Colomb J, Kaiser L, Chabaud MA, Preat T. Parametric and genetic analysis of Drosophila appetitive long-term memory and sugar motivation. Genes, Brain and Behavior. 2009; 8(4):407–415. doi: 10.1111/j.1601-183x.2009.00482.x.

Dalgleish T. The emotional brain. Nature Reviews Neuroscience. 2004; 5(7):583–589. doi: 10.1038/nrn1432.

Davis RL. Mushroom bodies and drosophila learning. Neuron. 1993; 11(1):1–14. doi: 10.1016/0896-6273(93)90266-t.

Delahunt CB, Riffell JA, Kutz JN. Biological Mechanisms for Learning: A Computational Model of Olfactory Learning in the Manduca sexta Moth, With Applications to Neural Nets. Frontiers in Computational Neuroscience. 2018; 12:102. doi: 10.3389/fncom.2018.00102.

Dubnau J, Grady L, Kitamoto T, Tully T. Disruption of neurotransmission in Drosophila mushroom body blocks retrieval but not acquisition of memory. Nature. 2001; 411(6836):476–480. doi: 10.1038/35078077.

Dylla KV, Raiser G, Galizia CG, Szyszka P. Trace Conditioning in Drosophila Induces Associative Plasticity in Mushroom Body Kenyon Cells and Dopaminergic Neurons. Frontiers in Neural Circuits. 2017; 11:42. doi: 10.3389/fncir.2017.00042.

Eichler K, Li F, Litwin-Kumar A, Park Y, Andrade I, Schneider-Mizell CM, Saumweber T, Huser A, Eschbach C, Gerber B, Fetter RD, Truman JW, Priebe CE, Abbott LF, Thum AS, Zlatic M, Cardona A. The complete connectome of a learning and memory centre in an insect brain. Nature. 2017 8; 548(7666):175–182. doi: 10.1038/nature23455.

Eschbach C, Fushiki A, Winding M, Schneider-Mizell CM, Shao M, Arruda R, Eichler K, Valdes-Aleman J, Ohyama T, Thum AS, Gerber B, Fetter RD, Truman JW, Litwin-Kumar A, Cardona A, Zlatic M. Recurrent architecture for adaptive regulation of learning in the insect brain. Nature Neuroscience. 2020; 23(4):544–555. doi: 10.1038/s41593-020-0607-9.

Faghihi F, Moustafa AA, Heinrich R, Wörgötter F. A computational model of conditioning inspired by Drosophila olfactory system. Neural Networks. 2017; 87:96–108. doi: 10.1016/j.neunet.2016.11.002.

Felsenberg J, Barnstedt O, Cognigni P, Lin S, Waddell S. Re-evaluation of learned information in Drosophila. Nature. 2017; 544(7649):240–244. doi: 10.1038/nature21716.

Felsenberg J, Jacob PF, Walker T, Barnstedt O, Edmondson-Stait AJ, Pleijzier MW, Otto N, Schlegel P, Sharifi N, Perisse E, Smith CS, Lauritzen JS, Costa M, Jefferis GSXE, Bock DD, Waddell S. Integration of Parallel Opposing Memories Underlies Memory Extinction. Cell. 2018; 175(3):709–722.e15. doi: 10.1016/j.cell.2018.08.021.

Finelli LA, Haney S, Bazhenov M, Stopfer M, Sejnowski TJ. Synaptic Learning Rules and Sparse Coding in a Model Sensory System. PLoS Computational Biology. 2008; 4(4):e1000062. doi: 10.1371/journal.pcbi.1000062.

Gerber B, Stocker RF, Tanimura T, Thum AS. Smelling, Tasting, Learning: Drosophila as a Study Case. In: Meyerhof W, Korsching S, editors. Chemosensory Systems in Mammals, Fishes, and Insects, vol. 47 Springer; 2009. p. 139–186. doi: 10.1007/400\_2008\_9.

Gerber B, Hendel T. Outcome expectations drive learned behaviour in larval Drosophila. Proceedings of the Royal Society B: Biological Sciences. 2006; 273(1604):2965–2968. doi: 10.1098/rspb.2006.3673.

Handler A, Graham TGW, Cohn R, Morantte I, Siliciano AF, Zeng J, Li Y, Ruta V. Distinct Dopamine Receptor Pathways Underlie the Temporal Sensitivity of Associative Learning. Cell. 2019; 178(1):60–75.e19. doi: 10.1016/j.cell.2019.05.040.

Hebb DO. The organization of behavior: A neuropsychological theory. Psychology Press; 2005.

Heisenberg M. Mushroom body memoir: from maps to models. Nature Reviews Neuroscience. 2003; 4(4):266–275. doi: 10.1038/nrn1074.

Hige T, Aso Y, Modi M, Rubin G, Turner G. Heterosynaptic Plasticity Underlies Aversive Olfactory Learning in Drosophila. Neuron. 2015; 88(5):985–998. doi: 10.1016/j.neuron.2015.11.003.

Huerta R, Huerta R, Nowotny T, Nowotny T, García-Sánchez M, Abarbanel HDI, Rabinovich MI. Learning Classification in the Olfactory System of Insects. Neural Computation. 2004; 16(8):1601–1640. doi: 10.1162/089976604774201613.

Huetteroth W, Perisse E, Lin S, Klappenbach M, Burke C, Waddell S. Sweet Taste and Nutrient Value Subdivide Rewarding Dopaminergic Neurons in Drosophila. Current Biology. 2015; 25(6):751–758. doi: 10.1016/j.cub.2015.01.036.

Ichinose T, Aso Y, Yamagata N, Abe A, Rubin GM, Tanimoto H. Reward signal in a recurrent circuit drives appetitive long-term memory formation. eLife. 2015; 4:e10719. doi: 10.7554/elife.10719.

Ito I, Ong RCy, Raman B, Stopfer M. Sparse odor representation and olfactory learning. Nature Neuroscience. 2008; 11(10):1177–1184. doi: 10.1038/nn.2192.

Jacob PF, Waddell S. Spaced Training Forms Complementary Long-Term Memories of Opposite Valence in Drosophila. Neuron. 2020; 106(6):977–991.e4. doi: 10.1016/j.neuron.2020.03.013.

Kallman BR, Kim H, Scott K. Excitation and inhibition onto central courtship neurons biases Drosophila mate choice. eLife. 2015; 4:e11188. doi: 10.7554/elife.11188.

Kamin LJ. Predictability, surprise, attention and conditioning. Punishment and Aversive Behaviour. 1967; p. 279–296.

Krashes MJ, Waddell S. Drosophila Aversive Olfactory Conditioning. Cold Spring Harbor Protocols. 2011; 2011(5):pdb.prot5608–pdb.prot5608. doi: 10.1101/pdb.prot5608.

Krashes MJ, DasGupta S, Vreede A, White B, Armstrong JD, Waddell S. A Neural Circuit Mechanism Integrating Motivational State with Memory Expression in Drosophila. Cell. 2009; 139(2):416–427. doi: 10.1016/j.cell.2009.08.035.

Krashes MJ, Keene AC, Leung B, Armstrong JD, Waddell S. Sequential Use of Mushroom Body Neuron Subsets during Drosophila Odor Memory Processing. Neuron. 2007; 53(1):103–115. doi: 10.1016/j.neuron.2006.11.021.

Krashes MJ, Waddell S. Drosophila Appetitive Olfactory Conditioning. Cold Spring Harbor Protocols. 2011; 2011(5):pdb.prot5609. doi: 10.1101/pdb.prot5609.

Lee TW, Girolami M, Sejnowski TJ. Independent Component Analysis Using an Extended Infomax Algorithm for Mixed Subgaussian and Supergaussian Sources. Neural Computation. 1999; 11(2):417–441. doi: 10.1162/089976699300016719.

Li F, Lindsey JW, Marin EC, Otto N, Dreher M, Dempsey G, Stark I, Bates AS, Pleijzier MW, Schlegel P, Nern A, Takemura Sy, Eckstein N, Yang T, Francis A, Braun A, Parekh R, Costa M, Scheffer LK, Aso Y, et al. The connectome of the adult Drosophila mushroom body provides insights into function. eLife. 2020; 9:e62576. doi: 10.7554/elife.62576.

Lin S, Owald D, Chandra V, Talbot C, Huetteroth W, Waddell S. Neural correlates of water reward in thirsty Drosophila. Nature Neuroscience. 2014; 17(11):1536–1542. doi: 10.1038/nn.3827.

Liu C, Plaçais PY, Yamagata N, Pfeiffer BD, Aso Y, Friedrich AB, Siwanowicz I, Rubin GM, Preat T, Tanimoto H. A subset of dopamine neurons signals reward for odour memory in Drosophila. Nature. 2012; 488(7412):512–516. doi: 10.1038/nature11304.

Liu WW, Wilson RI. Glutamate is an inhibitory neurotransmitter in the Drosophila olfactory system. Proceedings of the National Academy of Sciences. 2013; 110(25):10294–10299. doi: 10.1073/pnas.1220560110.

Liu X, Davis RL. The GABAergic anterior paired lateral neuron suppresses and is suppressed by olfactory learning. Nature Neuroscience. 2009; 12(1):53–59. doi: 10.1038/nn.2235.

Lulham A, Bogacz R, Vogt S, Brown MW. An Infomax Algorithm Can Perform Both Familiarity Discrimination and Feature Extraction in a Single Network. Neural Computation. 2011; 23(4):909–926. doi: 10.1162/neco\_a\_00097.

MacLean PD. Psychosomatic Disease and the “Visceral Brain”. Psychosomatic Medicine. 1949; 11(6):338–353. doi: 10.1097/00006842-194911000-00003.

Mao Z, Davis RL. Eight Different Types of Dopaminergic Neurons Innervate the Drosophila Mushroom Body Neuropil: Anatomical and Physiological Heterogeneity. Frontiers in Neural Circuits. 2009; 3:5. doi: 10.3389/neuro.04.005.2009.

May CE, Rosander J, Gottfried J, Dennis E, Dus M. Dietary sugar inhibits satiation by decreasing the central processing of sweet taste. eLife. 2020; 9:e54530. doi: 10.7554/elife.54530.

McCarthy Ev, Wu Y, deCarvalho T, Brandt C, Cao G, Nitabach MN. Synchronized Bilateral Synaptic Inputs to Drosophila melanogaster Neuropeptidergic Rest/Arousal Neurons. The Journal of Neuroscience. 2011; 31(22):8181–8193. doi: 10.1523/jneurosci.2017-10.2011.

McCurdy LY, Sareen P, Davoudian PA, Nitabach MN. Dopaminergic mechanism underlying reward-encoding of punishment omission during reversal learning in Drosophila. Nature Communications. 2021 2; 12(1):1115. doi: 10.1038/s41467-021-21388-w.

McGuire SE, Le PT, Davis RL. The Role of Drosophila Mushroom Body Signaling in Olfactory Memory. Science. 2001; 293(5533):1330–1333. doi: 10.1126/science.1062622.

Michels B, Chen Yc, Saumweber T, Mishra D, Tanimoto H, Schmid B, Engmann O, Gerber B. Cellular site and molecular mode of synapsin action in associative learning. Learning & Memory. 2011; 18(5):332–344. doi: 10.1101/lm.2101411.

Milyaev N, Osumi-Sutherland D, Reeve S, Burton N, Baldock RA, Armstrong JD. The Virtual Fly Brain browser and query interface. Bioinformatics. 2012; 28(3):411–415. doi: 10.1093/bioinformatics/btr677.

Niv Y. Reinforcement learning in the brain. Journal of Mathematical Psychology. 2009; 53(3):139–154. doi: 10.1016/j.jmp.2008.12.005.

Owald D, Felsenberg J, Talbot CB, Das G, Perisse E, Huetteroth W, Waddell S. Activity of Defined Mushroom Body Output Neurons Underlies Learned Olfactory Behavior in Drosophila. Neuron. 2015; 86(2):417–427. doi: 10.1016/j.neuron.2015.03.025.

Papez JW. A proposed mechanism of emotion. Archives of Neurology & Psychiatry. 1937; 38(4):725–743. doi: 10.1001/archneurpsyc.1937.02260220069003.

Pavlowsky A, Schor J, Plaçais PY, Preat T. A GABAergic Feedback Shapes Dopaminergic Input on the Drosophila Mushroom Body to Promote Appetitive Long-Term Memory. Current Biology. 2018; 28(11):1783–1793.e4. doi: 10.1016/j.cub.2018.04.040.

Peng F, Chittka L. A Simple Computational Model of the Bee Mushroom Body Can Explain Seemingly Complex Forms of Olfactory Learning and Memory. Current Biology. 2017; 27(2):224–230. doi: 10.1016/j.cub.2016.10.054.

Perisse E, Owald D, Barnstedt O, Talbot C, Huetteroth W, Waddell S. Aversive Learning and Appetitive Motivation Toggle Feed-Forward Inhibition in the Drosophila Mushroom Body. Neuron. 2016; 90(5):1086–1099. doi: 10.1016/j.neuron.2016.04.034.

Plaçais PY, Trannoy S, Friedrich A, Tanimoto H, Preat T. Two Pairs of Mushroom Body Efferent Neurons Are Required for Appetitive Long-Term Memory Retrieval in Drosophila. Cell Reports. 2013; 5(3):769–780. doi: 10.1016/j.celrep.2013.09.032.

Plutchik R. The Nature of Emotions. American Scientist. 2001; 89(4):344–350.

Pribbenow C, Chen Yc, Heim MM, Laber D, Reubold S, Reynolds E, Balles I, Grimalt RS, Rauch C, Rösner J, Alquicira TFdV, Owald D. Postsynaptic plasticity of cholinergic synapses underlies the induction and expression of appetitive memories in Drosophila. bioRxiv. 2021; p. 2021.07.01.450776. doi: 10.1101/2021.07.01.450776.

Rescorla RA, Wagner AR. A theory of Pavlovian conditioning: Variations in the effectiveness of reinforcement and nonreinforcement. In: Classical conditioning II: current research and theory New York: Appleton-Century-Crofts; 1972. p. 64–99.

Roxo MR, Franceschini PR, Zubaran C, Kleber FD, Sander JW. The Limbic System Conception and Its Historical Evolution. TheScientificWorldJOURNAL. 2011; 11:2427–2440. doi: 10.1100/2011/157150.

Saumweber T, Rohwedder A, Schleyer M, Eichler K, Chen Yc, Aso Y, Cardona A, Eschbach C, Kobler O, Voigt A, Durairaja A, Mancini N, Zlatic M, Truman JW, Thum AS, Gerber B. Functional architecture of reward learning in mushroom body extrinsic neurons of larval Drosophila. Nature Communications. 2018; 9(1):1104. doi: 10.1038/s41467-018-03130-1.

Schleyer M, Fendt M, Schuller S, Gerber B. Associative Learning of Stimuli Paired and Unpaired With Reinforcement: Evaluating Evidence From Maggots, Flies, Bees, and Rats. Frontiers in Psychology. 2018; 9:1494. doi: 10.3389/fpsyg.2018.01494.

Schleyer M, Saumweber T, Nahrendorf W, Fischer B, Alpen Dv, Pauls D, Thum A, Gerber B. A behavior-based circuit model of how outcome expectations organize learned behavior in larval Drosophila. Learning & Memory. 2011; 18(10):639–653. doi: 10.1101/lm.2163411.

Schleyer M, Weiglein A, Thoener J, Strauch M, Hartenstein V, Weigelt MK, Schuller S, Saumweber T, Eichler K, Rohwedder A, Merhof D, Zlatic M, Thum AS, Gerber B. Identification of dopaminergic neurons that can both establish associative memory and acutely terminate its behavioral expression. Journal of Neuroscience. 2020; 40(31):JN-RM–0290-20. doi: 10.1523/jneurosci.0290-20.2020.

Schroll C, Riemensperger T, Bucher D, Ehmer J, Völler T, Erbguth K, Gerber B, Hendel T, Nagel G, Buchner E, Fiala A. Light-Induced Activation of Distinct Modulatory Neurons Triggers Appetitive or Aversive Learning in Drosophila Larvae. Current Biology. 2006; 16(17):1741–1747. doi: 10.1016/j.cub.2006.07.023.

Schwaerzel M, Monastirioti M, Scholz H, Friggi-Grelin F, Birman S, Heisenberg M. Dopamine and Octopamine Differentiate between Aversive and Appetitive Olfactory Memories in Drosophila. Journal of Neuroscience. 2003; 23(33):10495–10502. doi: 10.1523/jneurosci.23-33-10495.2003.

Senapati B, Tsao CH, Juan YA, Chiu TH, Wu CL, Waddell S, Lin S. A neural mechanism for deprivation state-specific expression of relevant memories in Drosophila. Nature Neuroscience. 2019; 22(12):2029–2039. doi: 10.1038/s41593-019-0515-z.

Smith D, Wessnitzer J, Webb B. A model of associative learning in the mushroom body. Biological Cybernetics. 2008; 99(2):89–103. doi: 10.1007/s00422-008-0241-1.

Springer M, Nawrot MP. A mechanistic model for reward prediction and extinction learning in the fruit fly. eNeuro. 2021; p. ENEURO.0549–20.2021. doi: 10.1523/eneuro.0549-20.2021.

Sten TH, Li R, Otopalik A, Ruta V. An arousal-gated visual circuit controls pursuit during Drosophila courtship. bioRxiv. 2020; p. 2020.08.31.275883. doi: 10.1101/2020.08.31.275883.

Tabone CJ, Belle JSd. Second-order conditioning in Drosophila. Learning & Memory. 2011; 18(4):250–253. doi: 10.1101/lm.2035411.

Takemura Sy, Aso Y, Hige T, Wong A, Lu Z, Xu CS, Rivlin PK, Hess H, Zhao T, Parag T, Berg S, Huang G, Katz W, Olbris DJ, Plaza S, Umayam L, Aniceto R, Chang LA, Lauchie S, Ogundeyi O, et al. A connectome of a learning and memory center in the adult Drosophila brain. eLife. 2017; 6:e26975. doi: 10.7554/elife.26975.

Tanaka NK, Tanimoto H, Ito K. Neuronal assemblies of the Drosophila mushroom body. Journal of Comparative Neurology. 2008; 508(5):711–755. doi: 10.1002/cne.21692.

Turner GC, Bazhenov M, Laurent G. Olfactory Representations by Drosophila Mushroom Body Neurons. Journal of Neurophysiology. 2008; 99(2):734–746. doi: 10.1152/jn.01283.2007.

Waddell S. Dopamine reveals neural circuit mechanisms of fly memory. Trends in Neurosciences. 2010; 33(10):457–464. doi: 10.1016/j.tins.2010.07.001.

Wessnitzer J, Young JM, Armstrong JD, Webb B. A model of non-elemental olfactory learning in Drosophila. Journal of Computational Neuroscience. 2012; 32(2):197–212. doi: 10.1007/s10827-011-0348-6.

Wu CL, Shih MF, Lee PT, Chiang AS. An Octopamine-Mushroom Body Circuit Modulates the Formation of Anesthesia-Resistant Memory in Drosophila. Current Biology. 2013; 23(23):2346–2354. doi: 10.1016/j.cub.2013.09.056.

Wu Y, Ren Q, Li H, Guo A. The GABAergic anterior paired lateral neurons facilitate olfactory reversal learning in Drosophila. Learning & Memory. 2012; 19(10). doi: 10.1101/lm.025726.112.

Wu Z, Guo A. A model study on the circuit mechanism underlying decision-making in Drosophila. Neural Networks. 2011; 24(4):333–344. doi: 10.1016/j.neunet.2011.01.002.

Yamagata N, Hiroi M, Kondo S, Abe A, Tanimoto H. Suppression of Dopamine Neurons Mediates Reward. PLoS Biology. 2016; 14(12):e1002586. doi: 10.1371/journal.pbio.1002586.

Young JM, Wessnitzer J, Armstrong JD, Webb B. Elemental and non-elemental olfactory learning in Drosophila. Neurobiology of Learning and Memory. 2011; 96(2):339–352. doi: 10.1016/j.nlm.2011.06.009.

Zhang YV, Ni J, Montell C. The Molecular Basis for Attractive Salt-Taste Coding in Drosophila. Science. 2013; 340(6138):1334–1338. doi: 10.1126/science.1234133.

Zhao C, Widmer YF, Diegelmann S, Petrovici MA, Sprecher SG, Senn W. Predictive olfactory learning in Drosophila. Scientific Reports. 2021; 11(1):6795. doi: 10.1038/s41598-021-85841-y.

Zhao F, Zeng Y, Guo A, Su H, Xu B. A neural algorithm for Drosophila linear and nonlinear decision-making. Scientific Reports. 2020; 10(1):18660. doi: 10.1038/s41598-020-75628-y.

Zhou M, Chen N, Tian J, Zeng J, Zhang Y, Zhang X, Guo J, Sun J, Li Y, Guo A, Li Y. Suppression of GABAergic neurons through D2-like receptor secures efficient conditioning in Drosophila aversive olfactory learning. Proceedings of the National Academy of Sciences. 2019; 116(11):201812342. doi: 10.1073/pnas.1812342116.

Zhu L, Mangan M, Webb B. Biomimetic and Biohybrid Systems, 9th International Conference, Living Machines 2020, Freiburg, Germany, July 28–30, 2020, Proceedings. Lecture Notes in Computer Science. 2020; p. 415–426. doi: 10.1007/978-3-030-64313-3\_39.

